# Markovian Weighted Ensemble Milestoning (M-WEM): Long-time Kinetics from Short Trajectories

**DOI:** 10.1101/2021.06.26.450057

**Authors:** Dhiman Ray, Sharon Emily Stone, Ioan Andricioaei

## Abstract

We introduce a rare-event sampling scheme, named Markovian Weighted Ensemble Milestoning (M-WEM), which inlays a weighted ensemble framework within a Markovian milestoning theory to efficiently calculate thermodynamic and kinetic properties of long-timescale biomolecular processes from short atomistic molecular dynamics simulations. M-WEM is tested on the Müller-Brown potential model, the conformational switching in alanine dipeptide, and the millisecond timescale protein-ligand unbinding in a trypsin-benzamidine complex. Not only can M-WEM predict the kinetics of these processes with quantitative accuracy, but it also allows for a scheme to reconstruct a multidimensional free energy landscape along additional degrees of freedom which are *not* part of the milestoning progress coordinate. For the ligand-receptor system, the experimental residence time, association and dissociation kinetics, and binding free energy could be reproduced using M-WEM within a simulation time of a few hundreds of nanoseconds, which is a fraction of the computational cost of other currently available methods, and close to four orders of magnitude less than the experimental residence time. Due to the high accuracy and low computational cost, the M-WEM approach can find potential application in kinetics and free-energy based computational drug design.

## Introduction

It is a challenge to quantify with accuracy the kinetics of rare events in molecular biophysics via computational means. Molecular dynamics (MD) simulations provide atomistically detailed movies of the structural and functional dynamics of biological macro-molecules. However, the majority of important dynamic processes in the cell involve broad length and time scales. A large fraction of such processes are rare over the timescale of the simulation. Energy barriers higher than thermal energy trap the simulated system in conformational basins of attraction, impeding proper sampling of all relevant states. Examples of rare processes include protein folding, conformational transitions, ligand binding and unbinding etc., which in most cases involve ~ 10^4^ – 10^6^ atoms including the natural solution environment. Despite phenomenal advances in computing hardware, atomistic MD simulations of such large systems still go, typically, up to multiple microseconds only. This is many orders of magnitude smaller than the timescale relevant to biological function, which is often in the range of seconds to hours.

The current study focuses on protein-ligand interactions; its adequate sampling is pivotal to computer-aided drug design. Molecular dynamics (MD) simulations provide mechanistic insights into such interactions at atomistic detail, and is one of the essential tools in the repertoire of the pharmaceutical research community. A wide range of alchemical free energy calculation methods^1–3^ and enhanced sampling methods (involving external biasing force)^4–7^ have been developed over the past few decades to calculate the binding free energy of a protein-ligand complex. Although the virtual screening of potential inhibitors is currently based on the binding free energy, the efficacy of a drug molecule is often dependent on the binding and unbinding kinetics or the residence time.^8, 9^ It is difficult to compute the kinetic properties from traditional enhanced sampling simulations, as the dynamics become non-physical due to the application of artificial biases (although there are methods to recover kinetics from simple constant force or constant velocity steered molecular dynamics^10–13^). On the other hand, using brute force MD simulation, one needs to sample multiple binding and unbinding events to obtain converged results for kinetic properties. This requires a simulation time many times higher than the timescale of one event, which itself is beyond the reach of even the most powerful supercomputers. This results in a dire need to develop theoretical methods and computational algorithms to make quantitative predictions about the kinetics of long timescale processes such as rare events from short timescale trajectories.

A category of methods involves transition path sampling (TPS), a concept introduced by Pratt^14^ and later developed by Chandler and co-workers,^15, 16^ to simulate transitions across energy barriers. Instead of applying external bias, path sampling methods utilize the statistical properties of the unbiased trajectory ensemble to compute experimental observables such as the kinetics of conformational transition or ligand unbinding, as well as molecular scale properties like ligand release pathways and mechanism.^17^ A different path sampling approach is the weighted ensemble (WE) method of Huber and Kim,^18^ which belongs to a broader category of variance-reduction algorithms that use “splitting” in the framework of Monte Carlo sampling (see, e.g., Kahn^19^). The WE method was further developed by Zuckerman, Chong and collaborators (see e.g., Ref.^20^); it also was established that the weighted ensemble is statistically exact.^21^ In this approach, the conformational space between the initial and final state is discretized into multiple bins and a number of short trajectories are propagated from the starting bin. Trajectory segments are split or merged when they enter a new bin to keep an equal number of trajectories in each bin. Appropriate weights are assigned to the new set of trajectories to conserve the total probability. It allows for the sampling of fast moving but low-weight trajectories that reach the final state well before the mean first passage time; this facilitates the calculation of converged kinetics, free energy and pathways at a relatively low computational cost. With the implementation in the open source software WESTPA, ^22^ the weighted ensemble method has seen a wide range of applications including folding and conformational transitions in proteins,^23–25^ formation of host-guest complexes,^26^ protein-peptide^27^ and protein-protein binding,^28^ ion permeation through protein channels, ^29^ viral capsid assembly^30^ and many others. Many new variants, as well as new analysis schemes for the traditional WE approach, have emerged in recent years, including WExplore,^31^ resampling of ensembles by variation optimization (REVO),^32^ history augmented Markov State Modeling (haMSM),^33^ the RED scheme,^34^ minimal adaptive binning (MAB),^35^ and micro-bin analysis.^36^ Particularly, the WExplore and REVO algorithms have been successfully applied to study the pathways and kinetics of protein-ligand dissociation,^32, 37, 38^ even for systems with residence times as high as seconds to minutes.^39, 40^

Another popular approach to study the kinetics of biophysical rare events is milestoning,^41–43^ which belongs to the larger category of trajectory stratification.^44–47^ In milestoning, multiple interfaces are placed along a reaction coordinate, and short MD trajectories are propagated in between the interfaces, which thus serve as milestones for the progress of the transition of interest. Analyzing the milestone-to-milestone transition statistics via a statistical framework, ^43^ the kinetics and free energy profile are estimated. This method has also been implemented in the software tools miles,^48^ ScMile^49^ and SEEKR,^50^ and has been used to study a variety of complex biological problems including protein allosteric transitions,^51^ membrane permeation by small molecules,^52–55^ protein small molecule interaction,^50, 56–58^ simple ligand-receptor binding, ^59^ peptide transport through protein channels,^60, 61^ DNA protein interaction,^62^ protein conformational dynamics^63^ etc. Apart from the necessity of having a predefined reaction coordinate, the milestones need to be placed far apart to preserve the assumption of Markovianity. ^42^ This itself increases the computational cost significantly, leaving aside the fact that two independent studies have shown that the majority of the total computational effort in milestoning simulation is spent on sampling along the milestone interfaces to generate starting structures in accordance with the equilibrium distribution.^60, 64^

A different variant of the milestoning approach has been developed: Markovian Milestoning with Voronoi Tesselation (MMVT),^55, 65^ which removes the necessity of performing additional sampling along milestone interface, reducing the overall computational cost to a large extent. The application of MMVT remained rather limited, being used primarily for studying small molecule transport through transmembrane proteins,^66–69^ substrate translocation through ATPase motor, ^70^ and the CO entry in myoglobin. ^71^ Only recently, the Markovian milestoning approach has been tested on ligand-receptor binding for crown-ether host-guest complexes and for the dissociation of a benzamidine ligand from the trypsin protein. ^64^ Despite cutting down the computational cost in sampling at the milestone interface, this approach still suffers from the Markovian assumption and can be significantly expensive for complex systems. ^64^

In our previous work, we attempted to improve the milestoning scheme by accelerating transitions between distant milestones via the application of directed wind forces. ^72^ This technique did increase the number of energetically uphill transitions, but the statistical properties of the computed observables were not significantly better. ^73^ More recently, we proposed the combined Weighted Ensemble Milestoning (WEM) scheme, where we performed WE simulations in between milestones to accelerate the convergence of the transition between adequately spaced milestones. ^74^ The WEM method not only produced accurate prediction of kinetics, free energy and time correlation function for small molecular systems like alanine dipeptide, ^74^ but we could also reproduce protein-ligand binding and unbinding rate constants and binding affinity, previously obtained from 30 *μ*s equilibrium simulation,^75^ in less than 100 ns of WEM simulation. ^76^

Yet, the current methodology and the implementation of WEM have a few drawbacks. First, the sampling of the degrees of freedom perpendicular to the reaction coordinate (RC) is significantly poor, particularly in situations where slow conformational changes of the protein are coupled to the ligand unbinding. ^76^ This can potentially be rectified by using multiple starting states on the milestone interface sampled from long umbrella-sampling simulations at the expense of a manifold increase in the computational cost similar to traditional milestoning. Second, the choice of the milestoning reaction coordinate (RC) is arbitrary and can possibly impact the quality of the results, depending on the complexity of the underlying free energy landscape. Moreover, a major hindrance of the large scale application of WEM technique is the complexity of the simulation protocol, which requires propagating many short trajectories and stopping them upon reaching a nearby milestone. ^76^ It requires frequent monitoring of the trajectory as well as frequent communication to the dynamics engine to stop the propagation if the progress coordinate reaches a particular value; this makes the WEM algorithm particularly inefficient to implement in Graphical Processing Unit (GPU) hardware.

We, thereby, present a novel Markovian Weighted Ensemble Milestoning (M-WEM) approach, in which we combine weighted ensemble with soft-wall^65^ based Markovian Milestoning, in an attempt to mitigate the deficiencies and improve the performance of the weighted ensemble milestoning technique. We first provide a detailed description of the theory of Markovian milestoning and the M-WEM approach. We then show the application of this method to the two-dimensional Müller-Brown potential, the conformational transition of alanine dipeptide, and the dissociation and association of the trypsin-benzamidine complex, a protein ligand system with a residence time beyond millisecond. The choice of the trypsin-benzamidine complex is inspired by the fact that many existing path sampling and enhanced sampling methods have been applied on this system, including Markov State Modeling (MSM),^77, 78^ Metadynamics,^79^ Adaptive Multilevel Splitting (AMS),^80^ Milestoning,^50^ MMVT, ^64^ WExplore, ^38^ and REVO. ^32^ So, we compare the accuracy of the results and the performance of M-WEM with these existing techniques, as well as with the experimental rate constants and free energy values obtained by Guillian and Thusius. ^81^ We also discuss a new approach to construct multidimensional free energy landscapes via post-analysis of MMVT and M-WEM trajectories obtained using a one-dimensional reaction coordinate, with a potential application in systems were orthogonal degrees of freedom are strongly coupled with the reaction coordinate.

## Theory

### Markovian Milestoning

The theoretical details of the Markovian milestoning with Voronoi tessellation (MMVT) approach is described elsewhere.^55, 64, 65^ Here, we provide only a brief description relevant to the current work.

In MMVT, the configurational space is discretized into Voronoi cells. A flat bottom potential is applied to each cell with half-harmonic walls placed at each milestone interface, preventing the trajectories from escaping out of the Voronoi cells. For a 1-dimensional reaction coordinate, used here, the flat bottom potential has the expression:

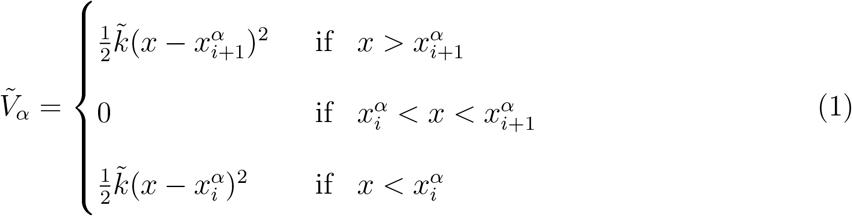

where *α* is the cell index, 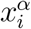 is the value of the reaction coordinate at the milestone *i* at the boundary of the cell *α*, and 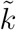 is the force constant; the total number of cells is Λ and the total number of milestones is *M*. One or more unbiased trajectories are propagated in each cell. The trajectories which cross the milestone interface are reflected back into the cell by the half harmonic restraint. As a result, the trajectories remain confined into one cell and perform many transitions between the milestones interfaces constituting the boundaries of the cell. The portions of the trajectory outside the cell are to be discarded before performing further analysis. This protocol is referred to as the *soft wall* restraint^65, 66^ which we adopt in the current work. Alternatively, a *hard wall* restraint^55, 64^ can also be used where the direction of velocity is switched when a trajectory crosses a milestone.

From these confined trajectories, the transition counts between milestones are recorded. A flux matrix 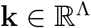 is constructed whose elements are given by:

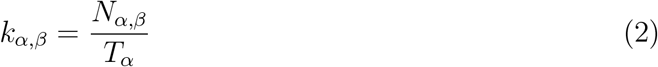

where *N_α,β_* is the number of transitions from cell *α* to cell *β* recorded from a trajectory propagated for time *T_α_* in the cell *α*. The equilibrium probability for each cell (π_*α*_) is the obtained by iteratively solving the linear equation (Eq. 3) in a self-consistent manner under the constraint of a constant total probability of one:

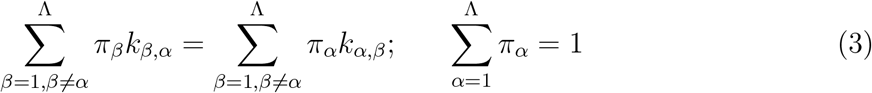

The free energy profile at each cell is then computed as:

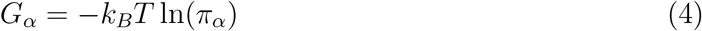

For calculating kinetics, the transition matrix 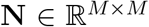 and the lifetime vector 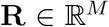 are constructed, whose elements are computed as follows:

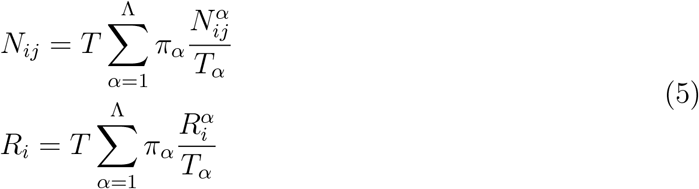

where 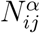 is the number of times a trajectory in cell *α* collides with milestone *j* after having last visited milestone *i*; 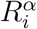 is the cumulative time the trajectory spends in cell *α* visiting milestone *i* and before reaching any other milestone. *T* is a constant for dimensional consistency, which is not necessary to compute because it cancels out at a later stage. A rate matrix 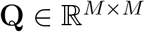 is then defined as:

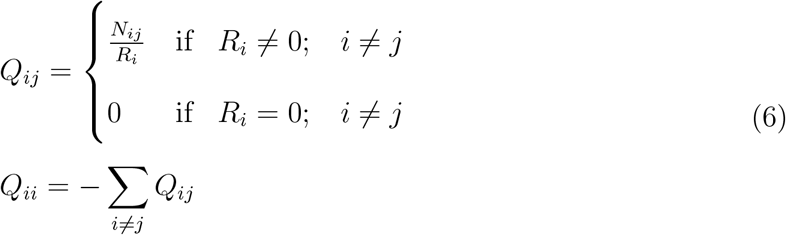

Considering milestone *M* is the target milestone, the mean first passage time of the process can be computed as:

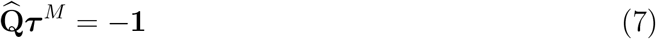

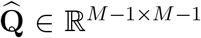 is the matrix obtained by deleting the last row and column of **Q**. **1** is a unit vector with *M* – 1 elements, and 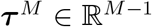 is the vector with entries 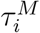 that are the MFPTs from milestone *i* to milestone *M*.

### Markovian Weighted Ensemble Milestoning (M-WEM)

In the current work we introduce the Markovian Weighted Ensemble Milestoning (M-WEM) approach, where the conventional MD trajectories in the Markovian milestoning framework are replaced by weighted ensemble simulation. A schematic representation of the M-WEM protocol is depicted in Fig. 1. WE bins are placed along the reaction coordinate in-between the milestone interfaces, as well as along a different coordinate to accelerate sampling along the milestone interface. The additional non-RC coordinate should ideally be locally orthogonal to the RC, but this is not a necessary condition. WE simulation is performed in this 2D progress coordinate space using the recently developed minimal adaptive binning (MAB) scheme. ^35^ As opposed to the traditional fixed binning scheme, the MAB approach adaptively changes the bin boundaries during the course of simulation, avoiding the requirement of an arbitrarily chosen predefined set of bins. It also provides an increased sampling of the conformational space. As the total number of occupied bins remains virtually unchanged throughout the simulation, the maximum amount of computational resources needed for the simulation can be easily estimated beforehand.^35^ A stochastic integrator, e.g., for the Langevin equation, is needed to propagate the dynamics to ensure that the new set of trajectories, generated after a splitting event, follow different paths despite emerging from a single parent trajectory.

**Figure 1:**
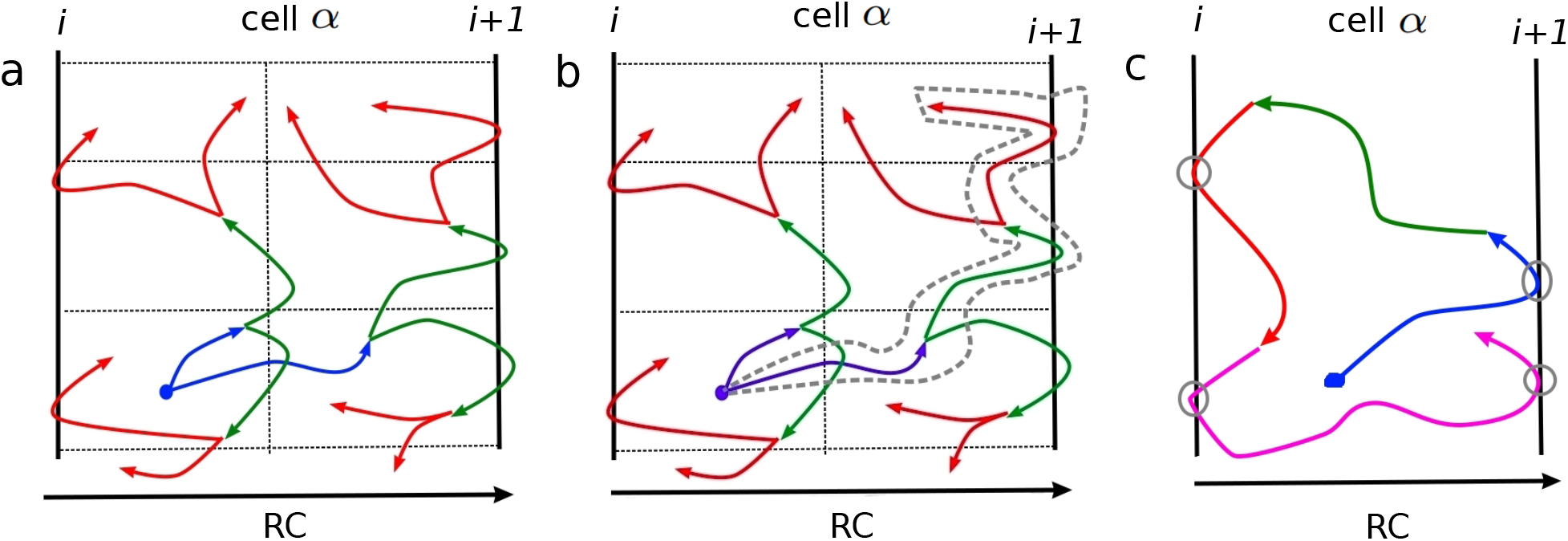
A schematic representation of the M-WEM simulation protocol. The thick lines indicate milestones (labeled as milestone index *i* and *i* + 1). The dotted lines indicate a WE bin boundaries (which are adapted during the simulation but in this figure we show fixed bins for clarity). Trajectories for different WE iteration is shown in different color scheme: Iteration 1: blue, iteration 2: green, iteration 3: red, and iteration 4: pink. (a) First, WE simulation is performed with harmonic walls placed at the milestone interfaces allowing for the trajectory to bounce back and forth. (b) The propagation history of individual trajectories are traced back from the last iteration (an example trace is highlighted with gray dashed line). (c) The milestone crossing events (gray circles) are recorded from each trace, and are used in subsequent analysis.

Unlike the MMVT approach with conventional MD, the WE trajectories hitting the milestones will have different weights. To properly take into account this effect, we take all the trajectory segments at the last iteration and trace them back to the first iteration to obtain separate trajectory traces. The weight of each trajectory trace is set equal to the weight of the corresponding trajectory segment in the last iteration.

The total number of trajectory traces in cell *α* (*M_α_*) is equal to the number of occupied bins × the number of trajectories per bin in the final iteration. The elements of the flux matrix **k** in this formalism are given by:

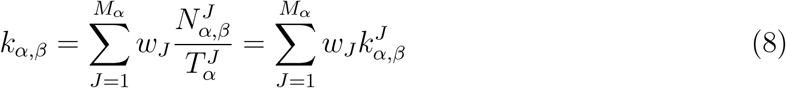

where *w_J_* is the weight of the *J*th trajectory trace. 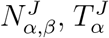, and 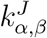 have similar definitions as in Eq. 2 except that they are computed just from the *J*th trace. The equilibrium probability distribution and the free energy profile are computed from the elements of the flux matrix obtained from Eq. 8 using Eq. 3 and 4, respectively.

For calculating kinetics, the *N_i,j_* and the *R_i_* matrix elements are to be constructed taking into account the different weights of the trajectory traces. The new transition matrix element becomes:

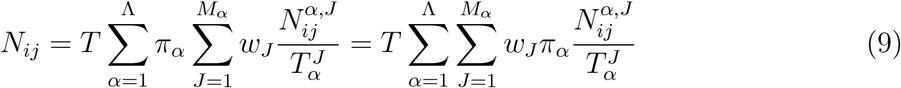

where 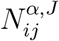 has the same definition as 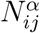 in Eq. 5 except it is for the *J*th trajectory trace. Now we define a pseudo transition matrix 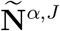, which is identical to the transition matrix **N** but computed only from one (*J*th) trajectory trace in the cell *α*. Its elements are given by:

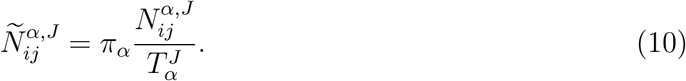

Now the total transition matrix **N** can be obtained as

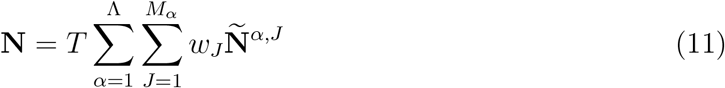

Similarly, the lifetime vector **R** is given by:

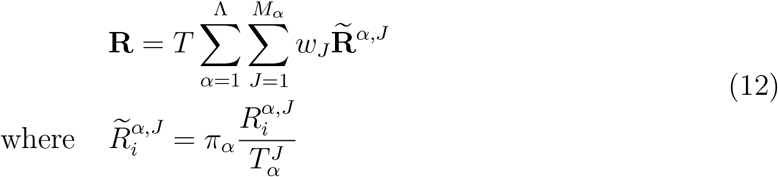

and 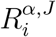 is equivalent to 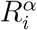 but computed only for the *J*th trajectory segment. Using the **N** and **R**, the rate matrix **Q** is computed using Eqs. 6 and 7.

### Error Analysis

Error analysis of milestoning-based simulations can be performed in a few different ways, primarily by generating an ensemble of rate matrices (**Q**). Then the desired properties (such as the MFPT) are calculated from many sample matrices and the uncertainty is estimated. When working with transition matrices (as in traditional milestoning), the kernels can be sampled from a beta distribution. ^60^ Similar to our previous work,^74, 76^ we generated the ensemble of rate matrices by sampling from a Bayesian type conditional probability,^56, 82, 83^ given by

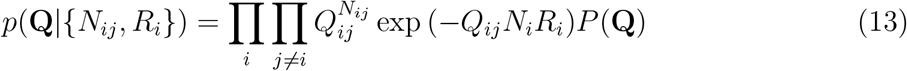

where *p*(**Q**) is a uniform prior, *N_ij_* is the number of trajectories transiting from milestone *i* to *j* and 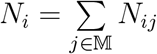, where 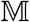 is the set of all milestones. We sampled the **Q** matrices from the distribution in Eq. 13 using a non-reversible element exchange Monte-Carlo scheme.^84^ One randomly chosen off-diagonal element and the diagonal element of the corresponding row of **Q** are updated to generate a new rate matrix **Q**′.

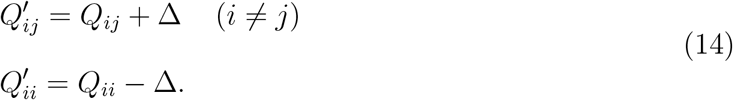

where Δ is a random number sampled from an exponential distribution of range [–*Q_ij_*, ∞) with mean zero. The new matrix **Q′** is accepted with a probability equal to min(1, *p*_accept_) where:

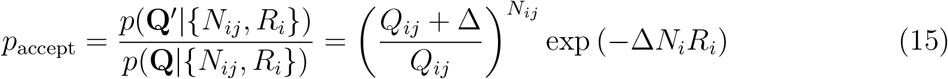

As each step modifies only one element, the sampled matrices are highly correlated.^84^ So we only considered the matrices sampled every 50 steps for uncertainty estimation. We recomputed the MFPT obtained from them using Eq. 7 and calculated mean and 95% confidence intervals for all MMVT simulations.

For M-WEM simulations, estimating errors using non-reversible element exchange Monte-Carlo can be difficult. The expression for *p*_accept_ (Eq. 15) has exponential dependence on the number of transitions *N_ij_* and number of trajectories reflected from milestone *i* (*N_i_*). In the case of MMVT simulations all such transitions have equal weight, unlike in M-WEM where the weights of transiting trajectories can vary to a large extent, often over many orders of magnitude. So using the expression in Eq. 15, which counts all transitions with equal importance, one is unable to sample the ensemble of transition matrices accurately in our M-WEM approach. For each observable (MFPT, free energy, etc.), we computed its estimate over an M-WEM simulation of increasing number of total iterations for each cell. We then compute the mean and 95% confidence interval of the sampled observables after the simulation is converged. The uncertainties obtained using this procedure capture the effect of the fluctuations of an observable around the mean after the simulation has converged. Such approach is commonly used in the literature to compute the error estimates of free energy profiles, as discussed, for example, in our previous work Ref.^85^ A more elaborate version of this technique is called block averaging, ^86^ which is a statistically rigorous way to compute error estimates in molecular dynamics simulations.^87^

Specific details of error analysis for each test system is mentioned in the Computational Methods section. For the M-WEM technique, the derivation of a more rigorous approach for error analysis, similar to the one described in Eqs. (13) - (15) will be addressed in future work.

### Efficiency Analysis

Because of the difference in the uncertainties of the measured values from different methods, a direct comparison between the simulation time or convergence time may not always correctly represent the relative efficiency between different approaches. To address this issue, we used the algorithmic efficiency metric, *η*, which takes into account the variance of the data apart from the total simulation time. ^74^ This metric is evaluated as:

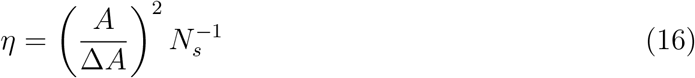

where *A* is the measured quantity and Δ*A* is the uncertainty of the measurement. *N_s_* is the number of force evaluations performed to calculate *A* (although it can be thought of as the simulation time if expressed in units of time instead of time steps). In brief, *η*^−1^ is the number of force evaluations required to measure our observable *A* with an uncertainty equal to its mean.^18^ The lower the value of *η*^−1^, the better the efficiency. For the first two systems, the Müller-Brown potential and the alanine dipeptide, we reported the value of *η*^−1^ where we calculated the error bars from multiple runs. We did not report *η*^−1^ for each individual run because the error of MMVT calculations, computed using Bayesian analysis, and the error bars of the M-WEM approach computed from averaging over iterations are not directly comparable. For the same reason, we do not report an algorithmic efficiency for the trypsin-benzamidine system, for which *η* will also depend on the quantity we are measuring.

### Calculation of *k*_off_ and *k*_on_

For the protein-ligand binding problem considered in this paper, we computed the unbinding rate constant (*k*_off_) as

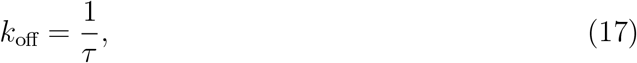

where *τ* is the ligand residence time, which is equivalent to the MFPT of transitioning from the bound state milestone to the unbound state milestone, and which, in turn, can be computed from the milestoning or M-WEM framework described above.

We noted in our previous work that the ligand-binding kinetics is often diffusion dominant. ^76^ So the rate determining step can be the arrival of the ligand on the outermost milestone surface, rather than going from the outermost milestone to the binding pocket. (Whether this is true, for a specific system, should be carefully evaluated on a case-by-case basis.) In our previous work, we delineated an analytical method to combine diffusion theory with milestoning transition kernel integration to compute the binding rate constant *k*_on_.^76^ We begin with the expression of the diffusion-dependent arrival rate of a small molecule on the surface of a sphere of radius *r*:^88^

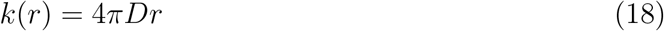

where *D* is the diffusion constant of the ligand in water. We make two modifications to this expression. First, we assume that the only conformations that can lead to binding are in the space explored by the ligand in the outermost milestone. So we scale the rate by the surface coverage factor *α*. The concept of coverage factor has been described more elaborately in our previous work. ^76^ To mention it only briefly here, *α* is the fraction of the surface area of the outermost milestone interface that is accessible to the ligand. Details of the calculation of *α* in this work are slightly different from our previous work. Instead of performing a Monte-Carlo simulation, here we create a 2D grid on the interface of the outermost milestone based on the solid angles. Then we calculate how many of those bins are occupied by the center of mass of the ligand in the outermost milestoning cell. The coverage factor *α* is then computed as

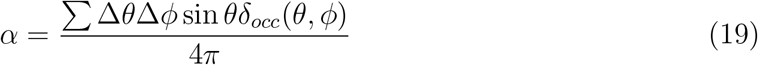

where *δ_occ_*(*θ, ϕ*) = 1, if the surface element Δ*θ*Δ*ϕ* contains the angular coordinate of the ligand, and zero otherwise.

Second, we consider that a ligand can only reach the bound state if it moves towards the protein from the outermost milestone surface. So we further scale the arrival rate by *K*_*M,M*–1_, the transition probability of going from the last (*M*th) milestone to the previous milestone. But as the transitional kernel is not directly available in the current Markovian milestoning framework, we compute this inward transition probability from the **N** matrix.

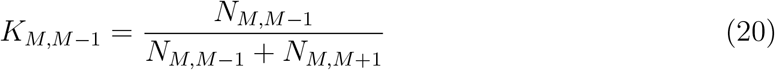

As *N*_*M,M*+1_ becomes undefined when *M* is the last milestone, we perform this analysis with the last but one milestone. Including these factors, the flux of binding trajectories through that milestone becomes

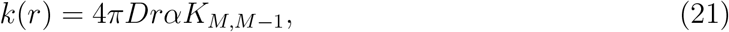

from which *k*_on_ is calculated in M^−1^s^−1^ units by multiplying the flux with Avogadro’s number *N_av_*

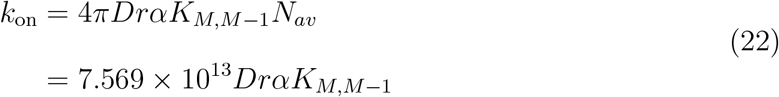

In the last step we used the diffusion constant of small molecule in cm^2^ s^−1^ unit and the *r* is provided in Å.

### Reconstruction of free energy landscape

The trajectories confined in the different Voronoi cells can be used to construct a higher dimensional free energy landscape, both for Markovian milestoning and M-WEM simulations. The trajectory data is first histogrammed in appropriate collective variables to obtain different independent histograms for each individual cell Λ. A weighted sum of these histograms are then performed to obtain the equilibrium high dimensional probability distribution

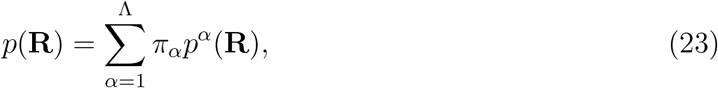

where *p*(**R**) is the probability density along the collective variable space **R**, and *p^α^*(**R**) is the same obtained from the histogram only in the cell *α*. In case of M-WEM the *p^α^*(**R**) is obtained from multiple (*M_α_*) trajectories of different weights as:

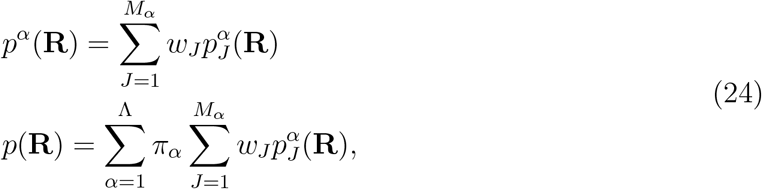

where 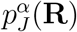 is the histogram of the *J*th trajectory in cell *α* in the CV space **R**. The free energy landscape is then reconstructed using,

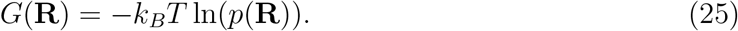

We demonstrate this approach for alanine dipeptide in the Results section.

### Committor

The committor of a point in a conformational space is the probability of a trajectory starting from that point to reach the final state before visiting the initial state.^89^ Recent work by Elber et al. established that it is possible to calculate the committor at the milestone interfaces.^48^ But, to the best of our knowledge, such approach has not been applied to Markovian milestoning techniques so far. We performed the committor calculation in the following way. First, we constructed a transition kernel (**K**), equivalent to conventional milestoning, from the **N** matrix:

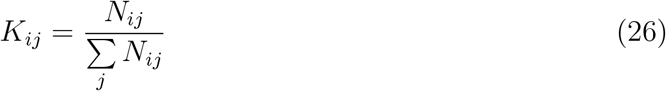

The transition kernel **K** is then modified with a boundary condition which ensures that the flux through the final state will remain “absorbed” there and will not return to the previous milestones. This is achieved by replacing the last row of the **K** matrix with zeros, except for the element corresponding to the last milestone for which the value is one.^48, 90^ For a three-milestone model, this can be illustrated as:

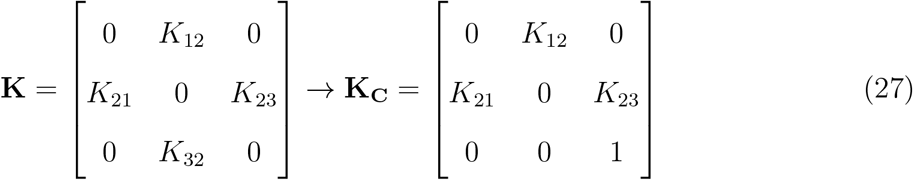

The vector containing the committor values of each milestone, **C**, is then calculated as:

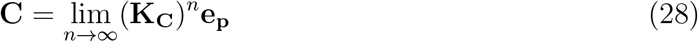

where **e_p_** is a unit vector, all elements of which, are zero except for the one corresponding to the final milestone. Multiple powers of **K_C_** are computed numerically until the committor converges. The results are considered to be converged when the change in the norm of the **C** vector to be less than 10^−3^.

## Computational Methods

We tested the Markovian Weighted Ensemble Milestoning approach on a toy model system of 2D Müller-Brown potential, conformational transition in alanine dipeptide and on the millisecond timescale protein-ligand unbinding in the trypsin-benzamidine system. In the first two systems, we performed long equilibrium simulation and Markovian milestoning simulation to compare with our M-WEM results.

### Müller-Brown Potential

The two-dimensional Müller-Brown potential^91^ is defined as

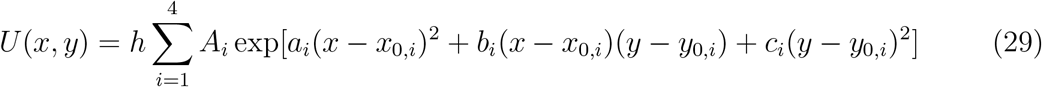

where *A* ∈ {−200, −100, −170, 15}, *α* ∈ { −1, – 1, −6.5, 0.7}, *b* ∈ {0, 0, 11, 0.6}, *c* ∈ { – 10, −10, −6.5, 0.7}, *x*_0_ ∈ {1, 0, −0.5, −1}, *y*_0_ ∈ {0, 0.5, 1.5, 1}, and *h* = 0.04. This system has a non-linear transition path with barrier height about 4.5 *k_B_T*. For the purpose of this model, we set *k_B_T* = 1.

To obtain a benchmark of the kinetics, a long overdamped Langevin dynamics simulation was propagated starting from (*x, y*) = (−0.5, 1.5), which corresponds to the minimum *A* in Figure 2. The minimum *B* is chosen to be the target state. The simulation was propagated for 10^7^ time steps, capturing 347 back and forth transitions. The free energy landscape obtained from this equilibrium trajectory is depicted in Figure 2. The mean first passage time (MFPT) to go from minima *A* to *B* is depicted in Table 1.

**Figure 2:**
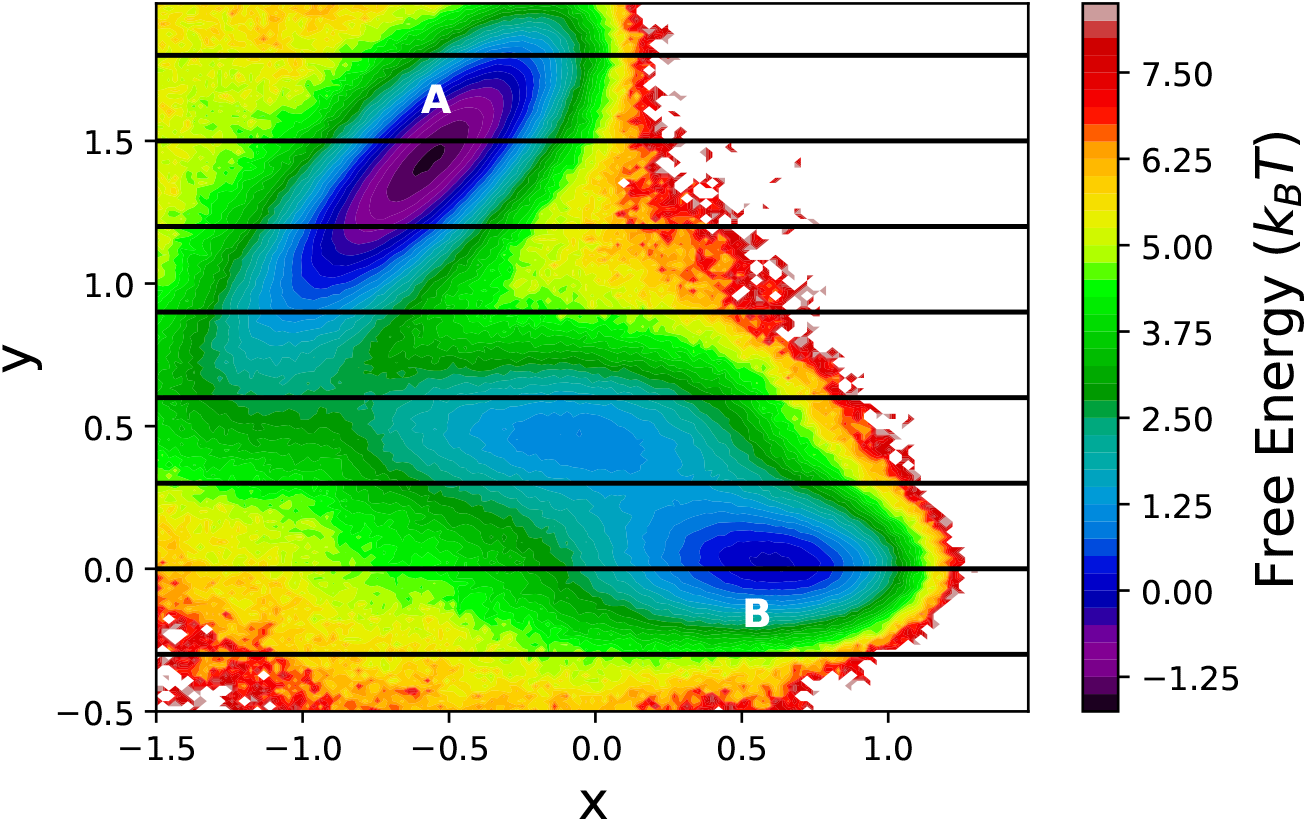
The free energy landscape of the Müller Brown potential explored using 10^7^ steps of over-damped Langevin dynamics simulation. The position of the milestones, used in MMVT and M-WEM calculations, are shown in black lines. The two minima relevant to this study are marked as A and B.

**Table 1:** Results of conventional Langevin dynamics, MMVT and M-WEM simulations for the Müller Brown potential

| Method | MFPT<br>($\times 10^3$ steps) | Simulation time<br>($\times 10^6$ steps) | $\eta^{-1}$<br>($\times 10^6$ steps) |
| --- | --- | --- | --- |
| Regular over-damped LD | $25.2 \pm 2.9$ | 10 | 0.13 |
| MMVT (trial 1) | $12.3 \pm 0.4$ | 10.5 | - |
| MMVT (trial 2) | $13.1 \pm 0.3$ | 10.5 | - |
| MMVT (trial 3) | $13.2 \pm 0.5$ | 10.5 | - |
| MMVT (average)<sup>a</sup> | $12.9 \pm 1.2$ | - | 0.09 |
| M-WEM (trial 1) | $22.2 \pm 3.2$ | 2.6 | - |
| M-WEM (trial 2) | $17.3 \pm 1.6$ | 2.6 | - |
| M-WEM (trial 3) | $28.2 \pm 4.6$ | 2.6 | - |
| M-WEM (average)<sup>a</sup> | $22.6 \pm 13.6$ | - | 0.94 |

<sup>a</sup>Computed from the three independent trials

For milestoning simulations, 8 milestones are placed at *y* = {−0.3, 0.0, 0.3, 0.6, 0.9, 1.2, 1.5, 1.8} at equal intervals. The reaction coordinate is chosen to be parallel to the *y* axis. This poor choice of RC was made intentionally to represent realistic situations where the arbitrarily chosen empirical RCs are used to study complex biomolecular processes with many coupled degrees of freedom. MFPTs were computed for transition from *y* = 1.5 to *y* = 0.0 for both the milestoning based methods.

For Markovian milestoning (MMVT) simulations, an overdamped Langevin dynamics simulation is propagated in 2D in all the seven cells in the spacing between 8 milestones. A half-harmonic wall is applied at both ends of the cell (milestones) with a force constant 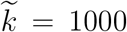, to confine the trajectories within the cell. Each simulation is propagated for 1.5 × 10^6^ steps in each cell. Transition statistics between each milestone pair is computed using the method described in theory section, and the MFPT has been computed.

For the M-WEM method, the procedure remained identical to the MMVT approach, except weighted ensemble simulation was performed in between the milestone interfaces as opposed to conventional dynamics. The 2D adaptive binning (MAB)^35^ was employed along *x* and *y* directions with 5 bins per dimension with 4 trajectory segments per bin. This led to 33 bins in total, including the additional bins in each direction for the most forward, backward and the bottleneck trajectories.^35^ A total of 300 iterations of WE simulation are performed in each cell, with a recycling interval of 10 steps. The transition rate matrices and MFPTs were computed every 10 iterations of WE simulation in each cell between iteration 260 and 300. The mean and error estimates were performed on the 5 sampled data points for the MFPT values between iteration 260 and 300.

### Alanine Dipeptide

The conformational transition of alanine dipeptide was simulated in the gas phase. The system was set up following the protocol described by Wei and Elber. ^49^ The 22-atom system is modelled using the CHARMM22 force field^92^ with a 10 Å cut-off distance for the inter-atomic interactions. A time step of 0.5 fs was used and all the bonds between heavy atoms and hydrogen were constrained using the SHAKE algorithm.^93^ All simulations were performed using the NAMD 2.13 package^94^ with the colvars^95^ module. The conformational change can be described adequately in the 2D coarse space of two backbone torsion angles Φ and Ψ. Half harmonic walls with a mild force constant (0.04 kcal mol^−1^ deg^−2^) are placed at the value of ±175 for both the Φ and Ψ angles to avoid transitions along the edges of the free energy surface (i.e., to remove periodicity).^49^ This will ensure that we observe the transition along the center of the free energy map.

The barrier height for the conformational transition of gas-phase alanine dipeptide is very high (> 10 kcal/mol) and it is very difficult to observe direct transitions at room temperature. Following the earlier work^49^ we performed the simulations at 600K temperature which allowed us to sample 285 transitions between the two free energy minima in 500 ns equilibrium simulation.

For both MMVT and M-WEM simulation, the reaction coordinate was chosen to be the Φ dihedral angle and milestones were placed at Φ = −80°, −60°, −40°, −20°, 0°, 20°, 40°, 60°, and 80°. The initial and final states were chosen to be the milestones at Φ = −80° and Φ = 60°. In case of MMVT simulation, 5 ns of conventional MD simulation was propagated in each cell confined between the two consecutive milestones, leading to a total computational effort of 40 ns. (The trajectories were extended to 10 ns with no difference in results. So the result of 5 ns simulation is reported). A force constant of 4 kcal mol^−1^ deg^−2^ were applied in the harmonic walls placed at the milestones, to confine the trajectories in between the milestones. The portion of the trajectories outside the cell has been removed prior to further analysis.

In case of M-WEM simulations, 5 WE bins were placed for each cell in Φ and Ψ coordinates leading up to 33 bins in total including separate bins for the forward, backward and the bottleneck trajectories. Four trajectory segments were propagated in each occupied bin. The progress coordinate values were recorded at very frequent interval (10 fs) to record the time of milestone crossings as accurately as possible. A total of 100 iterations of WE simulation are performed in each cell, with a recycling time of 1 ps. The transition rate matrices and MFPTs were computed every 2 iterations of WE simulation in each cell between iteration 2 and 10 and every 10 iterations between iteration 10 and 100, to monitor the convergence of the results. The convergence plots and related discussion is provided in the supporting information. The mean and error estimates were performed on the 5 sampled data points for the MFPT values between iteration 60 and 100.

### Trypsin-Benzamidine Complex

The system setup for the trypsin-benzamidine complex is identical to the work by Votapka et al. ^50^ The structure, parameter and topology files were obtained from the authors of Ref. 50. We point the reader to their original publication ^50^ for more details. To mention briefly, the atomic coordinates of the protein-ligand complex were obtained from Protein Data Bank (PBD) PDB ID: 3PTB.^96^ The protonation states of ASP, GLU and HIS residues were determined at pH 7.7 which was used in this study to replicate the experimental condition. ^81^ The protein was modelled using AMBER ff14SB force field^97^ and Generalized Amber Force Field (GAFF)^98^ parameters were used for the ligand. The structure was solvated in a truncated octahedron box of TIP4Pew^99^ water molecules and 8 Cl^−^ ions were added to neutralize the system. Overall the system contains ~23000 atoms. All MD simulations were performed using NAMD 2.14b2 package^94^ with a time step of 2 fs. A Langevin integrator with a damping coefficient of 5 ps^−1^ was used to keep the temperature constant at 298 K. A Langevin piston was used to maintain the pressure at 1 atm.

The bound state structure was equilibrated for 10 ns in NPT ensemble. From the end point of this simulation, the ligand was pulled out of the binding pocket using a 10 ns steered molecular dynamics (SMD) simulation. The reaction coordinate (RC) description is identical to previous work,^50^ i.e. the center of mass distance between the benzamidine ligand and the *C_α_* atoms of the following residues near the binding pocket: 190, 191, 192, 195, 213, 215, 216, 219, 220, 224, and 228 (numbered according to PDB: 3PTB). During the SMD simulation, a moving harmonic restraint of 1 kcal mol^−1^ Å^−2^ was applied on the RC with a pulling velocity of ~ 1.5 Å/ns. The collective variables were biased and monitored using the colvars module. ^95^ Representative structures for seeding the milestoning simulations were sampled from the SMD trajectory.

Concentric spherical milestones were placed at the following values of the RC: 1.0, 1.5, 2.0, 2.5, 3.0, 3.5, 4.0, 5.0, 6.0, 8.0, 10.0, 12.0, and 13.0 Å. These values are similar to previous studies^50, 64^ except for a few additional milestones, as we were unable to observe energetically uphill transitions otherwise. The separation between milestones should be such that the transition timescales between one milestone to the other should be larger than the decay time of the velocity auto-correlation function of the RC. We checked this condition in our system, as discussed in detail in the Supporting Information.

The following distinction is worth noting at this point. As in Markov State Modeling (which assumes the formalism of continuous-time Markov chains) the transitions between conformational macrostates^100^ are to be Markovian. However, the dynamics inside the macrostates delimited by the milestones may not necessarily be Markovian, and it is for that dynamics that we check for the decay of velocity autocorrelation. Milestoning theory is built upon two key assumptions: the reaction channels are localized and the committor can be represented as a function of the reaction coordinate only. A sufficient condition to make these assumptions valid is that the dynamics be overdamped. To ensure this condition, milestones need to be placed sufficiently far from each other such that the timescale of transitions between milestones is higher than the decorrelation time of the velocity along the reaction coordinate.^83^

A total of 12 cells were constructed in the spacing between 13 milestones. For each cell, a flat bottom potential (Eqn. 1) is applied with a force constant of 100 kcal mol^−1^ Å^−2^ for the harmonic walls present at the milestones. First, the representative structure (sampled from SMD) is equilibrated at the center of the cell for 1 ns by restraining the RC via a harmonic potential. The force constant was gradually increased to 500 kcal mol^−1^ Å^−2^ over the first 500 ps and kept constant over the last 500 ps. From the end point of the 1 ns equilibration Weighted ensemble (WE) simulations were propagated for 300 iterations with a recycle time *δt* of 2 ps. A two-dimensional MAB scheme was used for the binning. The two progress coordinates were the RC and the RMSD of the ligand with respect to the representative structure (sampled from SMD) corresponding to the specific cell. The progress coordinates were recorded using the colvars module. ^95^ The total computational cost of the M-WEM simulation was approximately 734 ns. The simulation was stopped at multiple points, at an interval of 10 WE iterations between 30 and 300 iterations for each cell. For each set, the trajectory traces were computed using which equilibrium probabilities, free energy profiles and MFPTs between the first milestone (at 1 Å) and the last milestone (at 13 Å) (residence time) was computed. This allowed us to monitor the convergence of residence time over the course of the simulation. The unbinding rate constant *k*_off_ was calculated as the inverse of residence time. Free energy profile, binding rate constant *k*_on_ and committors were calculated following the procedure described in the Theory section. The error bars for all quantities were calculated from the last five iterations sampled (i.e. iteration 160-200 for values reported at iteration 200 and iteration 260-300 for values reported at iteration 300). Before any calculation, the probabilities of the voronoi cells are modified 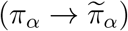 to take into account the Jacobian factor appearing due to the different surface area of milestones with different radius: 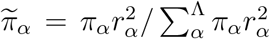 where *r_α_* is the radius of the cell *α* which we choose to be the radius of a sphere equidistant from the two milestones surrounding the cell.

## Results

### Müller-Brown Potential

For the two-dimensional toy model of Müller Brown potential, we performed three independent trials for both MMVT and M-WEM simulations and the results are presented in Table 1. The MFPT of the transition from milestone at *y* = 1.5 to *y* = 0.0, computed using M-WEM approach, shows quantitative agreement with MFPT of the transition from minimum A to minimum B in regular overdamped Langevin simulation. The results of MMVT simulation are off by a factor of ~ 2. Although the simulation time for MMVT and regular MD were comparable, the M-WEM simulations produced converged results with ~ 4 times less computational expense. The algorithmic efficiency of M-WEM, on the contrary, is poorer compared to the MMVT approach because of the larger variance of the MFPTs obtained from the M-WEM method. Although the computational gain is not significant in case of this low dimensional model system, these results serve as a proof of concept of our method in rare event sampling problem. It also indicates that despite the choice of a poor and simplistic RC, accurate MFPTs can be calculated using M-WEM method.

### Alanine Dipeptide

Next, we tested the performance of MMVT and M-WEM methods on the conformational transition of Alanine dipeptide. The results were compared to a 500 ns conventional MD simulation. The free energy landscape along the Φ and Ψ torsion angles for the gas phase Alanine dipeptide (obtained from equilibrium MD simulation) is shown in Fig. 3. The mean first passage time (MFPT) of transition from milestone Φ = −80° to milestone Φ = 60° is in agreement with the MFPT of transition from free energy minima A to B obtained from long equilibrium MD simulation (Table 2). The M-WEM results show slightly better agreement, but the difference is not very significant; in fact, the error bars of the MMVT and M-WEM simulations obtained from independent runs overlap with each other. Both these methods produced accurate results within one order of magnitude less computational cost in comparison to the equilibrium MD. Although the M-WEM simulations took about twice as much computational effort as the MMVT simulation for full 100 iterations, the MFPT results converged as early as in 20-30 iterations (See Supporting Information). Moreover, going from the 2D model of the Müller potential to the molecular model of alanine dipeptide, we see a significant improvement of the algorithmic efficiency as the *η*^−1^ of M-WEM is now slightly lower than that of MMVT. Nonetheless, both have poorer efficiency than regular; the latter yields a tighter confidence interval from ~300 transitions, whereas MMVT or M-WEM uncertainties are estimated from three independent runs.

**Figure 3:**
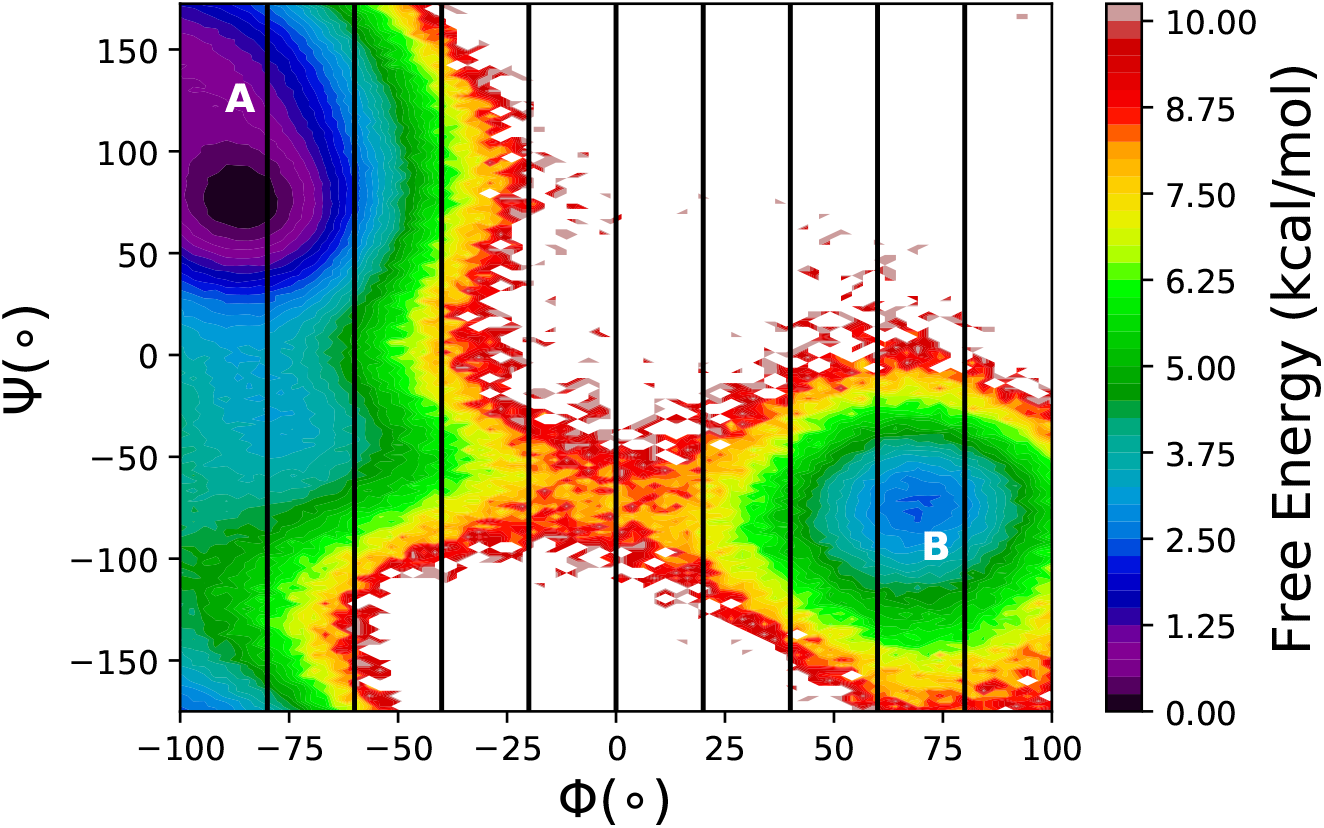
The free energy landscape of the gas phase Alanine dipeptide along the Φ and Ψ torsion angles, from 500ns equilibrium MD simulation. The position of the milestones, used in MMVT and M-WEM calculations, are shown in black lines. The two conformations relevant to this study are marked as A and B.

**Table 2:** Results of conventional MD, MMVT and M-WEM simulations for the Alanine dipeptide.

| Method | MFPT<br>(ps) | Simulation time<br>(ns) | $\eta^{-1}$<br>(ns) |
| --- | --- | --- | --- |
| Regular MD<sup>a</sup> | 1583±188 | 500 | 7.05 |
| MMVT trial 1 | 1465±167 | 40 | - |
| MMVT trial 2 | 951±13 | 40 | - |
| MMVT trial 3 | 984±8 | 40 | - |
| MMVT (average)<sup>b</sup> | 1133±715 | - | 15.93 |
| M-WEM trial 1 | 1690±112 | 86.4 | - |
| M-WEM trial 2 | 1290±69 | 84.4 | - |
| M-WEM trial 3 | 1286±123 | 83.2 | - |
| M-WEM (average)<sup>b</sup> | 1422±576 | - | 13.78 |

<sup>a</sup>From state A to B in figure 3, not from milestone to milestone.
<sup>b</sup>Computed from the three independent trials

The committor values at milestone interfaces for all three trials of M-WEM calculation are depicted in figure 4. The results from different trials are in excellent agreement with each other and all of them shows a committor value of ~ 0.5 for the milestone at Φ = 0°. A committor value of 0.5 indicates the transition state (TS). The milestone at Φ = 0° is indeed present on top of the free energy barrier aka TS as evident from the free energy landscape in Fig. 3.

**Figure 4:**
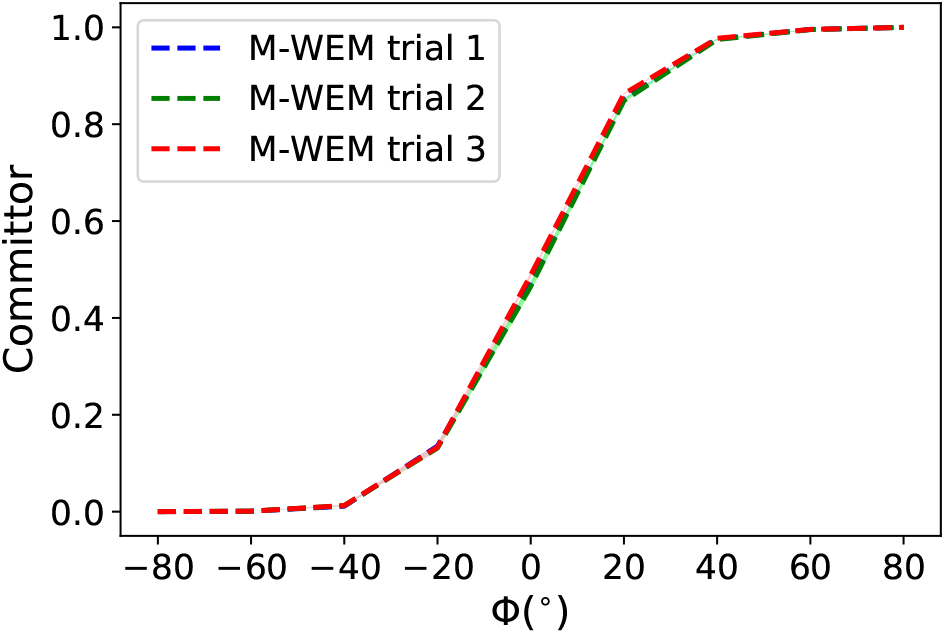
The committor values as a function of the milestoning coordinate Φ for the Alanine dipeptide system.

We applied our free energy reconstruction protocol to recover the free energy landscape along the Φ and Ψ degrees of freedom. The crude probability distribution for each cell is obtained by histogramming the M-WEM and MMVT simulation data projected on those two degrees of freedom. This unscaled distribution for individual cells (*p^α^*(Φ, Ψ)) as obtained from the M-WEM calculation is shown in figure 5a. Then the true probability distribution (*p*(Φ, Ψ)) is computed by re-scaling the distributions corresponding to each cell with the weight of their probabilities obtained using Eq. 3 (Fig. 5b).

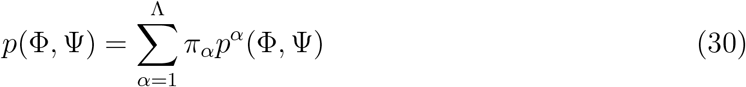

**Figure 5:**
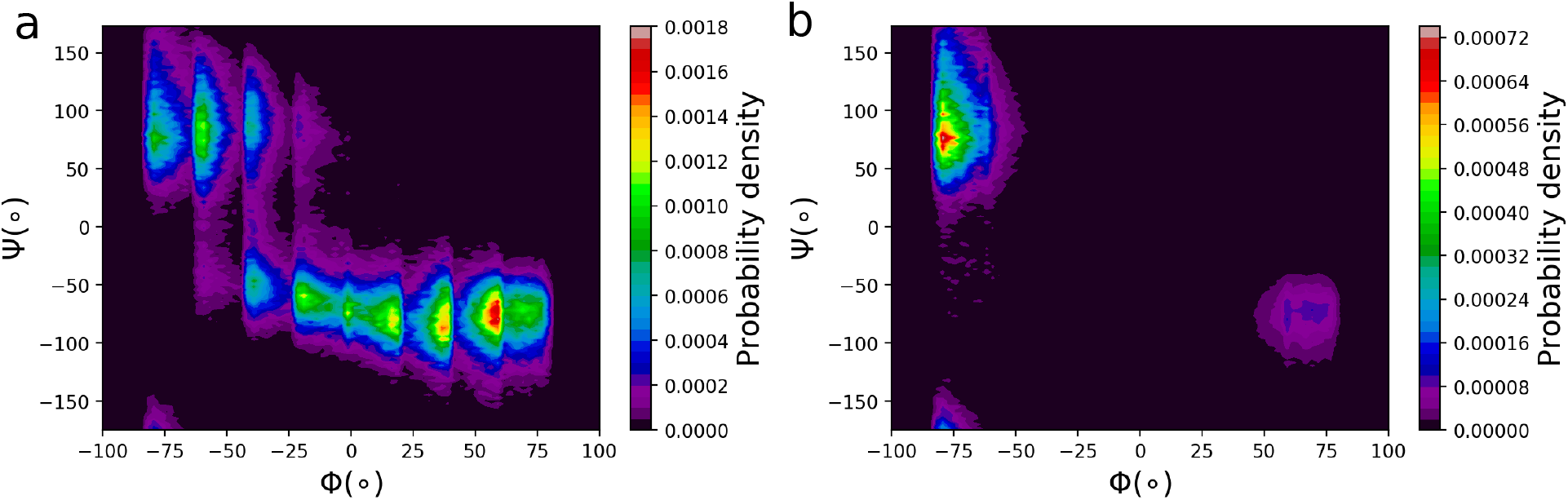
Reconstruction of equilibrium probability distribution (b) from raw unscaled probability distribution from M-WEM trajectories in each cell (a).

**Figure 6:**
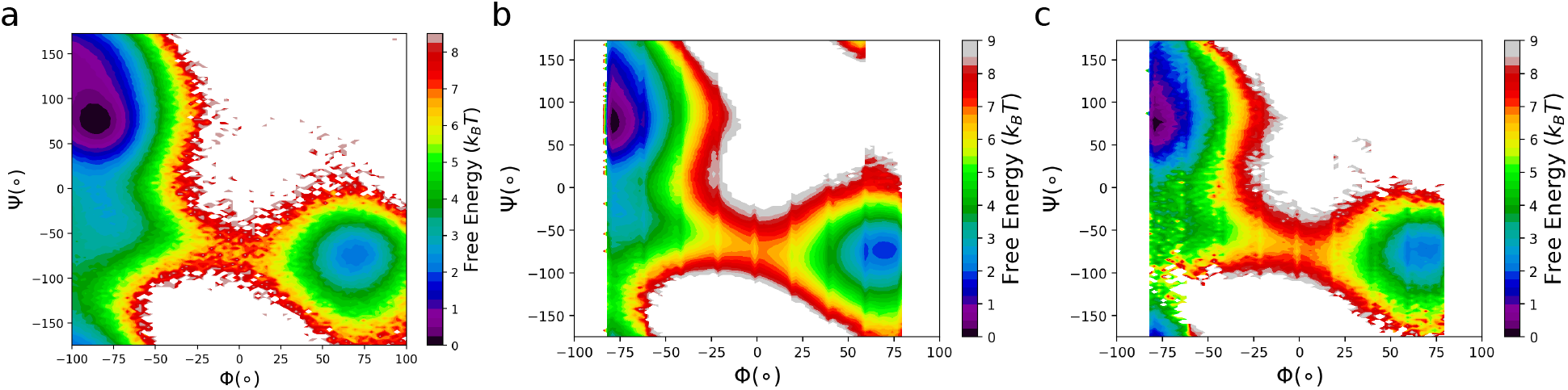
(a) Free energy landscape of gas phase alanine dipeptide obtained from equilibrium MD simulation. (b) Reconstructed free energy landscape from MMVT simulation. (c) Reconstructed free energy landscape from M-WEM simulation (trial 1). (For a better comparison the free energy landscape is constructed from M-WEM iteration 40 with approximately equal amount of total computational cost in comparison to the MMVT calculation.)

The summation is over all Λ cells and *π_α_* is the equilibrium probability of each cell.

This rescaled probability distribution is the used to reconstruct the free energy landscape (*G*(Φ, Ψ)) for the conformational transition of alanine dipeptide

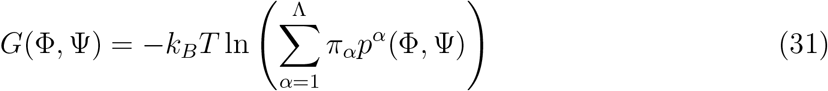

The reconstructed free energy surface for both MMVT and M-WEM simulations are in excellent agreement with one obtained from 500 ns conventional MD simulation, but confined only between the initial and final milestone i.e. −80° < Φ < 80°. This provides a way to study the free energy landscape of orthogonal degrees of freedom, which are coupled with the RC, but are not taken into consideration while devising the milestoning progress coordinate.

### Trypsin-Benzamidine Complex

Finally, we applied the M-WEM approach to calculate the kinetics and free energy for a protein ligand binding and unbinding problem. We chose the system of trypsin-benzamidine complex because of primarily two reasons. First, this system is studied extensively using MD simulations with various enhanced sampling and path sampling methods. Moreover, the residence time of the ligand is in the millisecond regime, which is beyond the reach of currently available computational power. Benzamidine is also a very potent ligand, with an experimental binding affinity (*K_d_*) of 1.2 ± 0.1 × 10^−5^ M.^81^ This is a challenging enough test system for the M-WEM method, and can also determine the utility of our approach in computer aided drug design.

The ligand residence time, unbinding rate constant (*k*_off_), binding rate constant (*k*_on_), and the binding free energy (Δ*G_b_*) have been computed from M-WEM simulation, and the results are compared with the SEEKR^50^ and MMVT SEEKR^64^ results (which used identical simulation condition as our work) and also with the experimental data^81^ (Table 3). All values obtained from M-WEM are in quantitative agreement with the experimental data. (We reported two sets of results for M-WEM, one after 200 iterations and another after 300 iterations). The *k*_off_ value, predicted from M-WEM simulation, is within the error bars of the experiment and within one order of magnitude of the SEEKR and MMVT SEEKR results. The same holds for residence time, which is the inverse of *k*_off_. Our *k*_on_ results are different from experimental value by a factor of ~ 4-5, while the results of MMVT SEEKR are approximately one order of magnitude higher. The Δ*G_b_* value computed from M-WEM, as *k_B_T* ln(*k*_off_/*k*_on_), is also in excellent agreement with the experimental number (within 1.5 kcal/mol). The error bars of the M-WEM results and the SEEKR and MMVT SEEKR are not directly comparable because the are computed differently, as described in the Theory section.

**Table 3:** Comparison of the results of the different milestoning based methods for the trypsin-benzamidine complex. (Number of iterations of M-WEM simulation are shown in parentheses.)

| Method | Residence time<br>(ms) | $k_{\text{off}}$<br>($\text{s}^{-1}$) | $k_{\text{on}}$<br>($\times 10^7 \text{ M}^{-1} \text{ s}^{-1}$) | $\Delta G_b$<br>(kcal/mol) | Simulation time<br>($\mu\text{s}$) |
| --- | --- | --- | --- | --- | --- |
| Experiment<sup>81</sup> | 1.7 | $600 \pm 300$ | 2.9 | $-6.7 \pm 0.05$ | - |
| SEEKR<sup>50</sup> | 12 | $83 \pm 14$ | $2.1 \pm 0.3$ | $-7.4 \pm 0.10$ | $\sim 19$ |
| MMVT SEEKR<sup>64</sup> | 5.6 | $174 \pm 9$ | $12 \pm 0.5$ | $-7.9 \pm 0.04$ | $\sim 4.4$ |
| MMVT SEEKR<sup>64</sup> | 16 | $62 \pm 6$ | $17 \pm 1.0$ | $-8.8 \pm 0.07$ | $\sim 2.9$ |
| M-WEM (200)<sup>a</sup> | $1.26 \pm 0.32$ | $791 \pm 197$ | $0.53 \pm 0.08$ | $-5.2 \pm 0.16$ | $\sim 0.48$ |
| M-WEM (300)<sup>a</sup> | $1.30 \pm 0.44$ | $769 \pm 261$ | $0.76 \pm 0.38$ | $-5.5 \pm 0.37$ | $\sim 0.73$ |

<sup>a</sup>Error bars are computed from the last 5 iterations sampled.

The convergence patterns of the residence time and *k*_off_ are depicted in Fig. 7. Both these values converged after about 150 iterations (~ 360 ns of total simulation time) except for small fluctuations. The *k*_off_ is computed indirectly as the inverse of residence time, which is directly obtained from M-WEM. So small fluctuations in the residence time get amplified in the *k*_off_ results in Fig. 7a. The computational cost of the M-WEM simulation is ~ 1 order of magnitude less than the other milestoning-based approaches^50, 64^ and the results are in better agreement with the experiment. Our *k*_off_ results are also closer to the experimental numbers in comparison to other methods used by Buch et al. ((9.5 ± 3.3) × 10^4^ s^−1^),^77^ Plattner et al. ((1.31 ± 1.09) × 10^4^ s^−1^),^78^ Tiwary et al. (9.1 ±2.5 s^−1^),^79^ Brotzakis et al. (4176±324 s^−1^)^101^ and Teo et al. (260±240 s^−1^);^80^ all these studies required multiple microseconds of simulation with some in the range of 50 *μ*s - 100 *μ*s.^77, 78^ A weighted ensemble-based approach has also been used to calculate the kinetics of this system by Dickson and Lotz (*k*_off_ = 5555 s^−1^)^38^ and Donyapour et al. (*k*_off_ = 266 s^−1^ and 840 s^−1^).^32^ But, unlike M-WEM, that method could only calculate the unbinding rate constant and dissociation pathways due to the use of non-equilibrium steady state. To their credit, the authors could distinguish multiple ligand release pathways, ^38^ which is difficult to achieve using milestoning-based simulations with discontinuous trajectories. Nevertheless, we tried to identify some key intermediates in the unbinding mechanism; we discuss them later in this paper.

**Figure 7:**
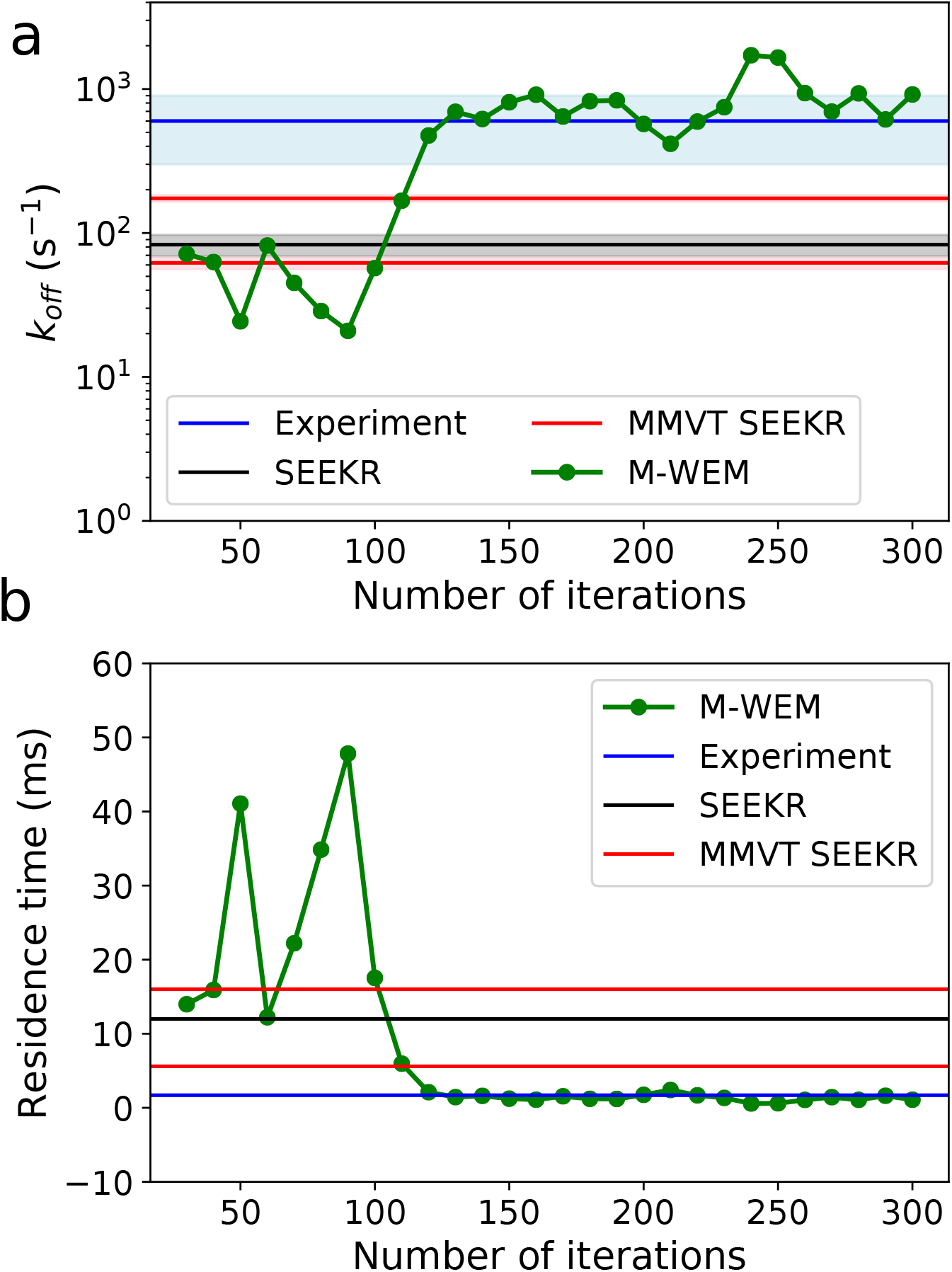
The convergence of (a) *k*_off_ and (b) ligand residence time for trypsin-benzamidine complex, as a function of M-WEM iterations. In figure (b) a linear scale is used for a better idea of the quality of the convergence.

**Figure 8:**
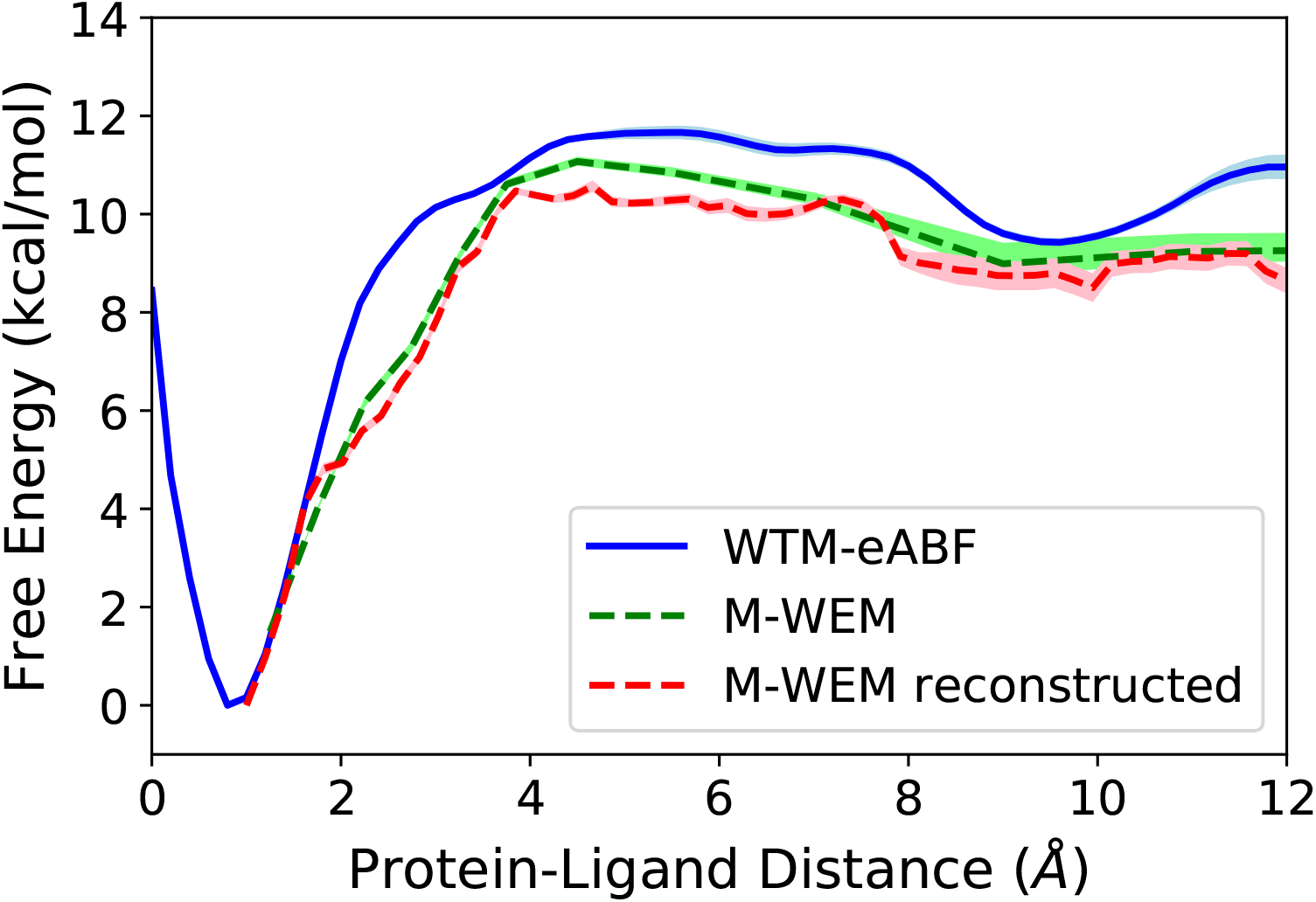
Free energy profile of the dissociation of the trypsin-benzamidine complex as a function of the milestoning reaction coordinate (the center of mass distance between the binding pocket residues and the benzamidine ligand. See Computational Methods section for details.) The results are compared between WTM-eABF simulation and the M-WEM calculation after iteration 200.

A one-dimensional free energy profile as a function of the milestoning reaction coordinate is constructed from the equilibrium probabilities (π*_α_*) obtained from the M-WEM simulation using Eq. 4. Alongside, a one dimensional free energy profile is reconstructed from the M-WEM trajectories following as described in the Theory section. Error bars in the free energy landscape are computed as the 95% confidence interval of the free energy profiles obtained between iteration 160 and 200 with an interval of 10 iterations. The two free energy profiles obtained from M-WEM using the two different techniques agree with each other and both are in reasonable agreement with the free energy surface obtained using well tempered meta-eABF (WTM-eABF) simulation^102, 103^ (see Supporting Information for details).

The committor values as a function of the milestoning reaction coordinate were computed and are indicated in Fig. 9. The results do not show much variation between 200 iterations and 300 iterations, both of which indicate that the transition state (committor = 0.5) is located between the milestones at 6 Å and 8 Å.

**Figure 9:**
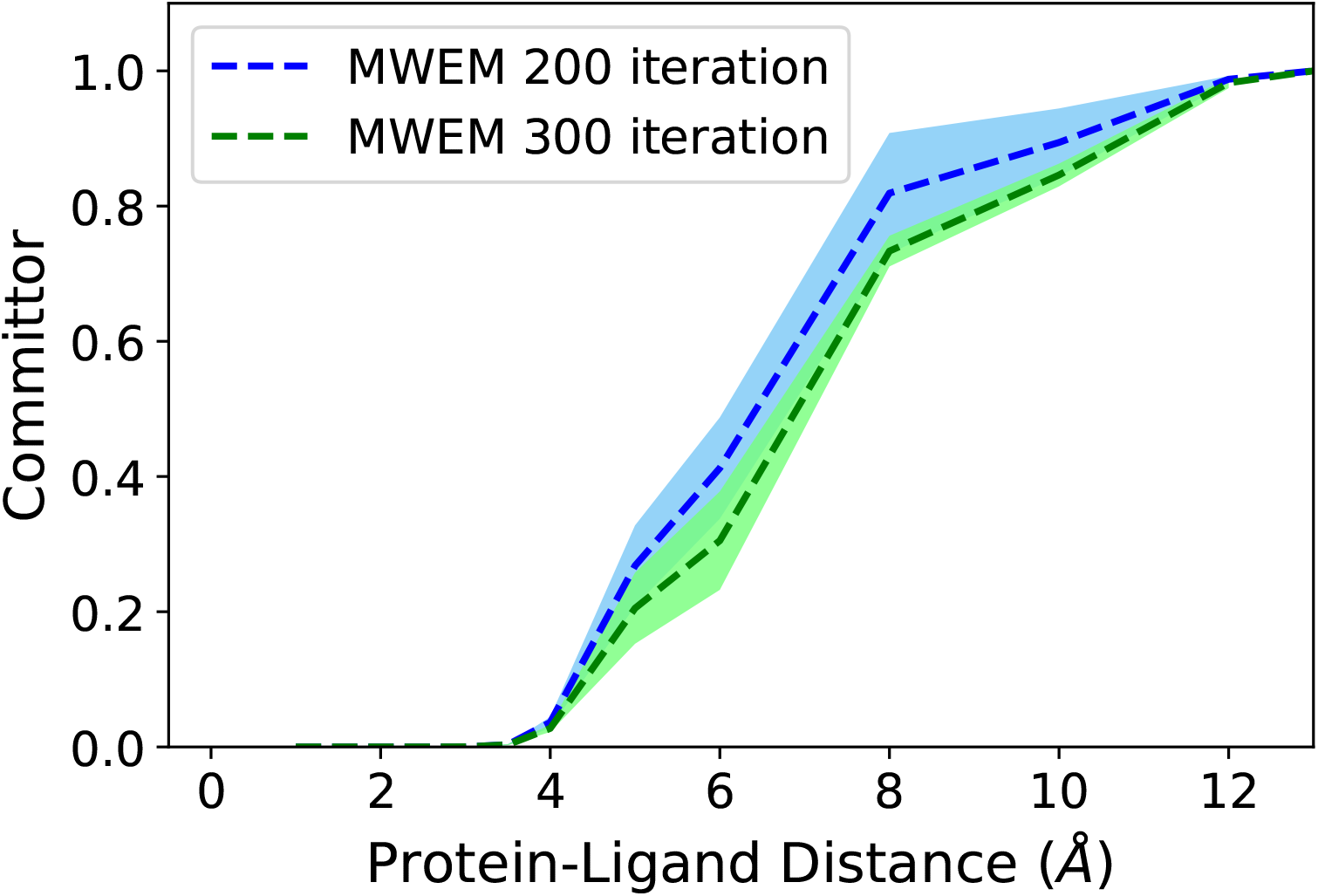
Committor values computed as a function of the milestoning reaction coordinate for the M-WEM simulation of the trypsin-benzamidine complex.

The distribution of the ligand around the protein for three cells (bound state, unbound state and the cell containing the TS) is depicted in Fig. 10. It shows the amount of threedimensional space explored by the ligand during the unbinding process. A two-dimensional projection of the ligand distribution for all cells is shown in Fig. 11. The fraction of the spherical surface covered by the ligand in the outermost cell (*α*) is used for the calculation of *k*_on_, as described in the Theory section. The increase in the exploration of the configuration space after 300 iterations in comparison to 200 iterations is small.

**Figure 10:**
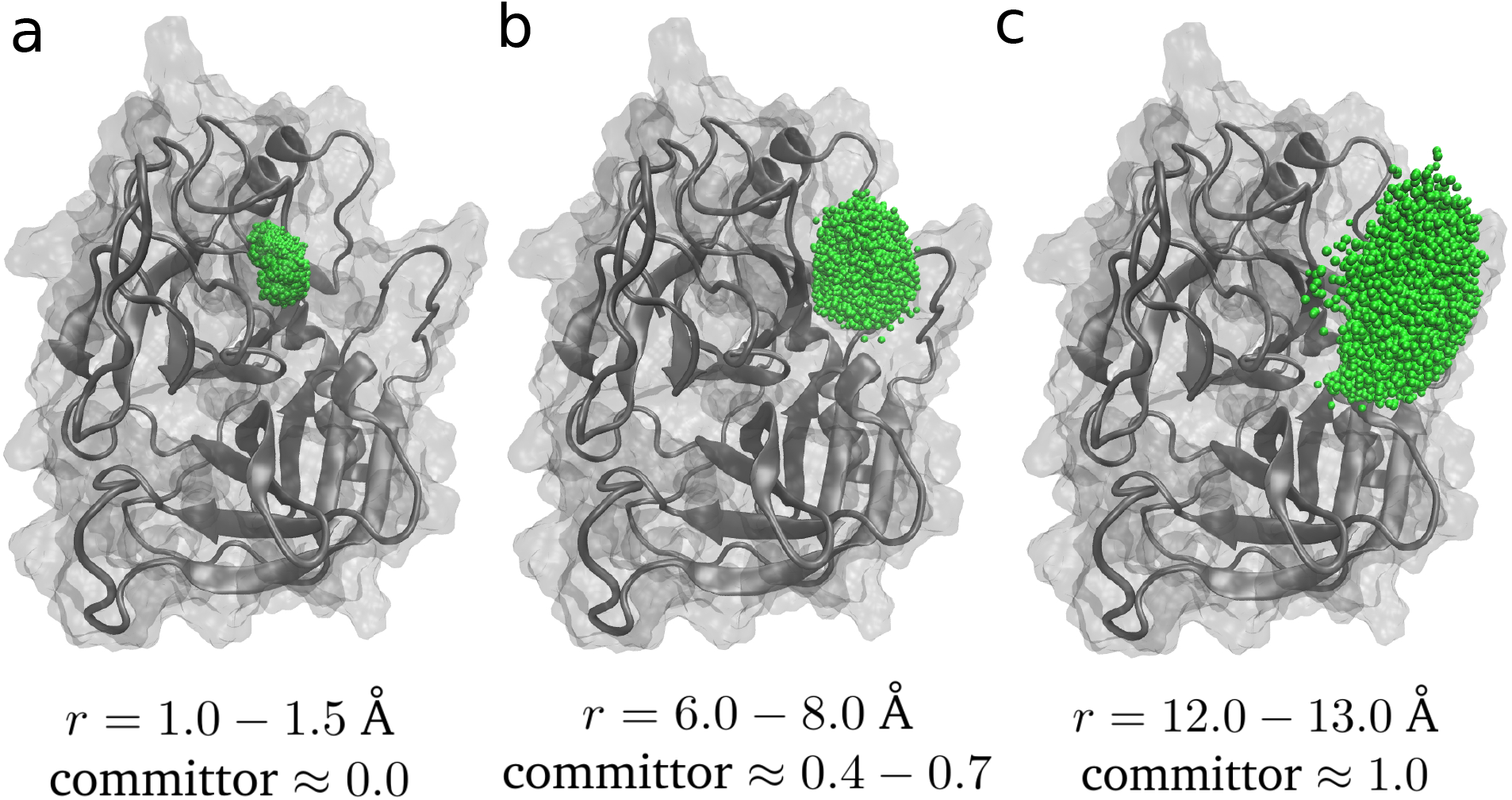
The distribution of the benzamidine ligand around the trypsin protein for three cells of the M-WEM simulations which, respectively, include: (a) the bound state, (b) an apparent transition state with committor value ~ 0.5, and (c) the unbound state.

**Figure 11:**
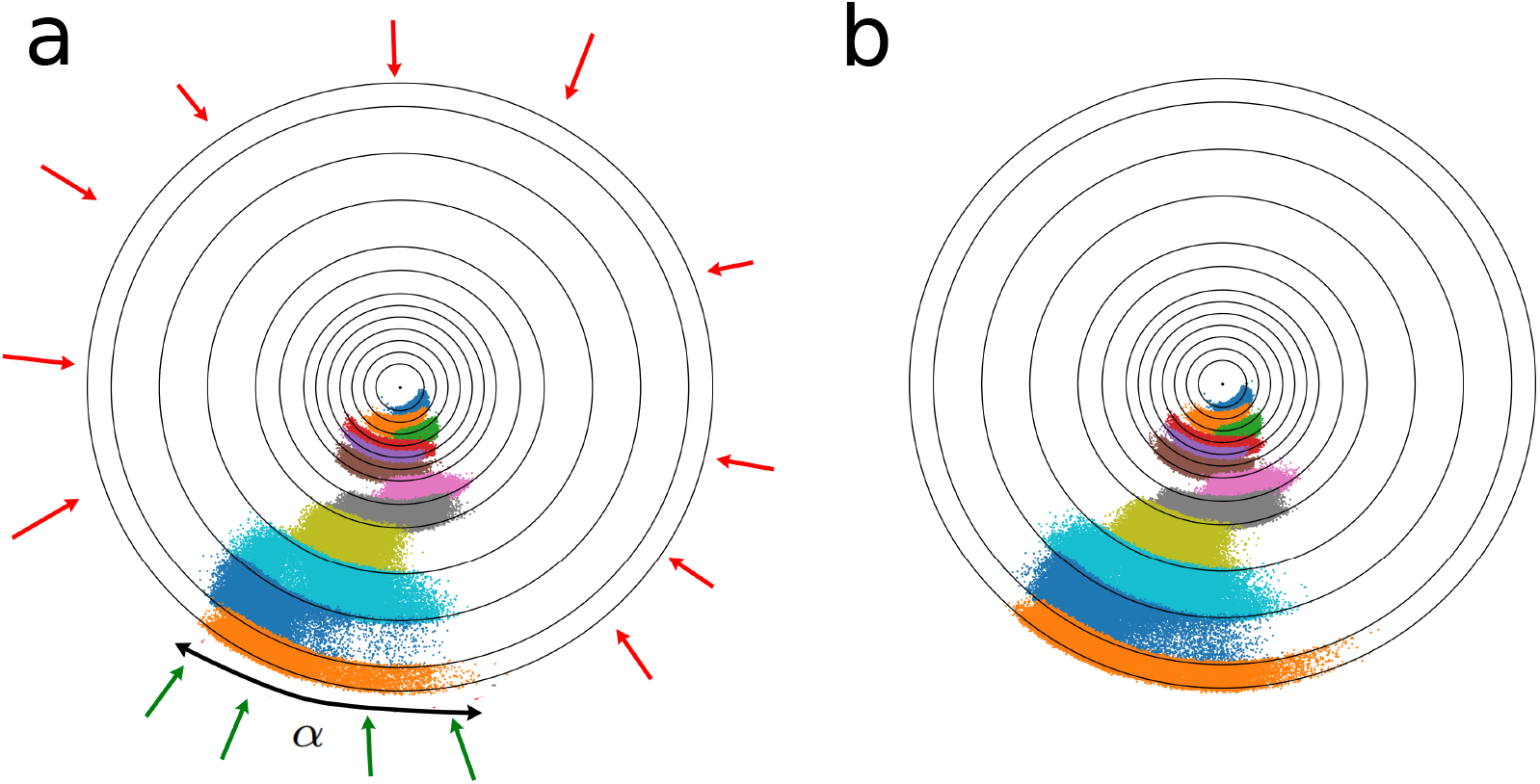
Two dimensional projection of the distribution of the ligands around the trypsin protein for (a) after M-WEM iteration 200 and (b) after iteration 300. The different colors represents structures from M-WEM simulations confined in different cells. The surface coverage *α*, used in the *k*_on_ calculation is also depicted in figure (a). For *k*_on_ calculation we assumed that the green trajectories can lead to binding events but the red trajectories cannot.

To get an idea of the intermediate states involved in the protein-ligand interaction, we clustered all the trajectory frames corresponding to each cell based on heavy-atom RMSD. The number of frames in each cell ranged between ~ 26, 000 – 30, 000, with one frame every 2 ps (the length of each WE segment). The clustering of the structures was performed using the GROMOS clustering algorithm^104^ implemented in GROMACS v2018.1^105^ with an RMSD cutoff of 0.9 Å. The cut-off was chosen such that the total number of clusters is between 10 and 40. All the cluster centers obtained from different cells were combined together and a second round of clustering is performed with an RMSD cutoff of 1.1 Å. This resulted in 14 clusters, some of which are depicted in Fig. 12. The structures are in qualitative agreement with the meta-stable states observed by Tiwary et al.^79^ and Brotzakis et al.,^101^ despite their use of a different enhanced sampling method and of a different version of the AMBER force field in the former study. Particularly, both our study and the work of Tiwary et al. show the presence of a meta-stable state in which the benzamidine is aligned in a reverse direction (the charged groups facing the aqueous environment and the hydrophobic ring facing the protein). A PDB file with all the clusters is provided in the Supporting Information.

**Figure 12:**
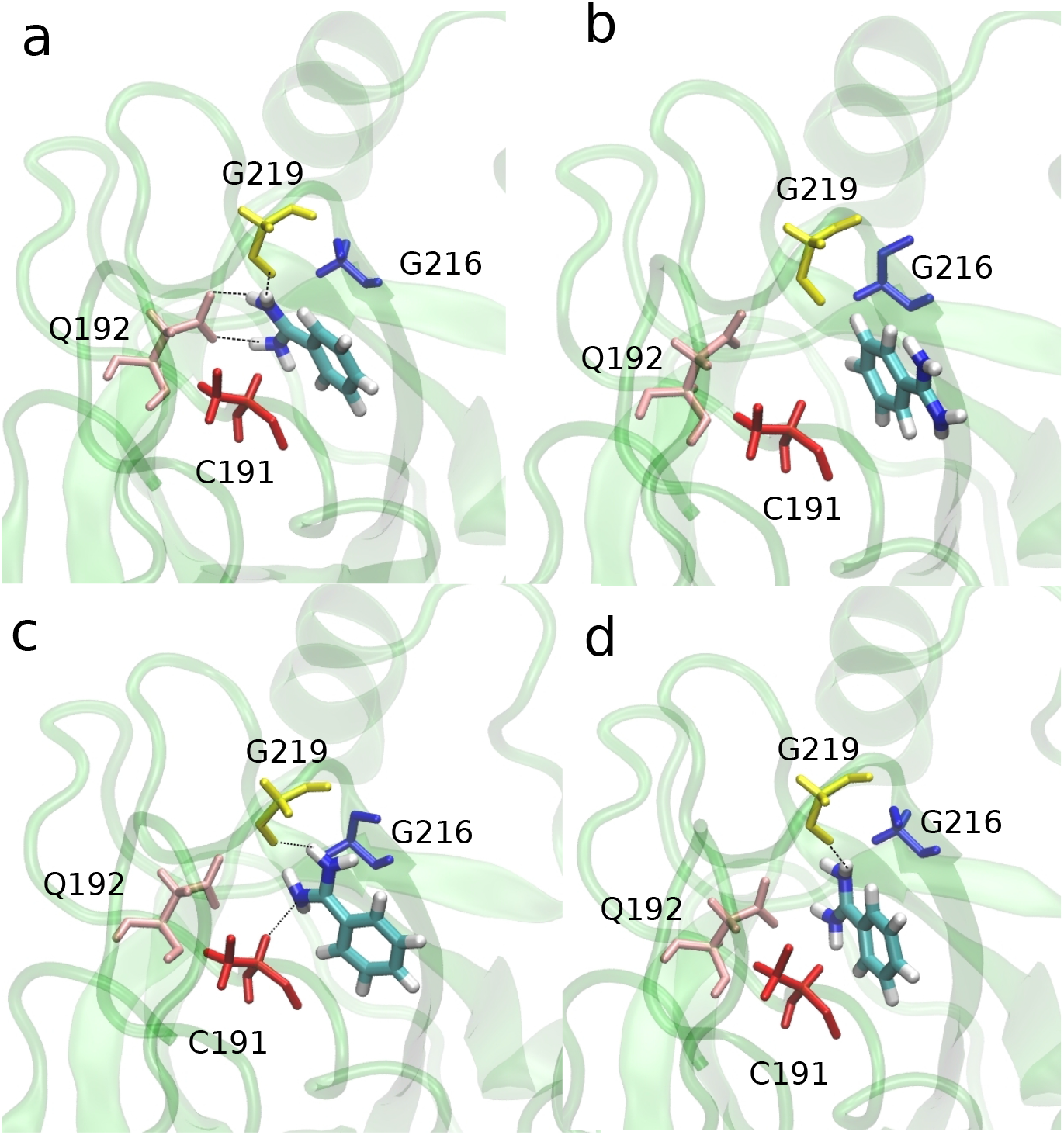
Representative structures sampled from clustering of the M-WEM trajectories of the binding/unbinding of trypsin-benzamidine complex. The ligand and the residues interacting with the ligand are shown in licorice. Hydrogen bonds between protein and ligand are shown in dashed line.

## Discussions and Conclusions

We developed a new path sampling approach which combines Markovian milestoning with a weighted ensemble scheme to efficiently calculate the kinetics and free energy of rare events using atomistic MD simulations. This method, which we call Markovian Weighted Ensemble Milestoning (M-WEM), has been applied to study the barrier crossing in a 2D toy system using the Müller-Brown potential, a conformational transition in alanine dipeptide, and, most importantly, to the dissociation and association of the trypsin-benzamidine complex, which has a millisecond scale residence time. For the Müller-Brown potential and the alanine dipeptide systems, the mean first passage time (MFPT) of conformational transition obtained from long equilibrium simulation was quantitatively reproduced by the M-WEM method at significantly lower computational cost. In the case of alanine dipeptide, we showed how one can also reproduce the two-dimensional free energy landscape as a function of two backbone torsion angles from one dimensional M-WEM and Markovian milestoning simulation, using a free energy re-scaling strategy based on the equilibrium probabilities of each milestone. This approach can be generalized to any other collective variables other than the milestoning coordinate, and can potentially elucidate the role of coupled orthogonal degrees of freedom in complex biophysical systems.

For the trypsin-benzamidine complex, the ligand residence time, *k*_off_, *k*_on_, and the binding free energy could be computed using the M-WEM method in about one order of magnitude less computational cost than the Markovian Milestoning based MMVT simulation, and 1-3 orders of magnitude less computational effort compared to other approaches previously used to study this system such as Markov state modeling, metadynamics, adaptive multilevel splitting, weighted ensemble, and traditional milestoning. Our results are in good agreement with the experimental data available for this system.

A key advantage of the M-WEM method is its simple workflow, which essentially requires the user to perform weighted ensemble simulation under flat bottom restraints. This is easy to implement in any simulation engine using an open-access weighted ensemble code such as WESTPA. We implemented M-WEM using the NAMD simulation engine and the WESTPA toolkit. Our implementation uses a minimal adaptive binning (MAB) scheme, which allows for the adaptation of the WE bins throughout the simulation to increase sampling in high energy regions. Consequently, it does not require preexisting knowledge of the energy landscape and can efficiently sample all possible transitions between milestone interfaces. Moreover, in contrast to traditional milestoning approaches, M-WEM (or Markovian milestoning in general) it does not require additional simulation (e.g. umbrella sampling) along the milestone interface to sample starting structures, a process which accounts for the majority of the total computational effort. In our previous work, we attempted to replace this expensive additional step using a weighted ensemble restrain-and-release scheme. ^76^ The Markovian milestoning technique completely removes this step as the trajectory, confined between two milestones, explores by itself the configurational space orthogonal to the milestoning coordinate. In the M-WEM approach, we accelerated this “orthogonal sampling”, by using 2D WE bins along two progress coordinates: the milestoning reaction coordinate (to accelerate milestone-to-milestone transitions) and also in another coordinate along the milestone interface. Due to dimensionality scaling, the advantage of the M-WEM over traditional Markovian milestoning is more pronounced in the case of trypsin-benzamidine complex, where results, in better agreement with the experiment, could be obtained using M-WEM simulation within a fraction of the computational cost of MMVT SEEKR calculations on the same system. Also, the M-WEM protocol does not need to stop the trajectory at milestone interfaces, avoiding frequent intervention to the dynamics engine, and therefore making it more efficient to implement in GPU-based hardware.

Apart from these unique achievements, M-WEM also shares some common advantages with our previously-developed WEM methodology. They include the possibility of massively parallelizing the simulations over each milestone, which will be even more pronounced in the current implementation, as MAB binning has been shown to utilize GPU-based hardware more efficiently than the traditional fixed-binning scheme we used in our earlier work. The convergence of the transition statistics in-between milestones is also quicker in M-WEM in comparison to MMVT, as evident from the results for the trypsin-benzamidine complex. We also show that a relatively crude reaction coordinate is capable of producing accurate kinetics, particularly in the cases of the Müller-Brown potential and the alanine dipeptide model.

The M-WEM approach, being a combination of two fairly complex path sampling methods, inherits all the assumption from each of the individual techniques. Similar to MMVT, M-WEM also requires the milestones to be sufficiently far apart so that the transitions between them are independent of the transitions from other milestones.^42, 83^ It also assumes a complete exploration of the configurational space in the milestone hypersurface. Although the weighted ensemble approach is invoked to satisfy these assumptions at a lower computational cost, it comes with additional assumptions inherited from the WE scheme. For example, each individual trajectory trace contributes independently towards the transition statistics despite sharing a significant portion of its propagation history.^20^ Additionally, due to the use of a stochastic integrator the timscales obtained from the M-WEM simulations can be dependent on the damping coefficent (*γ*) of the Langevin dynamics.^24, 106^ However, Hall et al. have shown that *γ* has little effect on ligand residence times obtained from explicit solvent simulation as long as *γ* > 0.1 ps^−1^.^106^ Our choice of *γ* = 5 ps^−1^ was motivated by previous milestoning and MMVT work on trypsin-benzamidine,^50, 64^ and also by the fact that a *γ* of 5 ps^−1^ reproduces the experimental diffusion constant of water most accurately. ^107^

A recent study using the MMVT SEEKR approach has predicted the *k*_off_ of trypsin-benzamidine complex to be 990±70 s^−1^, ^108^ a value that is relatively closer to the experimental number compared to the previous result of 62±6 s^−1^ (Table 3). However, the fact that a virtually identical simulation scheme can produce results that are more than one order of magnitude apart clearly shows the high level of uncertainty involved in the milestoning approach. So, despite having a better agreement with the experimental value compared to some previous studies, the order-of-magnitude agreement of *k*_off_ remains the key achievement of the M-WEM approach.

One of the limitations of the current implementation of M-WEM is the use of an analytical approach to compute the binding rate constant. The alternative is to use a multiscale Brownian dynamics (BD) approach,^50, 56, 64^ which is more rigorous but more computationally expensive. However, BD methods allow us to include the effect of position-dependent variation of the diffusion constant, as well as of the ionic strength of the solution, both of which are absent in our current implementation.

Our M-WEM method can find application in studying the kinetics and free energy of biomolecular rare events not only for the purpose of fundamental understanding of biological processes, but also for kinetics-driven computer aided drug design. Evidence has emerged over the past decade showing that the efficacy of a small molecule therapeutic drug is more correlated with the residence time than with the binding affinity.^8, 9^ Yet, the majority of the drug design effort in the pharmaceutical industry is based on binding free energy; among other things, this is because it is easier to compute than kinetics. The M-WEM approach is a cheap alternative to computationally expensive traditional enhanced sampling and to path sampling methods, and can be included in a computational drug design pipeline using both binding free energy and kinetics. In the future, we plan to test this method on protein-ligand systems with longer residence time, e.g.,in the range of minutes to hours, a time frame more characteristic of the drug molecules used in practical application. The increased sampling of orthogonal coordinates in M-WEM can also facilitate the study of systems where a protein conformational change is coupled to a ligand-binding coordinate. Overall, our novel Markovian Weighted Ensemble Milestoning approach is expected to be successful in predicting the free energy and kinetics of biophysical rare events with quantitative accuracy, and it holds the potential of becoming a useful tool in the large-scale computational screening of therapeutic drugs.

## Supporting information

MWEM-main.zip

clusters.pdb

## Acknowledgement

The authors thank Rommie Amaro and Lane Votapka for kindly sharing the structure and topology files of the trypsin-benzamidine complex. The authors thank Trevor Gokey for providing scripts for protein structure visualization and analysis. This work was supported partially by the National Science Foundation (NSF) via grant MCB 2028443. SES acknowledges support through Pfizer - La Jolla Academic Industrial Relations (AIR) Diversity Research Fellowship in Chemistry. The authors thank the University of California Irvine High Performance Computing (HPC) facility for providing the computational resources. The authors declare no competing financial interest.

## Data Availability Statement

The implementation of the M-WEM method using WESTPA software and NAMD simulation package is available from the github repository: https://github.com/dhimanray/MWEM. Additional scripts for input file preparation, data analysis and visualization are provided in the same repository. A compressed zip file of the codes is provided in the supporting information. All the rest of the data are provided in the manuscript and in the supporting information. Additional data are available from the corresponding author upon request.

## Supporting Information Available

### Supporting Information Text

Discussion on the convergence of M-WEM simulations for Alanine dipeptide system, the computational details for the meta-eABF simulation of the trypsin-benzamidine complex, and discussion on the appropriate placement of milestones are provided in the supporting information. Fig. S1-S3 are included in the supporting information.

### clusters.pdb

A pdb file containing 14 clusters for the trypsin-benzamidine complex

### MWEM-main.zip

A compressed folder containing the codes for M-WEM implementation.

## Graphical TOC Entry

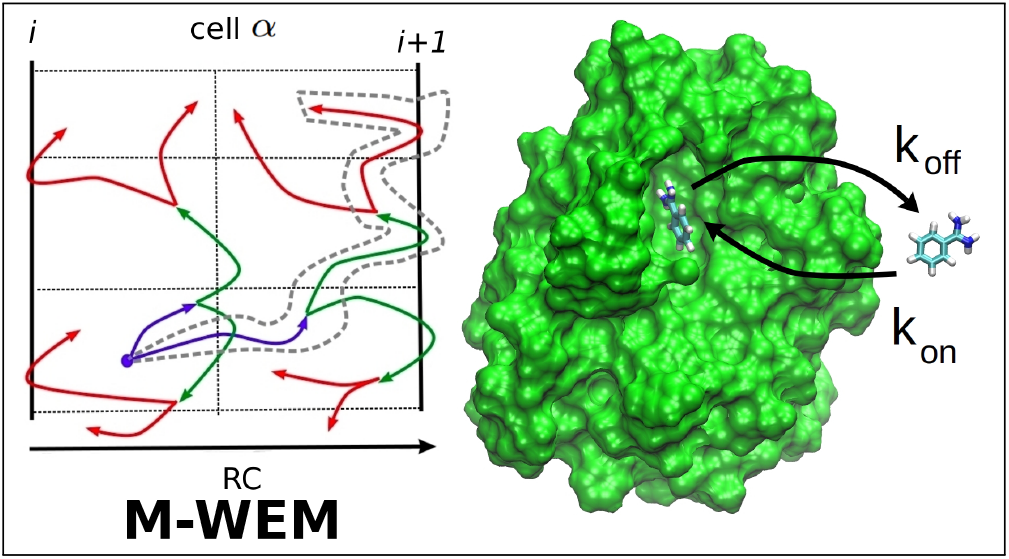

## Notes

https://github.com/dhimanray/MWEM

### Competing Interest Statement

The authors have declared no competing interest.

