## Supplementary material for "Markovian Weighted Ensemble Milestoning (M-WEM): Long-time Kinetics from Short Trajectories": MWEM-main.zip: CONTRIBUTING.md

# How to contribute

We welcome contributions from external contributors, and this document
describes how to merge code changes into this mwem. 

## Getting Started

* Make sure you have a [GitHub account](https://github.com/signup/free).
* [Fork](https://help.github.com/articles/fork-a-repo/) this repository on GitHub.
* On your local machine,
  [clone](https://help.github.com/articles/cloning-a-repository/) your fork of
  the repository.

## Making Changes

* Add some really awesome code to your local fork.  It's usually a [good
  idea](http://blog.jasonmeridth.com/posts/do-not-issue-pull-requests-from-your-master-branch/)
  to make changes on a
  [branch](https://help.github.com/articles/creating-and-deleting-branches-within-your-repository/)
  with the branch name relating to the feature you are going to add.
* When you are ready for others to examine and comment on your new feature,
  navigate to your fork of mwem on GitHub and open a [pull
  request](https://help.github.com/articles/using-pull-requests/) (PR). Note that
  after you launch a PR from one of your fork's branches, all
  subsequent commits to that branch will be added to the open pull request
  automatically.  Each commit added to the PR will be validated for
  mergability, compilation and test suite compliance; the results of these tests
  will be visible on the PR page.
* If you're providing a new feature, you must add test cases and documentation.
* When the code is ready to go, make sure you run the test suite using pytest.
* When you're ready to be considered for merging, check the "Ready to go"
  box on the PR page to let the mwem devs know that the changes are complete.
  The code will not be merged until this box is checked, the continuous
  integration returns checkmarks,
  and multiple core developers give "Approved" reviews.

# Additional Resources

* [General GitHub documentation](https://help.github.com/)
* [PR best practices](http://codeinthehole.com/writing/pull-requests-and-other-good-practices-for-teams-using-github/)
* [A guide to contributing to software packages](http://www.contribution-guide.org)
* [Thinkful PR example](http://www.thinkful.com/learn/github-pull-request-tutorial/#Time-to-Submit-Your-First-PR)
