## Supplementary material for "Markovian Weighted Ensemble Milestoning (M-WEM): Long-time Kinetics from Short Trajectories": MWEM-main.zip: PULL_REQUEST_TEMPLATE.md

## Description
Provide a brief description of the PR's purpose here.

## Todos
Notable points that this PR has either accomplished or will accomplish.
  - [ ] TODO 1

## Questions
- [ ] Question1

## Status
- [ ] Ready to go
