## Supplementary material for "Markovian Weighted Ensemble Milestoning (M-WEM): Long-time Kinetics from Short Trajectories": MWEM-main.zip: CI.yaml

```
name: CI

on:
  # GitHub has started calling new repo's first branch "main" https://github.com/github/renaming
  # Existing codes likely still have "master" as the primary branch
  # Both are tracked here to keep legacy and new codes working
  push:
    branches:
      - "master"
      - "main"
  pull_request:
    branches:
      - "master"
      - "main"
  schedule:
    # Nightly tests run on master by default:
    #   Scheduled workflows run on the latest commit on the default or base branch.
    #   (from https://help.github.com/en/actions/reference/events-that-trigger-workflows#scheduled-events-schedule)
    - cron: "0 0 * * *"

jobs:
  test:
    name: Test on ${{ matrix.os }}, Python ${{ matrix.python-version }}
    runs-on: ${{ matrix.os }}
    strategy:
      matrix:
        os: [macOS-latest, ubuntu-latest, windows-latest]
        python-version: [3.7, 3.8, 3.9]

    steps:
    - uses: actions/checkout@v1

    - name: Additional info about the build
      shell: bash
      run: |
        uname -a
        df -h
        ulimit -a


    # More info on options: https://github.com/conda-incubator/setup-miniconda
    - uses: conda-incubator/setup-miniconda@v2
      with:
        python-version: ${{ matrix.python-version }}
        environment-file: devtools/conda-envs/test_env.yaml

        channels: conda-forge,defaults

        activate-environment: test
        auto-update-conda: false
        auto-activate-base: false
        show-channel-urls: true

    - name: Install package

      # conda setup requires this special shell
      shell: bash -l {0}
      run: |
        python -m pip install . --no-deps
        conda list


    - name: Run tests

      # conda setup requires this special shell
      shell: bash -l {0}

      run: |
        pytest -v --cov=mwem --cov-report=xml --color=yes mwem/tests/

    - name: CodeCov
      uses: codecov/codecov-action@v1
      with:
        file: ./coverage.xml
        flags: unittests
        name: codecov-${{ matrix.os }}-py${{ matrix.python-version }}
```
