## Supplementary material for "Markovian Weighted Ensemble Milestoning (M-WEM): Long-time Kinetics from Short Trajectories": MWEM-main.zip: CODE_OF_CONDUCT.md

# Contributor Covenant Code of Conduct

## Our Pledge

In the interest of fostering an open and welcoming environment, we as
contributors and maintainers pledge to making participation in our project and
our community a harassment-free experience for everyone, regardless of age,
body size, disability, ethnicity, gender identity and expression, level of
experience, nationality, personal appearance, race, religion, or sexual
identity and orientation.

## Our Standards

Examples of behavior that contributes to creating a positive environment include:

* Using welcoming and inclusive language
* Being respectful of differing viewpoints and experiences
* Gracefully accepting constructive criticism
* Focusing on what is best for the community
* Showing empathy towards other community members

Examples of unacceptable behavior by participants include:

* The use of sexualized language or imagery and unwelcome sexual attention or advances
* Trolling, insulting/derogatory comments, and personal or political attacks
* Public or private harassment
* Publishing others' private information, such as a physical or electronic address, without explicit permission
* Other conduct which could reasonably be considered inappropriate in a professional setting

## Our Responsibilities

Project maintainers are responsible for clarifying the standards of acceptable
behavior and are expected to take appropriate and fair corrective action in
response to any instances of unacceptable behavior.

Project maintainers have the right and responsibility to remove, edit, or
reject comments, commits, code, wiki edits, issues, and other contributions
that are not aligned to this Code of Conduct, or to ban temporarily or
permanently any contributor for other behaviors that they deem inappropriate,
threatening, offensive, or harmful.

Moreover, project maintainers will strive to offer feedback and advice to
ensure quality and consistency of contributions to the code.  Contributions
from outside the group of project maintainers are strongly welcomed but the
final decision as to whether commits are merged into the codebase rests with
the team of project maintainers.

## Scope

This Code of Conduct applies both within project spaces and in public spaces
when an individual is representing the project or its community. Examples of
representing a project or community include using an official project e-mail
address, posting via an official social media account, or acting as an
appointed representative at an online or offline event. Representation of a
project may be further defined and clarified by project maintainers.

## Enforcement

Instances of abusive, harassing, or otherwise unacceptable behavior may be
reported by contacting the project team at ''. The project team will
review and investigate all complaints, and will respond in a way that it deems
appropriate to the circumstances. The project team is obligated to maintain
confidentiality with regard to the reporter of an incident. Further details of
specific enforcement policies may be posted separately.

Project maintainers who do not follow or enforce the Code of Conduct in good
faith may face temporary or permanent repercussions as determined by other
members of the project's leadership.

## Attribution

This Code of Conduct is adapted from the [Contributor Covenant][homepage],
version 1.4, available at
[http://contributor-covenant.org/version/1/4][version]

[homepage]: http://contributor-covenant.org
[version]: http://contributor-covenant.org/version/1/4/
