## Supplementary material for "Markovian Weighted Ensemble Milestoning (M-WEM): Long-time Kinetics from Short Trajectories": MWEM-main.zip: test_env.yaml

```
name: test
channels:

  - conda-forge

  - defaults
dependencies:
    # Base depends
  - python
  - pip

    # Testing
  - pytest
  - pytest-cov
  - codecov

    # Pip-only installs
  #- pip:
  #  - codecov
```
