## Supplementary material for "Markovian Weighted Ensemble Milestoning (M-WEM): Long-time Kinetics from Short Trajectories": MWEM-main.zip: before_install.sh

```
# Temporarily change directory to $HOME to install software
pushd .
cd $HOME
# Make sure some level of pip is installed
python -m ensurepip

# Install Miniconda
if [ "$TRAVIS_OS_NAME" == "osx" ]; then
    # Make OSX md5 mimic md5sum from linux, alias does not work
    md5sum () {
        command md5 -r "$@"
    }
    MINICONDA=Miniconda3-latest-MacOSX-x86_64.sh
else
    MINICONDA=Miniconda3-latest-Linux-x86_64.sh
fi
MINICONDA_HOME=$HOME/miniconda
MINICONDA_MD5=$(wget -qO- https://repo.anaconda.com/miniconda/ | grep -A3 $MINICONDA | sed -n '4p' | sed -n 's/ *<td>\(.*\)<\/td> */\1/p')
wget -q https://repo.anaconda.com/miniconda/$MINICONDA
if [[ $MINICONDA_MD5 != $(md5sum $MINICONDA | cut -d ' ' -f 1) ]]; then
    echo "Miniconda MD5 mismatch"
    exit 1
fi
bash $MINICONDA -b -p $MINICONDA_HOME

# Configure miniconda
export PIP_ARGS="-U"
# New to conda >=4.4
echo ". $MINICONDA_HOME/etc/profile.d/conda.sh" >> ~/.bashrc  # Source the profile.d file
echo "conda activate" >> ~/.bashrc  # Activate conda
source ~/.bashrc  # source file to get new commands
#export PATH=$MINICONDA_HOME/bin:$PATH  # Old way, should not be needed anymore
    
conda config --add channels conda-forge
    
conda config --set always_yes yes
conda install conda conda-build jinja2 anaconda-client
conda update --quiet --all

# Restore original directory
popd
```
