## Supplementary material for "Markovian Weighted Ensemble Milestoning (M-WEM): Long-time Kinetics from Short Trajectories": MWEM-main.zip: README.md

# Development, testing, and deployment tools

This directory contains a collection of tools for running Continuous Integration (CI) tests, 
conda installation, and other development tools not directly related to the coding process.


## Manifest

### Continuous Integration

You should test your code, but do not feel compelled to use these specific programs. You also may not need Unix and 
Windows testing if you only plan to deploy on specific platforms. These are just to help you get started.

The items in this directory have been left for legacy purposes since the change to GitHub Actions, 
They will likely be removed in a future version.

* `legacy-miniconda-setup`: A preserved copy of a helper directory which made Linux and OSX based testing through [Travis-CI](https://about.travis-ci.com/) simpler
  * `before_install.sh`: Pip/Miniconda pre-package installation script for Travis. No longer needed thanks to 
    [GitHub Actions](https://docs.github.com/en/free-pro-team@latest/actions) and the [conda-incubator/setup-miniconda Action](https://github.com/conda-incubator/setup-miniconda)

### Conda Environment:

This directory contains the files to setup the Conda environment for testing purposes

* `conda-envs`: directory containing the YAML file(s) which fully describe Conda Environments, their dependencies, and those dependency provenance's
  * `test_env.yaml`: Simple test environment file with base dependencies. Channels are not specified here and therefore respect global Conda configuration
  
### Additional Scripts:

This directory contains OS agnostic helper scripts which don't fall in any of the previous categories
* `scripts`
  * `create_conda_env.py`: Helper program for spinning up new conda environments based on a starter file with Python Version and Env. Name command-line options


## How to contribute changes
- Clone the repository if you have write access to the main repo, fork the repository if you are a collaborator.
- Make a new branch with `git checkout -b {your branch name}`
- Make changes and test your code
- Ensure that the test environment dependencies (`conda-envs`) line up with the build and deploy dependencies (`conda-recipe/meta.yaml`)
- Push the branch to the repo (either the main or your fork) with `git push -u origin {your branch name}`
  * Note that `origin` is the default name assigned to the remote, yours may be different
- Make a PR on GitHub with your changes
- We'll review the changes and get your code into the repo after lively discussion!


## Checklist for updates
- [ ] Make sure there is an/are issue(s) opened for your specific update
- [ ] Create the PR, referencing the issue
- [ ] Debug the PR as needed until tests pass
- [ ] Tag the final, debugged version 
   *  `git tag -a X.Y.Z [latest pushed commit] && git push --follow-tags`
- [ ] Get the PR merged in

## Versioneer Auto-version
[Versioneer](https://github.com/warner/python-versioneer) will automatically infer what version 
is installed by looking at the `git` tags and how many commits ahead this version is. The format follows 
[PEP 440](https://www.python.org/dev/peps/pep-0440/) and has the regular expression of:
```regexp
\d+.\d+.\d+(?\+\d+-[a-z0-9]+)
```
If the version of this commit is the same as a `git` tag, the installed version is the same as the tag, 
e.g. `mwem-0.1.2`, otherwise it will be appended with `+X` where `X` is the number of commits 
ahead from the last tag, and then `-YYYYYY` where the `Y`'s are replaced with the `git` commit hash.
