## Supplementary material for "Markovian Weighted Ensemble Milestoning (M-WEM): Long-time Kinetics from Short Trajectories": MWEM-main.zip: create_conda_env.py

```
import argparse
import os
import re
import glob
import shutil
import subprocess as sp
from tempfile import TemporaryDirectory
from contextlib import contextmanager
# YAML imports
try:
    import yaml  # PyYAML
    loader = yaml.load
except ImportError:
    try:
        import ruamel_yaml as yaml  # Ruamel YAML
    except ImportError:
        try:
            # Load Ruamel YAML from the base conda environment
            from importlib import util as import_util
            CONDA_BIN = os.path.dirname(os.environ['CONDA_EXE'])
            ruamel_yaml_path = glob.glob(os.path.join(CONDA_BIN, '..',
                                                      'lib', 'python*.*', 'site-packages',
                                                      'ruamel_yaml', '__init__.py'))[0]
            # Based on importlib example, but only needs to load_module since its the whole package, not just
            # a module
            spec = import_util.spec_from_file_location('ruamel_yaml', ruamel_yaml_path)
            yaml = spec.loader.load_module()
        except (KeyError, ImportError, IndexError):
            raise ImportError("No YAML parser could be found in this or the conda environment. "
                              "Could not find PyYAML or Ruamel YAML in the current environment, "
                              "AND could not find Ruamel YAML in the base conda environment through CONDA_EXE path. " 
                              "Environment not created!")
    loader = yaml.YAML(typ="safe").load  # typ="safe" avoids odd typing on output


@contextmanager
def temp_cd():
    """Temporary CD Helper"""
    cwd = os.getcwd()
    with TemporaryDirectory() as td:
        try:
            os.chdir(td)
            yield
        finally:
            os.chdir(cwd)


# Args
parser = argparse.ArgumentParser(description='Creates a conda environment from file for a given Python version.')
parser.add_argument('-n', '--name', type=str,
                    help='The name of the created Python environment')
parser.add_argument('-p', '--python', type=str,
                    help='The version of the created Python environment')
parser.add_argument('conda_file',
                    help='The file for the created Python environment')

args = parser.parse_args()

# Open the base file
with open(args.conda_file, "r") as handle:
    yaml_script = loader(handle.read())

python_replacement_string = "python {}*".format(args.python)

try:
    for dep_index, dep_value in enumerate(yaml_script['dependencies']):
        if re.match('python([ ><=*]+[0-9.*]*)?$', dep_value):  # Match explicitly 'python' and its formats
            yaml_script['dependencies'].pop(dep_index)
            break  # Making the assumption there is only one Python entry, also avoids need to enumerate in reverse
except (KeyError, TypeError):
    # Case of no dependencies key, or dependencies: None
    yaml_script['dependencies'] = []
finally:
    # Ensure the python version is added in. Even if the code does not need it, we assume the env does
    yaml_script['dependencies'].insert(0, python_replacement_string)

# Figure out conda path
if "CONDA_EXE" in os.environ:
    conda_path = os.environ["CONDA_EXE"]
else:
    conda_path = shutil.which("conda")
if conda_path is None:
    raise RuntimeError("Could not find a conda binary in CONDA_EXE variable or in executable search path")

print("CONDA ENV NAME  {}".format(args.name))
print("PYTHON VERSION  {}".format(args.python))
print("CONDA FILE NAME {}".format(args.conda_file))
print("CONDA PATH      {}".format(conda_path))

# Write to a temp directory which will always be cleaned up
with temp_cd():
    temp_file_name = "temp_script.yaml"
    with open(temp_file_name, 'w') as f:
        f.write(yaml.dump(yaml_script))
    sp.call("{} env create -n {} -f {}".format(conda_path, args.name, temp_file_name), shell=True)
```
