## Supplementary material for "Markovian Weighted Ensemble Milestoning (M-WEM): Long-time Kinetics from Short Trajectories": MWEM-main.zip: README.md

# Static Doc Directory

Add any paths that contain custom static files (such as style sheets) here,
relative to the `conf.py` file's directory. 
They are copied after the builtin static files,
so a file named "default.css" will overwrite the builtin "default.css".

The path to this folder is set in the Sphinx `conf.py` file in the line: 
```python
templates_path = ['_static']
```

## Examples of file to add to this directory
* Custom Cascading Style Sheets
* Custom JavaScript code
* Static logo images
