## Supplementary material for "Markovian Weighted Ensemble Milestoning (M-WEM): Long-time Kinetics from Short Trajectories": MWEM-main.zip: README.md

# Templates Doc Directory

Add any paths that contain templates here, relative to  
the `conf.py` file's directory.
They are copied after the builtin template files,
so a file named "page.html" will overwrite the builtin "page.html".

The path to this folder is set in the Sphinx `conf.py` file in the line: 
```python
html_static_path = ['_templates']
```

## Examples of file to add to this directory
* HTML extensions of stock pages like `page.html` or `layout.html`
