## Supplementary material for "Markovian Weighted Ensemble Milestoning (M-WEM): Long-time Kinetics from Short Trajectories": MWEM-main.zip: conf.py

```
# -*- coding: utf-8 -*-
#
# Configuration file for the Sphinx documentation builder.
#
# This file does only contain a selection of the most common options. For a
# full list see the documentation:
# http://www.sphinx-doc.org/en/stable/config

# -- Path setup --------------------------------------------------------------

# If extensions (or modules to document with autodoc) are in another directory,
# add these directories to sys.path here. If the directory is relative to the
# documentation root, use os.path.abspath to make it absolute, like shown here.

# Incase the project was not installed
import os
import sys
sys.path.insert(0, os.path.abspath('..'))

import mwem


# -- Project information -----------------------------------------------------

project = 'MWEM'
copyright = ("2021, Dhiman Ray. Project structure based on the "
             "Computational Molecular Science Python Cookiecutter version 1.5")
author = 'Dhiman Ray'

# The short X.Y version
version = ''
# The full version, including alpha/beta/rc tags
release = ''


# -- General configuration ---------------------------------------------------

# If your documentation needs a minimal Sphinx version, state it here.
#
# needs_sphinx = '1.0'

# Add any Sphinx extension module names here, as strings. They can be
# extensions coming with Sphinx (named 'sphinx.ext.*') or your custom
# ones.
extensions = [
    'sphinx.ext.autosummary',
    'sphinx.ext.autodoc',
    'sphinx.ext.mathjax',
    'sphinx.ext.viewcode',
    'sphinx.ext.napoleon',
    'sphinx.ext.intersphinx',
    'sphinx.ext.extlinks',
]

autosummary_generate = True
napoleon_google_docstring = False
napoleon_use_param = False
napoleon_use_ivar = True

# Add any paths that contain templates here, relative to this directory.
templates_path = ['_templates']

# The suffix(es) of source filenames.
# You can specify multiple suffix as a list of string:
#
# source_suffix = ['.rst', '.md']
source_suffix = '.rst'

# The master toctree document.
master_doc = 'index'

# The language for content autogenerated by Sphinx. Refer to documentation
# for a list of supported languages.
#
# This is also used if you do content translation via gettext catalogs.
# Usually you set "language" from the command line for these cases.
language = None

# List of patterns, relative to source directory, that match files and
# directories to ignore when looking for source files.
# This pattern also affects html_static_path and html_extra_path .
exclude_patterns = ['_build', 'Thumbs.db', '.DS_Store']

# The name of the Pygments (syntax highlighting) style to use.
pygments_style = 'default'


# -- Options for HTML output -------------------------------------------------

# The theme to use for HTML and HTML Help pages.  See the documentation for
# a list of builtin themes.
#
html_theme = 'sphinx_rtd_theme'

# Theme options are theme-specific and customize the look and feel of a theme
# further.  For a list of options available for each theme, see the
# documentation.
#
# html_theme_options = {}

# Add any paths that contain custom static files (such as style sheets) here,
# relative to this directory. They are copied after the builtin static files,
# so a file named "default.css" will overwrite the builtin "default.css".
html_static_path = ['_static']

# Custom sidebar templates, must be a dictionary that maps document names
# to template names.
#
# The default sidebars (for documents that don't match any pattern) are
# defined by theme itself.  Builtin themes are using these templates by
# default: ``['localtoc.html', 'relations.html', 'sourcelink.html',
# 'searchbox.html']``.
#
# html_sidebars = {}


# -- Options for HTMLHelp output ---------------------------------------------

# Output file base name for HTML help builder.
htmlhelp_basename = 'mwemdoc'


# -- Options for LaTeX output ------------------------------------------------

latex_elements = {
    # The paper size ('letterpaper' or 'a4paper').
    #
    # 'papersize': 'letterpaper',

    # The font size ('10pt', '11pt' or '12pt').
    #
    # 'pointsize': '10pt',

    # Additional stuff for the LaTeX preamble.
    #
    # 'preamble': '',

    # Latex figure (float) alignment
    #
    # 'figure_align': 'htbp',
}

# Grouping the document tree into LaTeX files. List of tuples
# (source start file, target name, title,
#  author, documentclass [howto, manual, or own class]).
latex_documents = [
    (master_doc, 'mwem.tex', 'MWEM Documentation',
     'mwem', 'manual'),
]


# -- Options for manual page output ------------------------------------------

# One entry per manual page. List of tuples
# (source start file, name, description, authors, manual section).
man_pages = [
    (master_doc, 'mwem', 'MWEM Documentation',
     [author], 1)
]


# -- Options for Texinfo output ----------------------------------------------

# Grouping the document tree into Texinfo files. List of tuples
# (source start file, target name, title, author,
#  dir menu entry, description, category)
texinfo_documents = [
    (master_doc, 'mwem', 'MWEM Documentation',
     author, 'mwem', 'Markovian Weighted Ensemble Milestoning (MWEM)',
     'Miscellaneous'),
]


# -- Extension configuration -------------------------------------------------
```
