## Supplementary material for "Markovian Weighted Ensemble Milestoning (M-WEM): Long-time Kinetics from Short Trajectories": MWEM-main.zip: make.bat

```
@ECHO OFF

pushd %~dp0

REM Command file for Sphinx documentation

if "%SPHINXBUILD%" == "" (
	set SPHINXBUILD=sphinx-build
)
set SOURCEDIR=.
set BUILDDIR=_build
set SPHINXPROJ=mwem

if "%1" == "" goto help

%SPHINXBUILD% >NUL 2>NUL
if errorlevel 9009 (
	echo.
	echo.The 'sphinx-build' command was not found. Make sure you have Sphinx
	echo.installed, then set the SPHINXBUILD environment variable to point
	echo.to the full path of the 'sphinx-build' executable. Alternatively you
	echo.may add the Sphinx directory to PATH.
	echo.
	echo.If you don't have Sphinx installed, grab it from
	echo.http://sphinx-doc.org/
	exit /b 1
)

%SPHINXBUILD% -M %1 %SOURCEDIR% %BUILDDIR% %SPHINXOPTS%
goto end

:help
%SPHINXBUILD% -M help %SOURCEDIR% %BUILDDIR% %SPHINXOPTS%

:end
popd
```
