## Supplementary material for "Markovian Weighted Ensemble Milestoning (M-WEM): Long-time Kinetics from Short Trajectories": MWEM-main.zip: Makefile

```
# Minimal makefile for Sphinx documentation
#

# You can set these variables from the command line.
SPHINXOPTS    =
SPHINXBUILD   = sphinx-build
SPHINXPROJ    = mwem
SOURCEDIR     = .
BUILDDIR      = _build

# Put it first so that "make" without argument is like "make help".
help:
	@$(SPHINXBUILD) -M help "$(SOURCEDIR)" "$(BUILDDIR)" $(SPHINXOPTS) $(O)

.PHONY: help Makefile

# Catch-all target: route all unknown targets to Sphinx using the new
# "make mode" option.  $(O) is meant as a shortcut for $(SPHINXOPTS).
%: Makefile
	@$(SPHINXBUILD) -M $@ "$(SOURCEDIR)" "$(BUILDDIR)" $(SPHINXOPTS) $(O)
```
