## Supplementary material for "Markovian Weighted Ensemble Milestoning (M-WEM): Long-time Kinetics from Short Trajectories": MWEM-main.zip: README.md

# Compiling MWEM's Documentation

The docs for this project are built with [Sphinx](http://www.sphinx-doc.org/en/master/).
To compile the docs, first ensure that Sphinx and the ReadTheDocs theme are installed.


```bash
conda install sphinx sphinx_rtd_theme 
```


Once installed, you can use the `Makefile` in this directory to compile static HTML pages by
```bash
make html
```

The compiled docs will be in the `_build` directory and can be viewed by opening `index.html` (which may itself 
be inside a directory called `html/` depending on what version of Sphinx is installed).


A configuration file for [Read The Docs](https://readthedocs.org/) (readthedocs.yaml) is included in the top level of the repository. To use Read the Docs to host your documentation, go to https://readthedocs.org/ and connect this repository. You may need to change your default branch to `main` under Advanced Settings for the project.

If you would like to use Read The Docs with `autodoc` (included automatically) and your package has dependencies, you will need to include those dependencies in your documentation yaml file (`docs/requirements.yaml`).
