## Supplementary material for "Markovian Weighted Ensemble Milestoning (M-WEM): Long-time Kinetics from Short Trajectories": MWEM-main.zip: __init__.py

```
"""
MWEM
Markovian Weighted Ensemble Milestoning (MWEM)
"""

# Add imports here
from .functions import *
from .milestone_analysis_functions import *
from .westpa_analysis_functions import *
from .milestone_analysis import *
from .committor import *

# Handle versioneer
from ._version import get_versions
versions = get_versions()
__version__ = versions['version']
__git_revision__ = versions['full-revisionid']
del get_versions, versions
```
