## Supplementary material for "Markovian Weighted Ensemble Milestoning (M-WEM): Long-time Kinetics from Short Trajectories": MWEM-main.zip: _version.py

```
# This file helps to compute a version number in source trees obtained from
# git-archive tarball (such as those provided by githubs download-from-tag
# feature). Distribution tarballs (built by setup.py sdist) and build
# directories (produced by setup.py build) will contain a much shorter file
# that just contains the computed version number.

# This file is released into the public domain. Generated by
# versioneer-0.18 (https://github.com/warner/python-versioneer)

"""Git implementation of _version.py."""

import errno
import os
import re
import subprocess
import sys


def get_keywords():
    """Get the keywords needed to look up the version information."""
    # these strings will be replaced by git during git-archive.
    # setup.py/versioneer.py will grep for the variable names, so they must
    # each be defined on a line of their own. _version.py will just call
    # get_keywords().
    git_refnames = "$Format:%d$"
    git_full = "$Format:%H$"
    git_date = "$Format:%ci$"
    keywords = {"refnames": git_refnames, "full": git_full, "date": git_date}
    return keywords


class VersioneerConfig:
    """Container for Versioneer configuration parameters."""


def get_config():
    """Create, populate and return the VersioneerConfig() object."""
    # these strings are filled in when 'setup.py versioneer' creates
    # _version.py
    cfg = VersioneerConfig()
    cfg.VCS = "git"
    cfg.style = "pep440"
    cfg.tag_prefix = ""
    cfg.parentdir_prefix = "None"
    cfg.versionfile_source = "mwem/_version.py"
    cfg.verbose = False
    return cfg


class NotThisMethod(Exception):
    """Exception raised if a method is not valid for the current scenario."""


LONG_VERSION_PY = {}
HANDLERS = {}


def register_vcs_handler(vcs, method):  # decorator
    """Decorator to mark a method as the handler for a particular VCS."""
    def decorate(f):
        """Store f in HANDLERS[vcs][method]."""
        if vcs not in HANDLERS:
            HANDLERS[vcs] = {}
        HANDLERS[vcs][method] = f
        return f
    return decorate


def run_command(commands, args, cwd=None, verbose=False, hide_stderr=False,
                env=None):
    """Call the given command(s)."""
    assert isinstance(commands, list)
    p = None
    for c in commands:
        try:
            dispcmd = str([c] + args)
            # remember shell=False, so use git.cmd on windows, not just git
            p = subprocess.Popen([c] + args, cwd=cwd, env=env,
                                 stdout=subprocess.PIPE,
                                 stderr=(subprocess.PIPE if hide_stderr
                                         else None))
            break
        except EnvironmentError:
            e = sys.exc_info()[1]
            if e.errno == errno.ENOENT:
                continue
            if verbose:
                print("unable to run %s" % dispcmd)
                print(e)
            return None, None
    else:
        if verbose:
            print("unable to find command, tried %s" % (commands,))
        return None, None
    stdout = p.communicate()[0].strip()
    if sys.version_info[0] >= 3:
        stdout = stdout.decode()
    if p.returncode != 0:
        if verbose:
            print("unable to run %s (error)" % dispcmd)
            print("stdout was %s" % stdout)
        return None, p.returncode
    return stdout, p.returncode


def versions_from_parentdir(parentdir_prefix, root, verbose):
    """Try to determine the version from the parent directory name.

    Source tarballs conventionally unpack into a directory that includes both
    the project name and a version string. We will also support searching up
    two directory levels for an appropriately named parent directory
    """
    rootdirs = []

    for i in range(3):
        dirname = os.path.basename(root)
        if dirname.startswith(parentdir_prefix):
            return {"version": dirname[len(parentdir_prefix):],
                    "full-revisionid": None,
                    "dirty": False, "error": None, "date": None}
        else:
            rootdirs.append(root)
            root = os.path.dirname(root)  # up a level

    if verbose:
        print("Tried directories %s but none started with prefix %s" %
              (str(rootdirs), parentdir_prefix))
    raise NotThisMethod("rootdir doesn't start with parentdir_prefix")


@register_vcs_handler("git", "get_keywords")
def git_get_keywords(versionfile_abs):
    """Extract version information from the given file."""
    # the code embedded in _version.py can just fetch the value of these
    # keywords. When used from setup.py, we don't want to import _version.py,
    # so we do it with a regexp instead. This function is not used from
    # _version.py.
    keywords = {}
    try:
        f = open(versionfile_abs, "r")
        for line in f.readlines():
            if line.strip().startswith("git_refnames ="):
                mo = re.search(r'=\s*"(.*)"', line)
                if mo:
                    keywords["refnames"] = mo.group(1)
            if line.strip().startswith("git_full ="):
                mo = re.search(r'=\s*"(.*)"', line)
                if mo:
                    keywords["full"] = mo.group(1)
            if line.strip().startswith("git_date ="):
                mo = re.search(r'=\s*"(.*)"', line)
                if mo:
                    keywords["date"] = mo.group(1)
        f.close()
    except EnvironmentError:
        pass
    return keywords


@register_vcs_handler("git", "keywords")
def git_versions_from_keywords(keywords, tag_prefix, verbose):
    """Get version information from git keywords."""
    if not keywords:
        raise NotThisMethod("no keywords at all, weird")
    date = keywords.get("date")
    if date is not None:
        # git-2.2.0 added "%cI", which expands to an ISO-8601 -compliant
        # datestamp. However we prefer "%ci" (which expands to an "ISO-8601
        # -like" string, which we must then edit to make compliant), because
        # it's been around since git-1.5.3, and it's too difficult to
        # discover which version we're using, or to work around using an
        # older one.
        date = date.strip().replace(" ", "T", 1).replace(" ", "", 1)
    refnames = keywords["refnames"].strip()
    if refnames.startswith("$Format"):
        if verbose:
            print("keywords are unexpanded, not using")
        raise NotThisMethod("unexpanded keywords, not a git-archive tarball")
    refs = set([r.strip() for r in refnames.strip("()").split(",")])
    # starting in git-1.8.3, tags are listed as "tag: foo-1.0" instead of
    # just "foo-1.0". If we see a "tag: " prefix, prefer those.
    TAG = "tag: "
    tags = set([r[len(TAG):] for r in refs if r.startswith(TAG)])
    if not tags:
        # Either we're using git < 1.8.3, or there really are no tags. We use
        # a heuristic: assume all version tags have a digit. The old git %d
        # expansion behaves like git log --decorate=short and strips out the
        # refs/heads/ and refs/tags/ prefixes that would let us distinguish
        # between branches and tags. By ignoring refnames without digits, we
        # filter out many common branch names like "release" and
        # "stabilization", as well as "HEAD" and "master".
        tags = set([r for r in refs if re.search(r'\d', r)])
        if verbose:
            print("discarding '%s', no digits" % ",".join(refs - tags))
    if verbose:
        print("likely tags: %s" % ",".join(sorted(tags)))
    for ref in sorted(tags):
        # sorting will prefer e.g. "2.0" over "2.0rc1"
        if ref.startswith(tag_prefix):
            r = ref[len(tag_prefix):]
            if verbose:
                print("picking %s" % r)
            return {"version": r,
                    "full-revisionid": keywords["full"].strip(),
                    "dirty": False, "error": None,
                    "date": date}
    # no suitable tags, so version is "0+unknown", but full hex is still there
    if verbose:
        print("no suitable tags, using unknown + full revision id")
    return {"version": "0+unknown",
            "full-revisionid": keywords["full"].strip(),
            "dirty": False, "error": "no suitable tags", "date": None}


@register_vcs_handler("git", "pieces_from_vcs")
def git_pieces_from_vcs(tag_prefix, root, verbose, run_command=run_command):
    """Get version from 'git describe' in the root of the source tree.

    This only gets called if the git-archive 'subst' keywords were *not*
    expanded, and _version.py hasn't already been rewritten with a short
    version string, meaning we're inside a checked out source tree.
    """
    GITS = ["git"]
    if sys.platform == "win32":
        GITS = ["git.cmd", "git.exe"]

    out, rc = run_command(GITS, ["rev-parse", "--git-dir"], cwd=root,
                          hide_stderr=True)
    if rc != 0:
        if verbose:
            print("Directory %s not under git control" % root)
        raise NotThisMethod("'git rev-parse --git-dir' returned error")

    # if there is a tag matching tag_prefix, this yields TAG-NUM-gHEX[-dirty]
    # if there isn't one, this yields HEX[-dirty] (no NUM)
    describe_out, rc = run_command(GITS, ["describe", "--tags", "--dirty",
                                          "--always", "--long",
                                          "--match", "%s*" % tag_prefix],
                                   cwd=root)
    # --long was added in git-1.5.5
    if describe_out is None:
        raise NotThisMethod("'git describe' failed")
    describe_out = describe_out.strip()
    full_out, rc = run_command(GITS, ["rev-parse", "HEAD"], cwd=root)
    if full_out is None:
        raise NotThisMethod("'git rev-parse' failed")
    full_out = full_out.strip()

    pieces = {}
    pieces["long"] = full_out
    pieces["short"] = full_out[:7]  # maybe improved later
    pieces["error"] = None

    # parse describe_out. It will be like TAG-NUM-gHEX[-dirty] or HEX[-dirty]
    # TAG might have hyphens.
    git_describe = describe_out

    # look for -dirty suffix
    dirty = git_describe.endswith("-dirty")
    pieces["dirty"] = dirty
    if dirty:
        git_describe = git_describe[:git_describe.rindex("-dirty")]

    # now we have TAG-NUM-gHEX or HEX

    if "-" in git_describe:
        # TAG-NUM-gHEX
        mo = re.search(r'^(.+)-(\d+)-g([0-9a-f]+)$', git_describe)
        if not mo:
            # unparseable. Maybe git-describe is misbehaving?
            pieces["error"] = ("unable to parse git-describe output: '%s'"
                               % describe_out)
            return pieces

        # tag
        full_tag = mo.group(1)
        if not full_tag.startswith(tag_prefix):
            if verbose:
                fmt = "tag '%s' doesn't start with prefix '%s'"
                print(fmt % (full_tag, tag_prefix))
            pieces["error"] = ("tag '%s' doesn't start with prefix '%s'"
                               % (full_tag, tag_prefix))
            return pieces
        pieces["closest-tag"] = full_tag[len(tag_prefix):]

        # distance: number of commits since tag
        pieces["distance"] = int(mo.group(2))

        # commit: short hex revision ID
        pieces["short"] = mo.group(3)

    else:
        # HEX: no tags
        pieces["closest-tag"] = None
        count_out, rc = run_command(GITS, ["rev-list", "HEAD", "--count"],
                                    cwd=root)
        pieces["distance"] = int(count_out)  # total number of commits

    # commit date: see ISO-8601 comment in git_versions_from_keywords()
    date = run_command(GITS, ["show", "-s", "--format=%ci", "HEAD"],
                       cwd=root)[0].strip()
    pieces["date"] = date.strip().replace(" ", "T", 1).replace(" ", "", 1)

    return pieces


def plus_or_dot(pieces):
    """Return a + if we don't already have one, else return a ."""
    if "+" in pieces.get("closest-tag", ""):
        return "."
    return "+"


def render_pep440(pieces):
    """Build up version string, with post-release "local version identifier".

    Our goal: TAG[+DISTANCE.gHEX[.dirty]] . Note that if you
    get a tagged build and then dirty it, you'll get TAG+0.gHEX.dirty

    Exceptions:
    1: no tags. git_describe was just HEX. 0+untagged.DISTANCE.gHEX[.dirty]
    """
    if pieces["closest-tag"]:
        rendered = pieces["closest-tag"]
        if pieces["distance"] or pieces["dirty"]:
            rendered += plus_or_dot(pieces)
            rendered += "%d.g%s" % (pieces["distance"], pieces["short"])
            if pieces["dirty"]:
                rendered += ".dirty"
    else:
        # exception #1
        rendered = "0+untagged.%d.g%s" % (pieces["distance"],
                                          pieces["short"])
        if pieces["dirty"]:
            rendered += ".dirty"
    return rendered


def render_pep440_pre(pieces):
    """TAG[.post.devDISTANCE] -- No -dirty.

    Exceptions:
    1: no tags. 0.post.devDISTANCE
    """
    if pieces["closest-tag"]:
        rendered = pieces["closest-tag"]
        if pieces["distance"]:
            rendered += ".post.dev%d" % pieces["distance"]
    else:
        # exception #1
        rendered = "0.post.dev%d" % pieces["distance"]
    return rendered


def render_pep440_post(pieces):
    """TAG[.postDISTANCE[.dev0]+gHEX] .

    The ".dev0" means dirty. Note that .dev0 sorts backwards
    (a dirty tree will appear "older" than the corresponding clean one),
    but you shouldn't be releasing software with -dirty anyways.

    Exceptions:
    1: no tags. 0.postDISTANCE[.dev0]
    """
    if pieces["closest-tag"]:
        rendered = pieces["closest-tag"]
        if pieces["distance"] or pieces["dirty"]:
            rendered += ".post%d" % pieces["distance"]
            if pieces["dirty"]:
                rendered += ".dev0"
            rendered += plus_or_dot(pieces)
            rendered += "g%s" % pieces["short"]
    else:
        # exception #1
        rendered = "0.post%d" % pieces["distance"]
        if pieces["dirty"]:
            rendered += ".dev0"
        rendered += "+g%s" % pieces["short"]
    return rendered


def render_pep440_old(pieces):
    """TAG[.postDISTANCE[.dev0]] .

    The ".dev0" means dirty.

    Eexceptions:
    1: no tags. 0.postDISTANCE[.dev0]
    """
    if pieces["closest-tag"]:
        rendered = pieces["closest-tag"]
        if pieces["distance"] or pieces["dirty"]:
            rendered += ".post%d" % pieces["distance"]
            if pieces["dirty"]:
                rendered += ".dev0"
    else:
        # exception #1
        rendered = "0.post%d" % pieces["distance"]
        if pieces["dirty"]:
            rendered += ".dev0"
    return rendered


def render_git_describe(pieces):
    """TAG[-DISTANCE-gHEX][-dirty].

    Like 'git describe --tags --dirty --always'.

    Exceptions:
    1: no tags. HEX[-dirty]  (note: no 'g' prefix)
    """
    if pieces["closest-tag"]:
        rendered = pieces["closest-tag"]
        if pieces["distance"]:
            rendered += "-%d-g%s" % (pieces["distance"], pieces["short"])
    else:
        # exception #1
        rendered = pieces["short"]
    if pieces["dirty"]:
        rendered += "-dirty"
    return rendered


def render_git_describe_long(pieces):
    """TAG-DISTANCE-gHEX[-dirty].

    Like 'git describe --tags --dirty --always -long'.
    The distance/hash is unconditional.

    Exceptions:
    1: no tags. HEX[-dirty]  (note: no 'g' prefix)
    """
    if pieces["closest-tag"]:
        rendered = pieces["closest-tag"]
        rendered += "-%d-g%s" % (pieces["distance"], pieces["short"])
    else:
        # exception #1
        rendered = pieces["short"]
    if pieces["dirty"]:
        rendered += "-dirty"
    return rendered


def render(pieces, style):
    """Render the given version pieces into the requested style."""
    if pieces["error"]:
        return {"version": "unknown",
                "full-revisionid": pieces.get("long"),
                "dirty": None,
                "error": pieces["error"],
                "date": None}

    if not style or style == "default":
        style = "pep440"  # the default

    if style == "pep440":
        rendered = render_pep440(pieces)
    elif style == "pep440-pre":
        rendered = render_pep440_pre(pieces)
    elif style == "pep440-post":
        rendered = render_pep440_post(pieces)
    elif style == "pep440-old":
        rendered = render_pep440_old(pieces)
    elif style == "git-describe":
        rendered = render_git_describe(pieces)
    elif style == "git-describe-long":
        rendered = render_git_describe_long(pieces)
    else:
        raise ValueError("unknown style '%s'" % style)

    return {"version": rendered, "full-revisionid": pieces["long"],
            "dirty": pieces["dirty"], "error": None,
            "date": pieces.get("date")}


def get_versions():
    """Get version information or return default if unable to do so."""
    # I am in _version.py, which lives at ROOT/VERSIONFILE_SOURCE. If we have
    # __file__, we can work backwards from there to the root. Some
    # py2exe/bbfreeze/non-CPython implementations don't do __file__, in which
    # case we can only use expanded keywords.

    cfg = get_config()
    verbose = cfg.verbose

    try:
        return git_versions_from_keywords(get_keywords(), cfg.tag_prefix,
                                          verbose)
    except NotThisMethod:
        pass

    try:
        root = os.path.realpath(__file__)
        # versionfile_source is the relative path from the top of the source
        # tree (where the .git directory might live) to this file. Invert
        # this to find the root from __file__.
        for i in cfg.versionfile_source.split('/'):
            root = os.path.dirname(root)
    except NameError:
        return {"version": "0+unknown", "full-revisionid": None,
                "dirty": None,
                "error": "unable to find root of source tree",
                "date": None}

    try:
        pieces = git_pieces_from_vcs(cfg.tag_prefix, root, verbose)
        return render(pieces, cfg.style)
    except NotThisMethod:
        pass

    try:
        if cfg.parentdir_prefix:
            return versions_from_parentdir(cfg.parentdir_prefix, root, verbose)
    except NotThisMethod:
        pass

    return {"version": "0+unknown", "full-revisionid": None,
            "dirty": None,
            "error": "unable to compute version", "date": None}
```
