## Supplementary material for "Markovian Weighted Ensemble Milestoning (M-WEM): Long-time Kinetics from Short Trajectories": MWEM-main.zip: committor.py

```
import numpy as np


def K_matrix(N_i_j):
    K = np.zeros(N_i_j.shape)
    for i in range(len(N_i_j)):
        tot = np.sum(N_i_j[i])
        #print(tot)
        K[i] = N_i_j[i] / tot

    return K

def committor(K_matrix,tol=1E-3,Niter=10000):
    N = len(K_matrix)
    K = K_matrix.copy()
    #make all elements of the first row to zero (bound state)
    for i in range(N):
        K[0,i] = 0.0
        K[N-1,i] = 0.0 #this is needed for the boundary condition in next step
    #boundary condition: trajectories get stuck at last milesone
    K[N-1,N-1] = 1.0

    K_new = K
    C = K[:,N-1]
    for i in range(Niter):
        K_new = np.dot(K_new,K)
        C_new = K_new[:,N-1] #commitor is the last column of K_new
        #print(np.linalg.norm(C_new - C))
        if np.linalg.norm(C_new - C) < tol:
            C = C_new
            print('converged at iteration: ',i)
            break
        else :
            C = C_new
    return C
```
