## Supplementary material for "Markovian Weighted Ensemble Milestoning (M-WEM): Long-time Kinetics from Short Trajectories": MWEM-main.zip: README.md

# Sample Package Data

This directory contains sample additional data you may want to include with your package.
This is a place where non-code related additional information (such as data files, molecular structures,  etc.) can 
go that you want to ship alongside your code.

Please note that it is not recommended to place large files in your git directory. If your project requires files larger
than a few megabytes in size it is recommended to host these files elsewhere. This is especially true for binary files
as the `git` structure is unable to correctly take updates to these files and will store a complete copy of every version
in your `git` history which can quickly add up. As a note most `git` hosting services like GitHub have a 1 GB per repository
cap.

## Including package data

Modify your package's `setup.py` file and the `setup()` command. Include the 
[`package_data`](http://setuptools.readthedocs.io/en/latest/setuptools.html#basic-use) keyword and point it at the 
correct files.

## Manifest

* `look_and_say.dat`: first entries of the "Look and Say" integer series, sequence [A005150](https://oeis.org/A005150)
