## Supplementary material for "Markovian Weighted Ensemble Milestoning (M-WEM): Long-time Kinetics from Short Trajectories": MWEM-main.zip: functions.py

```
"""
functions.py
Markovian Weighted Ensemble Milestoning (MWEM)

Handles the primary functions
"""


def canvas(with_attribution=True):
    """
    Placeholder function to show example docstring (NumPy format)

    Replace this function and doc string for your own project

    Parameters
    ----------
    with_attribution : bool, Optional, default: True
        Set whether or not to display who the quote is from

    Returns
    -------
    quote : str
        Compiled string including quote and optional attribution
    """

    quote = "The code is but a canvas to our imagination."
    if with_attribution:
        quote += "\n\t- Adapted from Henry David Thoreau"
    return quote


if __name__ == "__main__":
    # Do something if this file is invoked on its own
    print(canvas())
```
