## Supplementary material for "Markovian Weighted Ensemble Milestoning (M-WEM): Long-time Kinetics from Short Trajectories": MWEM-main.zip: milestone_analysis.py

```
import numpy as np
import pickle
import scipy.stats as st
import mwem


def milestoning(crossings_file,weights_file,indices,milestones,cutoff,dt,start_milestone,end_milestone,
        radial=False,cell_prob_file='cell_prob.dat',reverse=False,returnNR=False,doMCMC=False,numMCMC=50000,intervalMCMC=10):
    '''
    Inputs
    ---------------------------------------------------------------------
    crossings_file: File name of the .pkl file containing crosssings data
                    If the file path is 'cell_3.5/crossings_40.pkl', enter
                    crossings_file = crossings_40.pkl

    weights_file:   File name of the .txt file containing trajectory weights
                    If the file path is 'cell_3.5/weights_40.txt', enter
                    weights_file = weights_40.txt

    indices:        Python list containing cell indices.
                    If cell indices are cell_0, cell_1, cell_2.5, cell_3.5
                    indices = [0,1,2.5,3.5]

    milestones:     List of milestone posiions. In 1D case
                    len(milestones) = number of cells + 1 

    cutoff:         Minumum value of weight to be considered

    dt:             Frequency at which the trajectory is saved
                    e.g. If the trajectory saved every 20 fs and the
                    desired unit of MFPT is millisecond then 
                    dt = 20.0*1E-12

    start_milestone: Zero based index of the milestone depicting the starting state

    end_milestone: Zero based index of the milestone depicting the final state

    radial:         If True, then milestones are sperical
                    Perform Jacobian correction

    cell_prob_file: Filename where the output of equilibrium probability 
                    per cell will be printed (default: 'cell_prob.dat')

    reverse:        If True calculates the MFPT of the backward (reverse) direction
                    default: False

    returnNR:       IF True, returns the N_ij and R_i matrices

    doMCMC:         If True, Monte Carlo error analysis is performed

    numMCMC:        Number of MCMC steps are to be performed for error estimation
                    Ignored if doMCMC=False.

    intervalMCMC:   Frequency at which MCMC matrices are sampled
                    Ignored if doMCMC=False.

    ------------------------------------------------------------------------------

    Outputs (returns)
    ------------------------------------------------------------------------------

    mfpt:           Mean first passage time from start milestone to end milestone

    mean_mfpt:      Avg MFPT computed from MCMC bootstrapping error analysis
                    Not returned if doMCMC=False

    lower_conf:     Lower confidence interval
                    Not returned if doMCMC=False

    upper_conf:     Upper confidence interval
                    Not returned if doMCMC=False

    -------------------------------------------------------------------------------

    Prints
    ------------------------------------------------------------------------------
    cell_prob_file: File containing probability distribution per cell

    '''

    N = len(milestones)

    #Check dimensions of lists in 1D case
    assert N == len(indices)+1

    

    #construct matrices
    k_a_b = np.zeros((N-1,N-1))

    #loop over cell
    for i in range(N-1):

        #filenames
        crossings_file_name = 'cell_%s/'%(str(indices[i]))+crossings_file
        weights_file_name = 'cell_%s/'%(str(indices[i]))+weights_file

        #load crossings trajectory
        crossings = pickle.load(open(crossings_file_name,'rb'))
        #print(i,len(crossings))
        #load weights of each trajectory
        weights = np.loadtxt(weights_file_name)


        #loop over trajectory traces
        for j in range(len(crossings)):
            #store trajectory trace in array
            trajectory_crossings = np.array(crossings[j])
            #print(trajectory_crossings.shape)

            if len(trajectory_crossings) != 0:

                #Compute Nab and Ta for individual traces
                N_a_b_traj, T_a_traj = mwem.compute_Nab_Ta(trajectory_crossings,N,i)
        
                #Compute kab matrix for individual traces
                k_a_b_traj = mwem.compute_k_a_b(N_a_b_traj, T_a_traj, i, milestones)

                #perform weighted sum over k_a_b's to get the overall k_a_b for the cell
                if weights[j] > cutoff:
                    k_a_b += weights[j]*k_a_b_traj


    #compute the stationary probability distrubution
    p = mwem.probability(k_a_b,Niter=100000)

    if radial == True:

        for i in range(len(p)):
            p[i] *= ((milestones[i]+milestones[i+1])*0.5)**2
        p /= np.sum(p)

    #print(k_a_b)
    #print(p)

    f1 = open(cell_prob_file,'w')
    print('#Normalized equilibrium probability at each cell',file=f1)
    for i in range(N-1):
        print((milestones[i] + milestones[i+1])*0.5,p[i],file=f1)
    f1.close()

    #--------------------------------------------------------
    #Compute kinetics
    #--------------------------------------------------------

    N_i_j = np.zeros((N,N))
    R_i = np.zeros(N)
    Nhit = np.zeros((N,N))

    #loop over cell
    for i in range(N-1):
        
        #filenames
        crossings_file_name = 'cell_%s/'%(str(indices[i]))+crossings_file
        weights_file_name = 'cell_%s/'%(str(indices[i]))+weights_file

        #load crossings trajectory
        crossings = pickle.load(open(crossings_file_name,'rb'))
        #print(i,len(crossings))
        #load weights of each trajectory
        weights = np.loadtxt(weights_file_name)

        #loop over trajectory traces
        for j in range(len(crossings)):
            #store trajectory trace in array
            trajectory_crossings = np.array(crossings[j])
        
            if len(trajectory_crossings) != 0:

                #compute Nij and Ri for individual trajectories
                N_ij_traj, R_i_traj = mwem.compute_Nij_Ri(trajectory_crossings,p,i,N)

                #compute Nhit for individual trajectories
                Nhit_traj = mwem.compute_Nhit(trajectory_crossings,i,N)

                #perform weighted sum
                if weights[j] > cutoff:
                    N_i_j += weights[j]*N_ij_traj
                    R_i   += weights[j]*R_i_traj

                #sum the Nhits
                Nhit += Nhit_traj

    #compute Q matrix
    if reverse == True:
        Q = mwem.Q_matrix_rev(N_i_j,R_i,N)
    else :
        Q = mwem.Q_matrix(N_i_j,R_i,N)

    #compute MFPTs
    start = start_milestone
    end = end_milestone

    #print(N_i_j)

    mfpt = mwem.MFPT(Q,0,end)[start]*dt

    #compute probabiliy K_i,i-1 for k_on calculation
    K_i_i_minus_1 = N_i_j[end_milestone-1,end_milestone-2]/np.sum(N_i_j[end_milestone-1])
    #K_rev_list = []

    #estimate error using MCMC if doMCMC is True
    if doMCMC == True:
        N_total = numMCMC
        interval = intervalMCMC
        Q_array = mwem.Monte_Carlo_bootstrapping(N_total,Q,R_i,Nhit,interval)

        MFPT_list = []
        for Q_sampled in Q_array:

            MFPTs = mwem.MFPT(Q_sampled,0,end) #MFPT computation has to start at 0 irrespective of initial milestone

            MFPT_list.append(dt*MFPTs[start])

        MFPT_list = np.array(MFPT_list)
        conf = st.t.interval(alpha=0.95, df=len(MFPT_list)-1, loc=np.mean(MFPT_list), scale=st.sem(MFPT_list))
    
        mean_mfpt = np.mean(MFPT_list)

        lower_conf = conf[0]

        upper_conf = conf[1]
    
    if doMCMC == True:
        if returnNR == True:
            return mfpt, mean_mfpt, lower_conf, upper_conf, K_i_i_minus_1, N_i_j, R_i
        else:
            return mfpt, mean_mfpt, lower_conf, upper_conf, K_i_i_minus_1
    else :
        if returnNR == True:
            return mfpt, K_i_i_minus_1, N_i_j, R_i
        else:
            return mfpt, K_i_i_minus_1

    
```
