## Supplementary material for "Markovian Weighted Ensemble Milestoning (M-WEM): Long-time Kinetics from Short Trajectories": MWEM-main.zip: milestone_analysis_functions.py

```
import numpy as np


#---------------------------------------------------------
#Read in transition counts between cells (N_alpha_beta)
#---------------------------------------------------------

def compute_Nab_Ta(trajectory_crossings,N,i):
    '''
    Compute N_a_b and T_a for one individual trajectory at one individual cell
    --------------------------------------------------------------------------
    trajectory_crossings : time vs crossings time series
    N : number of milestones = number of cells + 1
    i : cell index
    '''
    
    N_a_b = np.zeros((N-1,N-1))

    T_a = np.zeros(N-1)

    l = trajectory_crossings
    right_crossings = np.sum(l[:,1])
    left_crossings = len(l) - right_crossings

    if i==0 :
        N_a_b[i,i+1] = right_crossings

    elif i==N-2 :
        N_a_b[i,i-1] = left_crossings

    else :
        N_a_b[i,i-1] = left_crossings
        N_a_b[i,i+1] = right_crossings

    T_a[i] = l[-1,0]

    return N_a_b, T_a

#-----------------------------------------------------------
#Compute probabilities and free energy per cell
#-----------------------------------------------------------

#compute transition flux matrix (k_alpha_beta)
def compute_k_a_b(N_a_b, T_a, i, milestones):
    '''
    Compute transition flux matrix (k_alpha_beta) for one individual cell
    and one individual trajectory trace
    '''

    N = len(N_a_b) + 1

    k_a_b = np.zeros((N-1,N-1))

    k_a_b[i] += (N_a_b[i]/T_a[i])

    return k_a_b

#----- Perform self consistent iterations to calculate stationar state ----#

def probability(k_a_b,Niter=10000):
    N = len(k_a_b) + 1

    #initial guess
    p_init = np.ones(N-1)
    p_init /= len(p_init)


    p = p_init


    for j in range(Niter):
        p_new = np.zeros(N-1)
        for i in range(N-1):
            if i==0:
                p_new[i] = p[i+1]*k_a_b[i+1,i]/k_a_b[i,i+1]

            elif i==N-2:
                p_new[i] = p[i-1]*k_a_b[i-1,i]/k_a_b[i,i-1]

            else :
                p_new[i] = (p[i-1]*k_a_b[i-1,i] + p[i+1]*k_a_b[i+1,i])/(k_a_b[i,i-1] + k_a_b[i,i+1])

        p_new /= np.sum(p_new)
        p = p_new

    return p


#-----------------------------------------------------------------------------
#Construct the Q matrix for kinetics calculation
#-----------------------------------------------------------------------------

def compute_Nij_Ri(trajectory_crossings,p,i,N):
    '''Construct Nij and Ri for an individual trajectory in one individual cell
    '''
    N_i_j = np.zeros((N,N))
    R_i = np.zeros(N)

#for i in range(N-1):
    l = trajectory_crossings

    T_a = l[-1,0]
    
    #compute N_i_j
    for j in range(1,len(l)):
        #hitting a milestone coming from a different milestone
        if l[j-1,1] == 0 and l[j,1] == 1 :
            N_i_j[i,i+1] += 1.0 * p[i]/T_a
        elif l[j-1,1] == 1 and l[j,1] == 0 :
            N_i_j[i+1,i] += 1.0 * p[i]/T_a

    #compute R_i
    for j in range(1,len(l)):
        if l[j-1,1] == 0:
            R_i[i] += (l[j,0] - l[j-1,0]) * p[i]/T_a
        elif l[j-1,1] == 1:
            R_i[i+1] += (l[j,0] - l[j-1,0]) * p[i]/T_a
    
    return N_i_j, R_i

#Construct the matrix of number of hitting points for one trajectory trace
def compute_Nhit(trajectory_crossings,i,N):
    Nhit = np.zeros((N,N))

    l = trajectory_crossings

    for j in range(1,len(l)):
        #hitting a milestone coming from a different milestone
        if l[j-1,1] == 0 and l[j,1] == 1 :
            Nhit[i,i+1] += 1.0 
        elif l[j-1,1] == 1 and l[j,1] == 0 :
            Nhit[i+1,i] += 1.0 

    return Nhit

def Q_matrix(N_i_j,R_i,N):
    '''
    Construct the Q matrix for kinetics calculation
    '''
    
    Q = np.zeros((N,N))

    for i in range(N):
        for j in range(N):
            if R_i[i] != 0:
                Q[i,j] = N_i_j[i,j]/R_i[i]

    for i in range(N):
        Q[i,i] = -np.sum(Q[i])

    return Q    

def Q_matrix_rev(N_i_j,R_i,N):
    '''
    Construct the reverse Q matrix for kinetics calculation
    of the backward transition

    Replacing i with N-1-i so that indices change linke this
    0 -> N-1
    1 -> N-2
    ...
    N-1 -> 0
    '''

    Q = np.zeros((N,N))

    for i in range(N):
        for j in range(N):
            if R_i[N-1-i] != 0:
                Q[i,j] = N_i_j[N-1-i,N-1-j]/R_i[N-1-i]

    for i in range(N):
        Q[i,i] = -np.sum(Q[i])

    return Q


#-------------------------------------------------------------------------------
#Construct the truncated rate matrix and compute MFPT
#-------------------------------------------------------------------------------


def MFPT(Q,start,end):
    '''
    Construct the truncated rate matrix and compute MFPT
    '''

    #dimention of rate matrix
    M = end - start

    Q_rate = np.zeros((M,M))

    I1 = np.ones(M)
    #print(I1)

    for i in range(M):
        for j in range(M):
            Q_rate[i,j] = Q[start+i,start+j]

    MFPTs = -np.linalg.solve(Q_rate,I1)

    return MFPTs

def Monte_Carlo_bootstrapping(N_total,K,t,Nhit,interval):
    '''Perform nonreversible element shift Monte Carlo to sample rate matrices
    Input
    -----
    N_total : Total number of MC moves (accepted and rejected)
    K : Rate Matrix
    t : lifetime vector (R_i)
    Nhit : Matrix containing number of hitting points
    interval : After how many MC moves a matrix is sampled
    Returns
    -------
    Q_list : (n,N,N) dim array where n is the number of sampled
             rate matrices
    '''
    N = len(t)

    Q_list = []

    for k in range(N_total):
        Q = K.copy()

        #choose one of the non-zero elements to change
        r1 = np.random.randint(0,N-1)
        if r1 == 0:
            Q_ab = Q[r1,r1+1]
            N_ab = Nhit[r1,r1-1]
        else :
            r2 = np.random.randint(0,1)
            if r2 == 0:
                Q_ab = Q[r1,r1-1]
                N_ab = Nhit[r1,r1-1]
            else :
                Q_ab = Q[r1,r1+1]
                N_ab = Nhit[r1,r1+1]


        delta = np.random.exponential(scale=Q_ab) - Q_ab


        log_pacc = N_ab*np.log((Q_ab + delta)/Q_ab) - delta * t[r1]*np.sum(Nhit[r1])

        r = np.random.uniform(low=0.0,high=1.0)

        if np.log(r) < log_pacc :  #accept
            Q[r1,r1] -= delta
            if r1 == 0:
                Q[r1,r1+1] += delta
            else :
                if r2 == 0 :
                    Q[r1,r1-1] += delta

                else :
                    Q[r1,r1+1] += delta


        #only include after "interval" steps
        if (k+1)%interval == 0:
            Q_list.append(Q)

    #convert from list to array before returning
    return np.array(Q_list)
```
