## Supplementary material for "Markovian Weighted Ensemble Milestoning (M-WEM): Long-time Kinetics from Short Trajectories": MWEM-main.zip: test_mwem.py

```
"""
Unit and regression test for the mwem package.
"""

# Import package, test suite, and other packages as needed
import mwem
import pytest
import sys

def test_mwem_imported():
    """Sample test, will always pass so long as import statement worked"""
    assert "mwem" in sys.modules
```
