## Supplementary material for "Markovian Weighted Ensemble Milestoning (M-WEM): Long-time Kinetics from Short Trajectories": MWEM-main.zip: westpa_analysis_functions.py

```
import numpy as np
import pickle

def pruning(traj,low,high,index):
    '''
    traj : one traced trajectory
    low : lower value of the collective variable (milestone postion)
    high : higher value of the collective variable (milestone postion)
    index : 0 based index of the RC among the stored progress coordinates
    '''
    prunned = []
    for i in range(len(traj)):

        if traj[i,index] > low and traj[i,index] < high:
            prunned.append(traj[i,index])

        elif traj[i,index] > high and traj[i-1,index] < high:
            prunned.append(traj[i,index])

        elif traj[i,index] < low and traj[i-1,index] > low:
            prunned.append(traj[i,index])

        else :
            pass

    return np.array(prunned)

def remove_extra(traj,tau,num_iterations,ndim=2):
    '''
    traj : trajectory traced
    '''
    traj = traj.T

    traj2 = np.zeros((ndim,(tau-1)*num_iterations))
    for i in range(ndim):
        traj2[i] = np.delete(traj[i], np.arange(0, traj[i].size, tau))
    traj2 = traj2.T

    return traj2

def compute_crossings(pruned_traj,low,high):
    '''
    pruned_traj : pruned trajectory
    '''
    #print 1 for crossing right milestone
    #print 0 for crossing left milestone

    crossings = []
    for i in range(len(pruned_traj)-1):
        if pruned_traj[i] < high and pruned_traj[i+1] > high:
            crossings.append([i+1,1])
        elif pruned_traj[i] > low and pruned_traj[i+1] < low:
            crossings.append([i+1,0])
    return crossings
```
