## Supplementary material for "Markovian Weighted Ensemble Milestoning (M-WEM): Long-time Kinetics from Short Trajectories": MWEM-main.zip: README.md

MWEM
==============================
[//]: # (Badges)
[![GitHub Actions Build Status](https://github.com/REPLACE_WITH_OWNER_ACCOUNT/mwem/workflows/CI/badge.svg)](https://github.com/REPLACE_WITH_OWNER_ACCOUNT/mwem/actions?query=workflow%3ACI)
[![codecov](https://codecov.io/gh/REPLACE_WITH_OWNER_ACCOUNT/MWEM/branch/master/graph/badge.svg)](https://codecov.io/gh/REPLACE_WITH_OWNER_ACCOUNT/MWEM/branch/master)


Markovian Weighted Ensemble Milestoning (M-WEM)

Relevant manuscript:  "Markovian Weighted Ensemble Milestoning (M-WEM): Long-time Kinetics from Short Trajectories"
Dhiman Ray, Sharon E. Stone, Ioan Andricioaei, 
BioRxiv, doi: https://doi.org/10.1101/2021.06.26.450057 

### Copyright

Copyright (c) 2021, Dhiman Ray


#### Acknowledgements
 
Project based on the 
[Computational Molecular Science Python Cookiecutter](https://github.com/molssi/cookiecutter-cms) version 1.5.
