## Supplementary material for "Markovian Weighted Ensemble Milestoning (M-WEM): Long-time Kinetics from Short Trajectories": MWEM-main.zip: milestone_analysis_committor.py

```
import numpy as np
import pickle
import scipy.stats as st
import mwem

milestones = [-80.0, -60.0, -40.0, -20.0, 0.0, 20.0, 40.0, 60.0, 80.0]

indices = [0,1,2,3,4,5,6,7]


num_iter_list = [ 2, 4, 6, 8, 10, 20, 30, 40, 50, 60, 70, 80, 90, 100]
#f_mfpt = open('MFPT_convergence.dat','w')
#f3 = open('convergence_no_err.dat','w')
#f4 = open('K_i_i-1.dat','w')
#print('#iteration   #mean MFPT (ms) #confidence interval(low) #confidence interval(high)',file=f_mfpt)

for num_iter in num_iter_list:
    if num_iter == 100:
        crossings_file = 'crossings.pkl'
        weights_file = 'weights.txt'
    else :
        crossings_file = 'crossings_%d.pkl'%num_iter
        weights_file = 'weights_%d.txt'%num_iter

    MFPT, mean_MFPT, lower_conf, upper_conf, k_rev, N_i_j, R_i = mwem.milestoning(crossings_file=crossings_file, weights_file=weights_file, indices=indices, milestones=milestones,
        cutoff=1E-8, dt=10*1E-3, start_milestone = 1, end_milestone = len(milestones)-2, radial=False, cell_prob_file='cell_probability/cell_prob_%d.dat'%num_iter, numMCMC = 50000, returnNR=True,
        intervalMCMC = 50)

#    print(num_iter,mean_MFPT, lower_conf, upper_conf, file=f_mfpt)

#    print(num_iter,MFPT,file=f3)

#    print(num_iter,k_rev,file=f4)

#    print("Iterartion ",num_iter," done")

    np.savetxt('N_i_j_files/N_i_j_%d.dat'%num_iter,N_i_j)


#f_mfpt.close()
#f3.close()
#f4.close()
```
