## Supplementary material for "Markovian Weighted Ensemble Milestoning (M-WEM): Long-time Kinetics from Short Trajectories": MWEM-main.zip: analysis.py

```
#Code to generate the pruned trajectories and crossings data from WESTPA output file (west.h5)
#should be executed in the environment where WESTPA is installed


from matplotlib import pyplot as plt
import numpy as np
import pickle

np.set_printoptions(threshold=np.inf)
import w_ipa


w = w_ipa.WIPI()
# At startup, it will load or run the analysis schemes specified in the configuration file (typically west.cfg)
w.main()

def pruning(traj,low,high,index):
    '''
    traj : one traced trajectory
    low : lower value of the collective variable (milestone postion)
    high : higher value of the collective variable (milestone postion)
    index : 0 based index of the RC among the stored progress coordinates
    '''
    prunned = []
    for i in range(len(traj)):

        if traj[i,index] > low and traj[i,index] < high:
            prunned.append(traj[i,index])

        elif traj[i,index] > high and traj[i-1,index] < high:
            prunned.append(traj[i,index])

        elif traj[i,index] < low and traj[i-1,index] > low:
            prunned.append(traj[i,index])

        else :
            pass

    return np.array(prunned)

def remove_extra(traj,tau,ndim=2):
    '''
    traj : trajectory traced
    '''
    traj = traj.T

    traj2 = np.zeros((ndim,(tau-1)*w.niters))
    for i in range(ndim):
        traj2[i] = np.delete(traj[i], np.arange(0, traj[i].size, tau))
    traj2 = traj2.T

    return traj2

def compute_crossings(pruned_traj,low,high):
    '''
    pruned_traj : pruned trajectory
    '''
    #print 1 for crossing right milestone
    #print 0 for crossing left milestone

    crossings = []
    for i in range(len(pruned_traj)-1):
        if pruned_traj[i] < high and pruned_traj[i+1] > high:
            crossings.append([i+1,1])
        elif pruned_traj[i] > low and pruned_traj[i+1] < low:
            crossings.append([i+1,0])
    return crossings


tau = 101  #pcoord length from westpa adaptive.py

w.iteration = w.niters
final_trajectories = w.current.seg_id

weights = np.array(w.current.weights)
np.savetxt('weights.txt',weights)
#print(weights)
trajectories = []

for j in range(len(final_trajectories)):
    #trace the trajectory backwards
    traj_trace = w.trace(final_trajectories[j])
    xy = np.array(traj_trace['pcoord'])

    #remove extra frames appearing due to repeating the last frame of previous segment
    traj = remove_extra(xy,tau)

    trajectories.append(traj)

#save the trajectories
with open('trajectories.pkl','wb') as f1:
    pickle.dump(trajectories,f1)


#pruning the trajectories
ylow  = -80.0
yhigh = -60.0

pruned_trajectories = []
for j in range(len(trajectories)):
    pruned_trajectories.append(pruning(trajectories[j],ylow,yhigh,index=0))

#print(pruned_trajectories[54].shape)
#save the pruned trajectories
with open('pruned_trajectories.pkl','wb') as f1:
    pickle.dump(pruned_trajectories,f1)


crossings = []
for j in range(len(pruned_trajectories)):
    crossings.append(compute_crossings(pruned_trajectories[j],ylow,yhigh))

#save the crossings
with open('crossings.pkl','wb') as f1:
    pickle.dump(crossings,f1)
```
