## Supplementary material for "Markovian Weighted Ensemble Milestoning (M-WEM): Long-time Kinetics from Short Trajectories": MWEM-main.zip: analyze_all_convergence.sh

```
#!/bin/bash

for i in {0..9}
do
	cd cell_$i
	for j in 30 40 50 60 70 80 90 100 110 120 130 140 150 160 170 180 190 220 240 260 280 300
	do
		cp analysis.py analysis_$j.py
		sed -i "s/w.niters/$j/g" analysis_$j.py
		sed -i "s/weights.txt/weights_$j.txt/g" analysis_$j.py
		sed -i "s/trajectories.pkl/trajectories_$j.pkl/g" analysis_$j.py
		sed -i "s/trajectories_pruned.pkl/trajectories_pruned_$j.pkl/g" analysis_$j.py
		sed -i "s/crossings.pkl/crossings_$j.pkl/g" analysis_$j.py
		python analysis_$j.py
	done
	cd ..
done

cd cell_2.5
for j in 30 40 50 60 70 80 90 100 110 120 130 140 150 160 170 180 190 220 240 260 280 300
do
	cp analysis.py analysis_$j.py
	sed -i "s/w.niters/$j/g" analysis_$j.py
	sed -i "s/weights.txt/weights_$j.txt/g" analysis_$j.py
	sed -i "s/trajectories.pkl/trajectories_$j.pkl/g" analysis_$j.py
	sed -i "s/trajectories_pruned.pkl/trajectories_pruned_$j.pkl/g" analysis_$j.py
	sed -i "s/crossings.pkl/crossings_$j.pkl/g" analysis_$j.py
	python analysis_$j.py
done
cd ..


cd cell_3.5
for j in 30 40 50 60 70 80 90 100 110 120 130 140 150 160 170 180 190 220 240 260 280 300
do
        cp analysis.py analysis_$j.py
        sed -i "s/w.niters/$j/g" analysis_$j.py
        sed -i "s/weights.txt/weights_$j.txt/g" analysis_$j.py
        sed -i "s/trajectories.pkl/trajectories_$j.pkl/g" analysis_$j.py
        sed -i "s/trajectories_pruned.pkl/trajectories_pruned_$j.pkl/g" analysis_$j.py
        sed -i "s/crossings.pkl/crossings_$j.pkl/g" analysis_$j.py
        python analysis_$j.py
done
cd ..
```
