## Supplementary material for "Markovian Weighted Ensemble Milestoning (M-WEM): Long-time Kinetics from Short Trajectories": MWEM-main.zip: computational_cost.py

```
import numpy as np
import pickle

indices = [0,1,2,2.5,3,3.5,4,5,6,7,8,9]

cost = 0.0

for i in range(len(indices)):
    a = pickle.load(open('cell_%s/trajectories_300.pkl'%str(indices[i]),'rb'))


    len_traj = len(a[0])


    #patch all trajectories one after another
    trajs_stacked = [j for k in a for j in k]
    
    cost_cell = len(trajs_stacked)*20*1E-6

    print("cell_%s cost : "%str(indices[i]), cost_cell, 'ns')

    cost += cost_cell


print("Total computational cost = ",cost,"ns")
```
