## Supplementary material for "Markovian Weighted Ensemble Milestoning (M-WEM): Long-time Kinetics from Short Trajectories": MWEM-main.zip: mean_krev_200.py

```
import numpy as np

l = np.loadtxt('K_i_i-1.dat')
a = l.T[1]


alpha = 0.01373 #obtained from fraction coverage

#D = 1E-5 #approximate value for small molecule in water (in cm^2/s)

r = 12 #radius of the outermost milestone in Angstrom

a = 7.569 * 1E+8 * r * alpha * a

mean_krev = np.mean(a[-10:-5])
std_krev = np.std(a[-10:-5])

conf_int = 2.571*std_krev/np.sqrt(5.0)

print('k_on = ',mean_krev,'+/-',conf_int)
```
