## Supplementary material for "Markovian Weighted Ensemble Milestoning (M-WEM): Long-time Kinetics from Short Trajectories": MWEM-main.zip: milestone_analysis_convergence.py

```
import numpy as np
import pickle
import scipy.stats as st
import mwem

milestones = [1.0,1.5,2.0,2.5,3.0,3.5,4.0,5.0,6.0,8.0,10.0,12.0,13.0]

indices = [0,1,2,2.5,3,3.5,4,5,6,7,8,9] 


num_iter_list = [ 30, 40, 50, 60, 70, 80, 90, 100, 110, 120, 130, 140, 150, 160, 170, 180, 190, 200, 220, 240, 260, 280, 300]
#f_mfpt = open('MFPT_convergence.dat','w')
#f3 = open('convergence_no_err.dat','w')
#f4 = open('K_i_i-1.dat','w')
#print('#iteration   #mean MFPT (ms) #confidence interval(low) #confidence interval(high)',file=f_mfpt)

for num_iter in num_iter_list:
    if num_iter == 200:
        crossings_file = 'crossings.pkl'
        weights_file = 'weights.txt'
    else :
        crossings_file = 'crossings_%d.pkl'%num_iter
        weights_file = 'weights_%d.txt'%num_iter

    MFPT, mean_MFPT, lower_conf, upper_conf, k_rev, N_i_j, R_i = mwem.milestoning(crossings_file=crossings_file, weights_file=weights_file, indices=indices, milestones=milestones, 
        cutoff=1E-8, dt=20.0*1E-12, start_milestone = 0, end_milestone = len(milestones)-1, radial=True, cell_prob_file='cell_probability/cell_prob_%d.dat'%num_iter,returnNR=True, numMCMC = 5000, intervalMCMC = 50)

#    print(num_iter,mean_MFPT, lower_conf, upper_conf, file=f_mfpt)

#    print(num_iter,MFPT,file=f3)

#    print(num_iter,k_rev,file=f4)

#    print("Iterartion ",num_iter," done")

    #f5 = open('R_i/R_i_%d.dat'%num_iter,'w')
    #for lifetime in R_i:
    #    print(lifetime,file=f5)
    #f5.close()

    np.savetxt('N_i_j_files/N_i_j_%d.dat'%num_iter,N_i_j)
    #for i in range(len(N_i_j)):
    #    for j in ra
    #    print(lifetime,file=f6)
    #f6.close()


#f_mfpt.close()
#f3.close()
#f4.close()
```
