## Supplementary material for "Markovian Weighted Ensemble Milestoning (M-WEM): Long-time Kinetics from Short Trajectories": MWEM-main.zip: build.py

```
import numpy as np
import os
import subprocess

milestones = [-80.0, -60.0, -40.0, -20.0, 0.0, 20.0, 40.0, 60.0, 80.0]

for i in range(len(milestones)-1):

    left = milestones[i]
    right = milestones[i+1]

    middle = 0.5*(left + right)

    dir_name = 'cell_%d'%i

    os.system('cp -r template %s'%dir_name)

    subprocess.call(["sed -i 's/CENTER/%0.2f/g' %s/equilibration/colvars.in"%(middle,dir_name)], shell=True)

    subprocess.call(["sed -i 's/LOW/%0.2f/g' %s/common_files/colvars.in"%(left,dir_name)], shell=True)
    subprocess.call(["sed -i 's/HIGH/%0.2f/g' %s/common_files/colvars.in"%(right,dir_name)], shell=True)
```
