## Supplementary material for "Markovian Weighted Ensemble Milestoning (M-WEM): Long-time Kinetics from Short Trajectories": MWEM-main.zip: adaptive.py

```
from __future__ import print_function, division

import numpy

from west.propagators import WESTPropagator

from west.systems import WESTSystem

from westpa.binning import RectilinearBinMapper

from westpa.binning import FuncBinMapper

from westpa.binning import RecursiveBinMapper

import logging

log = logging.getLogger('westpa.rc')

PI = numpy.pi

from numpy import *

pcoord_dtype = numpy.float32

#THESE ARE THE PARAMETERS YOU CAN CHANGE

bintargetcount=4 #number of walkers per bin

numberofdim=2  # number of dimensions

binsperdim=[5,5]   # You will have prod(binsperdim)+numberofdim*(2+2*splitIsolated)+activetarget bins total

pcoordlength=101 # length of the pcoord

maxcap=[inf,inf]	#for each dimension enter the maximum number at which binning can occur, if you do not wish to have a cap use inf

mincap=[-inf,-inf]  #for each dimension enter the minimum number at which binning can occur, if you do not wish to have a cap use -inf

targetstate=[None]    #enter boundaries for target state or None if there is no target state in that dimension

targetstatedirection=[None]  #if your target state is meant to be greater that the starting pcoor use 1 or else use -1. This will be done for each dimension in your simulation

activetarget=0		#if there is no target state make this zero

splitIsolated=1     #choose 0 to disable the use of bottleneck walkers (not recomended)

#########

def function_map(coords, mask, output):

	splittingrelevant=True #This is to make sure splitting is relevant (not relevant for binner after recycling for example)
	
	originalcoords=copy(coords) #It is a good idea to keep an original array

	maxlist=[] #Preparing array to contain maximum pcoords in each dimension

	minlist=[] #Preparing array to contain minimum pcoords in each dimension

	difflist=[] #Preparing array to contain "bottleneck" values in positive direction for each dimensio #Preparing array to contain "bottleneck" values in negative direction for each dimensionn

	flipdifflist=[] #Preparing array to contain "bottleneck" values in negative direction for each dimension

	for n in range(numberofdim): #going through each dimension

		try:    #because binning should be handled different for recycled trajectories we load in a binbounds.txt created after an iteration completes

			extremabounds=loadtxt('binbounds.txt') 

			currentmax=amax(extremabounds[:,n])

			currentmin=amin(extremabounds[:,n])

		except: #during initialization this may not exitst so use current coords for extrema

			currentmax=amax(coords[:,n])

			currentmin=amin(coords[:,n])

		if maxcap[n]<currentmax: #Checking the maxcap in each dimension

                	currentmax=maxcap[n]

		if mincap[n]>currentmin: #Checking the mincap in each dimension

                	currentmin=mincap[n]

		maxlist.append(currentmax) #Need arrays for our extrema since there may be multiple depending on number of dimension

		minlist.append(currentmin)

		try:	#Recycled trajectories should not be tagged and will throw exception to be handled by except statement

			temp=column_stack((coords[:,n],coords[:,numberofdim])) #Create an array containing progress coordinates of one dimension and associated probailities

			temp=temp[temp[:,0].argsort()] #Sort this by progress coordinate

			for p in range(len(temp)):  #This just deals with the fact that currently received probailities are in float32 (it is probably best to disregard probailities smaller than E-39 anyway for tagging

                		if temp[p][1]==0:

                			temp[p][1]=10**-39

			fliptemp=flipud(temp) #Recived sorted array in opposite direction

			difflist.append(0) #Provide starting minimum of 0 (in very unlikely case of pcoord 0 and no tagged this could cause arbitrary tag (very minor impact), work to fix)

			flipdifflist.append(0)	

			maxdiff=0 

			flipmaxdiff=0

			for i in range(1,len(temp)-1): #calculating of the "bottleneck" values, we need to sum all of the probability past a potential "bottleneck"

				comprob=0

				flipcomprob=0

				j=i+1

				while j<len(temp): #calculating the cumulative probability past the each walker in each direction

					comprob=comprob+temp[j][1]

					flipcomprob=flipcomprob+fliptemp[j][1]

					j=j+1

				if temp[i][0]<maxcap[n] and temp[i][0]>mincap[n]:

					if (-log(comprob)+log(temp[i][1]))>maxdiff: #we want to find the point where the difference between the walker and the cumulative probability past it is at a maximum, we use logarithms to compare differences

						difflist[n]=temp[i][0]

						maxdiff=-log(comprob)+log(temp[i][1])

				if fliptemp[i][0]<maxcap[n] and fliptemp[i][0]>mincap[n]:

					if (-log(flipcomprob)+log(fliptemp[i][1]))>flipmaxdiff:

						flipdifflist[n]=fliptemp[i][0]

						flipmaxdiff=-log(flipcomprob)+log(fliptemp[i][1])

		except:

			splittingrelevant=False  #if an error is thrown tagging of bottleneck walkers is not needed

	for i in range(len(output)): #this section deals with proper assignment of walkers to bins

		binnumber=2*numberofdim #essentially the bin number 

		for n in range(numberofdim):

			if (activetarget==1) and targetstate[n] is not None:

				if (originalcoords[i,n]*targetstatedirection[n]) >= (targetstate[n]*targetstatedirection[n]): #if the target state has been reached assign to following bin

					binnumber=prod(binsperdim)+numberofdim*2

			if (binnumber==prod(binsperdim)+numberofdim*2): #this ends the loop if binned in target state, n= numberofdim should not go in above line because of elif statements 

				n=numberofdim

			elif coords[i,n]>=maxlist[n] or originalcoords[i,n]>=maxcap[n]: #assign maxima or those over max cap to own bin

				binnumber= 2*n

				n=numberofdim

			elif coords[i,n]<=minlist[n] or originalcoords[i,n]<=mincap[n]: #assign minima or those under minima to own bin

				binnumber =2*n+1

				n=numberofdim

			elif splittingrelevant and coords[i,n]==difflist[n] and splitIsolated==1: #assign bottleneck walker in one direction to bin

				binnumber=prod(binsperdim)+numberofdim*2+2*n+activetarget

				n=numberofdim

			elif splittingrelevant and coords[i,n]==flipdifflist[n] and splitIsolated==1: #assign bottleneck walker in other direction to bin

				binnumber=prod(binsperdim)+numberofdim*2+2*n+activetarget+1

				n=numberofdim

		if binnumber==2*numberofdim: #calculate  binning for evenly spaced bins

			for j in range(numberofdim):

				binnumber = binnumber + (digitize(coords[i][j],linspace(minlist[j],maxlist[j],binsperdim[j]+1))-1)*prod(binsperdim[0:j])

		output[i]=binnumber

	return output

#######
	
class System(WESTSystem): #class initialization

	def initialize(self):

		self.pcoord_ndim = numberofdim

		self.pcoord_len = pcoordlength

		self.pcoord_dtype = numpy.float32 

		self.bin_mapper = FuncBinMapper(function_map, prod(binsperdim)+numberofdim*(2+2*splitIsolated)+activetarget) #Changed binsperbin to binsperdim
 
		self.bin_target_counts = numpy.empty((self.bin_mapper.nbins,), numpy.int_)

		self.bin_target_counts[...] = bintargetcount
```
