## Supplementary material for "Markovian Weighted Ensemble Milestoning (M-WEM): Long-time Kinetics from Short Trajectories": MWEM-main.zip: driver.py

```
import logging
log = logging.getLogger(__name__)
import numpy
import operator
from math import ceil
import random


import westpa
from west import Segment

class ConsistencyError(RuntimeError):
    pass

class AccuracyError(RuntimeError):
    pass

class NewWeightEntry:
    NW_SOURCE_RECYCLED = 0
    
    def __init__(self, source_type, weight, prev_seg_id=None,
                 prev_init_pcoord=None, prev_final_pcoord=None, new_init_pcoord=None,
                 target_state_id = None, initial_state_id = None):
        self.source_type = source_type
        self.weight = weight
        self.prev_seg_id = prev_seg_id
        self.prev_init_pcoord = numpy.asarray(prev_init_pcoord) if prev_init_pcoord is not None else None
        self.prev_final_pcoord = numpy.asarray(prev_final_pcoord) if prev_final_pcoord is not None else None 
        self.new_init_pcoord = numpy.asarray(new_init_pcoord) if new_init_pcoord is not None else None
        self.target_state_id = target_state_id        
        self.initial_state_id = initial_state_id
        
    def __repr__(self):
        return ('<{} object at 0x{:x}: weight={self.weight:g} target_state_id={self.target_state_id} prev_final_pcoord={self.prev_final_pcoord}>'
                .format(self.__class__.__name__, id(self), self=self))

class WEDriver:
    '''A class implemented Huber & Kim's weighted ensemble algorithm over Segment objects.
    This class handles all binning, recycling, and preparation of new Segment objects for the
    next iteration. Binning is accomplished using system.bin_mapper, and per-bin target counts
    are from system.bin_target_counts. 
    
    The workflow is as follows:
    
      1) Call `new_iteration()` every new iteration, providing any recycling targets that are
         in force and any available initial states for recycling.
      2) Call `assign()` to assign segments to bins based on their initial and end points. This
         returns the number of walkers that were recycled.
      3) Call `run_we()`, optionally providing a set of initial states that will be used to
         recycle walkers.
         
    Note the presence of flux_matrix, transition_matrix,
    current_iter_segments, next_iter_segments, recycling_segments,
    initial_binning, final_binning, next_iter_binning, and new_weights (to be documented soon).
    '''
    
    weight_split_threshold = 2.0
    weight_merge_cutoff = 1.0
        
    def __init__(self, rc=None, system=None):
        self.rc = rc or westpa.rc
        self.system = system or self.rc.get_system_driver()
        
        # Whether to adjust counts to exactly match target count
        self.do_adjust_counts = True 
        
        # bin mapper and per-bin target counts (see new_iteration for initialization)
        self.bin_mapper = None
        self.bin_target_counts = None
        
        # Mapping of bin index to target state
        self.target_states = None 
        
        # binning on initial points
        self.initial_binning = None
        
        # binning on final points (pre-WE)
        self.final_binning   = None
                                
        # binning on initial points for next iteration
        self.next_iter_binning = None
                
        # Flux and rate matrices for the current iteration
        self.flux_matrix = None
        self.transition_matrix = None
        
        # Information on new weights (e.g. from recycling) for the next iteration
        self.new_weights = None
        
        # Set of initial states passed to run_we() that are actually used for
        # recycling targets
        self.used_initial_states = None
        
        self.avail_initial_states = None
        
        self.process_config()
    
    def process_config(self):
        config = self.rc.config
        
        config.require_type_if_present(['west', 'we', 'adjust_counts'], bool)
        
        self.do_adjust_counts = config.get(['west', 'we', 'adjust_counts'], True)
        log.info('Adjust counts to exactly match target_counts: {}'.format(self.do_adjust_counts))
        
        self.weight_split_threshold = config.get(['west', 'we', 'weight_split_threshold'], self.weight_split_threshold)
        log.info('Split threshold: {}'.format(self.weight_split_threshold))
        
        self.weight_merge_cutoff = config.get(['west', 'we', 'weight_merge_cutoff'], self.weight_merge_cutoff)
        log.info('Merge cutoff: {}'.format(self.weight_merge_cutoff))
        
        
    @property
    def next_iter_segments(self):
        '''Newly-created segments for the next iteration'''
        if self.next_iter_binning is None:
            raise RuntimeError('cannot access next iteration segments before running WE')
        
        for _bin in self.next_iter_binning:
            for walker in _bin:
                yield walker
                
    @property
    def current_iter_segments(self):
        '''Segments for the current iteration'''
        for _bin in self.final_binning:
            for walker in _bin:
                yield walker
                
    @property
    def next_iter_assignments(self):
        '''Bin assignments (indices) for initial points of next iteration.'''
        if self.next_iter_binning is None:
            raise RuntimeError('cannot access next iteration segments before running WE')
        
        for ibin, _bin in enumerate(self.next_iter_binning):
            for _walker in _bin:
                yield ibin
                
    @property
    def current_iter_assignments(self):
        '''Bin assignments (indices) for endpoints of current iteration.'''
        for ibin,_bin in enumerate(self.final_binning):
            for walker in _bin:
                yield ibin
                
    @property
    def recycling_segments(self):
        '''Segments designated for recycling'''
        if len(self.target_states):
            for (ibin,tstate) in self.target_states.items():
                for segment in self.final_binning[ibin]:
                    yield segment                
        else:
            return
    
    @property
    def n_recycled_segs(self):
        '''Number of segments recycled this iteration'''
        count = 0
        for _segment in self.recycling_segments:
            count += 1
        return count
    
    @property
    def n_istates_needed(self):
        '''Number of initial states needed to support recycling for this iteration'''
        n_istates_avail = len(self.avail_initial_states)
        return max(0, self.n_recycled_segs - n_istates_avail)
                
    def clear(self):
        '''Explicitly delete all Segment-related state.'''
        
        del self.initial_binning, self.final_binning, self.next_iter_binning
        del self.flux_matrix, self.transition_matrix
        del self.new_weights, self.used_initial_states, self.avail_initial_states
        
        self.initial_binning = None
        self.final_binning = None
        self.next_iter_binning = None
        self.flux_matrix = None
        self.transition_matrix = None
        self.avail_initial_states = None
        self.used_initial_states = None
        self.new_weights = None
        
    def new_iteration(self, initial_states=None, target_states=None, new_weights=None, bin_mapper=None, bin_target_counts=None):
        '''Prepare for a new iteration. ``initial_states`` is a sequence of all InitialState objects valid
        for use in to generating new segments for the *next* iteration (after the one being begun with the call to
        new_iteration); that is, these are states available to recycle to. Target states which generate recycling events
        are specified in ``target_states``, a sequence of TargetState objects. Both ``initial_states`` 
        and ``target_states`` may be empty as required.
        
        The optional ``new_weights`` is a sequence of NewWeightEntry objects which will 
        be used to construct the initial flux matrix.
        
        The given ``bin_mapper`` will be used for assignment, and ``bin_target_counts`` used for splitting/merging
        target counts; each will be obtained from the system object if omitted or None.        
        '''
        
        self.clear()
        
        new_weights = new_weights or []
        if initial_states is None:
            initial_states = initial_states or []
        
        # update mapper, in case it has changed on the system driver and has not been overridden
        if bin_mapper is not None:
            self.bin_mapper = bin_mapper
        else:
            self.bin_mapper = self.system.bin_mapper
            
        if bin_target_counts is not None:
            self.bin_target_counts = bin_target_counts
        else:
            self.bin_target_counts = numpy.array(self.system.bin_target_counts).copy()
        nbins = self.bin_mapper.nbins                
        log.debug('mapper is {!r}, handling {:d} bins'.format(self.bin_mapper, nbins))
                
        self.initial_binning    = self.bin_mapper.construct_bins()
        self.final_binning      = self.bin_mapper.construct_bins()
        self.next_iter_binning  = None
        
        flux_matrix = self.flux_matrix = numpy.zeros((nbins,nbins), dtype=numpy.float64)
        transition_matrix = self.transition_matrix = numpy.zeros((nbins,nbins), numpy.uint)
        
        # map target state specifications to bins
        target_states = target_states or []
        self.target_states = {}
        for tstate in target_states:
            tstate_assignment = self.bin_mapper.assign([tstate.pcoord])[0]
            self.target_states[tstate_assignment] = tstate
            log.debug('target state {!r} mapped to bin {}'.format(tstate, tstate_assignment))
            self.bin_target_counts[tstate_assignment] = 0
            
        # loop over recycled segments, adding entries to the flux matrix appropriately
        if new_weights:
            init_pcoords = numpy.empty((len(new_weights), self.system.pcoord_ndim), dtype=self.system.pcoord_dtype)
            prev_init_pcoords = numpy.empty((len(new_weights), self.system.pcoord_ndim), dtype=self.system.pcoord_dtype)
            
            for (ientry,entry) in enumerate(new_weights):
                init_pcoords[ientry] = entry.new_init_pcoord
                prev_init_pcoords[ientry] = entry.prev_init_pcoord
            
            init_assignments = self.bin_mapper.assign(init_pcoords)
            prev_init_assignments = self.bin_mapper.assign(prev_init_pcoords)
            
            for (entry, i, j) in zip(new_weights, prev_init_assignments, init_assignments):
                flux_matrix[i,j] += entry.weight
                transition_matrix[i,j] += 1
                
            del init_pcoords, prev_init_pcoords, init_assignments, prev_init_assignments
        
        self.avail_initial_states = {state.state_id: state for state in initial_states}
        self.used_initial_states = {}
        
    def add_initial_states(self, initial_states):
        '''Add newly-prepared initial states to the pool available for recycling.'''
        for state in initial_states:
            self.avail_initial_states[state.state_id] = state
            
    @property
    def all_initial_states(self):
        '''Return an iterator over all initial states (available or used)'''
        for state in self.avail_initial_states.values():
            yield state
        for state in self.used_initial_states.values():
            yield state
                
    def assign(self, segments, initializing=False, endprop=False, recycling=False):
        '''Assign segments to initial and final bins, and update the (internal) lists of used and available
        initial states. If ``initializing`` is True, then the "final" bin assignments will
        be identical to the initial bin assignments, a condition required for seeding a new iteration from
        pre-existing segments.'''

        # collect initial and final coordinates into one place        
        all_pcoords = numpy.empty((2,len(segments), self.system.pcoord_ndim+1), dtype=self.system.pcoord_dtype)
        if recycling:
            for iseg, segment in enumerate(segments):
                all_pcoords[0,iseg] = numpy.append(segment.pcoord[0,:],-1)
                all_pcoords[1,iseg] = numpy.append(segment.pcoord[-1,:],-1)
        else:
            for iseg, segment in enumerate(segments):
                all_pcoords[0,iseg] = numpy.append(segment.pcoord[0,:],segment.weight)
                all_pcoords[1,iseg] = numpy.append(segment.pcoord[-1,:],segment.weight)
        if endprop:
                numpy.savetxt('binbounds.txt',all_pcoords[1,:,:])
        # assign based on initial and final progress coordinates
        initial_assignments = self.bin_mapper.assign(all_pcoords[0,:,:])
        if initializing:
            final_assignments = initial_assignments
        else:
            final_assignments = self.bin_mapper.assign(all_pcoords[1,:,:])
        initial_binning = self.initial_binning
        final_binning = self.final_binning
        flux_matrix = self.flux_matrix
        transition_matrix = self.transition_matrix
        for (segment,iidx,fidx) in zip(segments, initial_assignments, final_assignments):
            initial_binning[iidx].add(segment)
            final_binning[fidx].add(segment)
            flux_matrix[iidx,fidx] += segment.weight
            transition_matrix[iidx,fidx] += 1
            
        n_recycled_total = self.n_recycled_segs
        n_new_states = n_recycled_total - len(self.avail_initial_states)
        
        log.debug('{} walkers scheduled for recycling, {} initial states available'.format(n_recycled_total, 
                                                                                           len(self.avail_initial_states)))
        
        if n_new_states > 0:
            return n_new_states
        else:
            return 0
    
    def _recycle_walkers(self):
        '''Recycle walkers'''
        
        # recall that every walker we deal with is already a new segment in the subsequent iteration,
        # so to recycle, we actually move the appropriate Segment from the target bin to the initial state bin
        
        self.new_weights = []
        
        n_recycled_walkers = len(list(self.recycling_segments))
        if not n_recycled_walkers:
            return
        elif n_recycled_walkers > len(self.avail_initial_states):
            raise ConsistencyError('need {} initial states for recycling, but only {} present'
                                   .format(n_recycled_walkers,len(self.avail_initial_states)))

        used_istate_ids = set()
        istateiter = iter(self.avail_initial_states.values())
        for (ibin,target_state) in self.target_states.items():
            target_bin = self.next_iter_binning[ibin]
            for segment in set(target_bin):
                initial_state = next(istateiter)                
                istate_assignment = self.bin_mapper.assign([initial_state.pcoord])[0]
                parent = self._parent_map[segment.parent_id]
                parent.endpoint_type = Segment.SEG_ENDPOINT_RECYCLED

                if log.isEnabledFor(logging.DEBUG):
                    log.debug('recycling {!r} from target state {!r} to initial state {!r}'.format(segment, target_state,
                                                                                                   initial_state))
                    log.debug('parent is {!r}'.format(parent))                
                
                
                segment.parent_id = -(initial_state.state_id+1)
                segment.pcoord[0] = initial_state.pcoord

                self.new_weights.append(NewWeightEntry(source_type=NewWeightEntry.NW_SOURCE_RECYCLED,
                                                       weight=parent.weight, prev_seg_id=parent.seg_id,
                                                       # the .copy() is crucial, otherwise the slice of pcoords will
                                                       # keep the parent segments' pcoord data alive unnecessarily long
                                                       prev_init_pcoord=parent.pcoord[0].copy(),
                                                       prev_final_pcoord=parent.pcoord[-1].copy(),
                                                       new_init_pcoord=initial_state.pcoord.copy(),
                                                       target_state_id=target_state.state_id,
                                                       initial_state_id=initial_state.state_id) )
                
                if log.isEnabledFor(logging.DEBUG):
                    log.debug('new weight entry is {!r}'.format(self.new_weights[-1]))

                self.next_iter_binning[istate_assignment].add(segment)
                
                initial_state.iter_used = segment.n_iter
                log.debug('marking initial state {!r} as used'.format(initial_state))
                used_istate_ids.add(initial_state.state_id)
                target_bin.remove(segment)
                 
            assert len(target_bin) == 0
            
        # Transfer newly-assigned states from "available" to "used"
        for state_id in used_istate_ids:
            self.used_initial_states[state_id] = self.avail_initial_states.pop(state_id)
            
                            
    def _split_walker(self, segment, m, bin):
        '''Split the walker ``segment`` (in ``bin``) into ``m`` walkers'''

        bin.remove(segment)
            
        new_segments = []
        for _inew in range(0,m):
            new_segment = Segment(n_iter = segment.n_iter, #previously incremented
                                  weight = segment.weight/m,
                                  parent_id = segment.parent_id,
                                  wtg_parent_ids = set(segment.wtg_parent_ids),
                                  pcoord = segment.pcoord.copy(),
                                  status = Segment.SEG_STATUS_PREPARED)
            new_segment.pcoord[0,:] = segment.pcoord[0,:]
            new_segments.append(new_segment)
            
        bin.update(new_segments)

        if log.isEnabledFor(logging.DEBUG):
            log.debug('splitting {!r} into {:d}:\n    {!r}'.format(segment, m, new_segments))
            
        return new_segments 
                        
    def _merge_walkers(self, segments, cumul_weight, bin):
        '''Merge the given ``segments`` in ``bin``, previously sorted by weight, into one conglomerate segment.
        ``cumul_weight`` is the cumulative sum of the weights of the ``segments``; this may be None to calculate here.'''
        
        if cumul_weight is None:
            cumul_weight = numpy.add.accumulate([segment.weight for segment in segments])
        
        glom = Segment(n_iter = segments[0].n_iter, # assumed correct (and equal among all segments)
                       weight = cumul_weight[len(segments)-1],
                       status = Segment.SEG_STATUS_PREPARED,
                       pcoord = self.system.new_pcoord_array(),
                       )
                
        # Select the history to use
        # The following takes a random number in the interval 0 <= x < glom.weight, then
        # sees where this value falls among the (sorted) weights of the segments being merged;
        # this ensures that a walker with (e.g.) twice the weight of its brethren has twice the
        # probability of having its history selected for continuation
        iparent = numpy.digitize((random.uniform(0,glom.weight),),cumul_weight)[0]
        gparent_seg = segments[iparent]
        
        # Inherit history from this segment ("gparent" stands for "glom parent", as opposed to historical
        # parent). 
        glom.parent_id = gparent_seg.parent_id
        glom.pcoord[0,:] = gparent_seg.pcoord[0,:]
        
        # Weight comes from all segments being merged, and therefore all their
        # parent segments
        glom.wtg_parent_ids = set()
        for segment in segments:
            glom.wtg_parent_ids |= segment.wtg_parent_ids
        
        # Remove merged walkers from consideration before treating initial states
        bin.difference_update(segments)            
            
        # The historical parent of gparent is continued; all others are marked as merged
        for segment in segments:
            if segment is gparent_seg:
                # we must ignore initial states here...
                if segment.parent_id >= 0:
                    self._parent_map[segment.parent_id].endpoint_type = Segment.SEG_ENDPOINT_CONTINUES
            else:
                # and "unuse" an initial state here (recall that initial states are in 1:1 correspondence
                # with the segments they initiate), except when a previously-split particle is being
                # merged
                if segment.parent_id >= 0:
                    self._parent_map[segment.parent_id].endpoint_type = Segment.SEG_ENDPOINT_MERGED
                else:
                    if segment.initial_state_id in {segment.initial_state_id for segment in bin}:
                        log.debug('initial state in use by other walker; not removing')
                    else:
                        initial_state = self.used_initial_states.pop(segment.initial_state_id)
                        log.debug('freeing initial state {!r} for future use (merged)'.format(initial_state))
                        self.avail_initial_states[initial_state.state_id] = initial_state
                        initial_state.iter_used = None

        if log.isEnabledFor(logging.DEBUG):
            log.debug('merging ({:d}) {!r} into 1:\n    {!r}'.format(len(segments), segments, glom))
                

        bin.add(glom)
        
    def _split_by_weight(self, ibin):
        '''Split overweight particles'''
        
        bin = self.next_iter_binning[ibin]
        target_count = self.bin_target_counts[ibin]
        segments = numpy.array(sorted(bin, key=operator.attrgetter('weight')), dtype=numpy.object_)
        weights = numpy.array(list(map(operator.attrgetter('weight'), segments)))
        ideal_weight = weights.sum() / target_count
 
        if len(bin) > 0:
            assert target_count > 0
               
        to_split = segments[weights > self.weight_split_threshold*ideal_weight]
        
        for segment in to_split:
            m = int(ceil(segment.weight / ideal_weight))
            self._split_walker(segment, m, bin)

    
    def _merge_by_weight(self, ibin):
        '''Merge underweight particles'''
        
        bin = self.next_iter_binning[ibin]
        target_count = self.bin_target_counts[ibin]
        weight = sum(map(operator.attrgetter('weight'), bin))
        target_count = self.bin_target_counts[ibin]
        ideal_weight = weight / target_count    
        
        while True:
            segments = numpy.array(sorted(bin, key=operator.attrgetter('weight')), dtype=numpy.object_)
            weights = numpy.array(list(map(operator.attrgetter('weight'), segments)))
            cumul_weight = numpy.add.accumulate(weights)
            
            to_merge = segments[cumul_weight <= ideal_weight*self.weight_merge_cutoff]
            if len(to_merge) < 2:
                return
            
            self._merge_walkers(to_merge, cumul_weight, bin)
    
    def _adjust_count(self, ibin):
        bin = self.next_iter_binning[ibin]
        target_count = self.bin_target_counts[ibin]
        weight_getter = operator.attrgetter('weight')

        # split        
        while len(bin) < target_count:
            log.debug('adjusting counts by splitting')
            # always split the highest probability walker into two
            segments = sorted(bin, key=weight_getter)
            self._split_walker(segments[-1], 2, bin)
            
        # merge
        while len(bin) > target_count:
            log.debug('adjusting counts by merging')
            # always merge the two lowest-probability walkers
            segments = sorted(bin, key=weight_getter)
            self._merge_walkers(segments[:2], cumul_weight=None, bin=bin)

    def _check_pre(self):
        for ibin, _bin in enumerate(self.next_iter_binning):
            if self.bin_target_counts[ibin] == 0 and len(_bin) > 0:
                raise ConsistencyError('bin {:d} has target count of 0 but contains {:d} walkers'.format(ibin, len(_bin)))

    def _check_post(self):
        for segment in self.next_iter_segments:
            if segment.weight == 0:
                raise ConsistencyError('segment {!r} has weight of zero')
            
    def _prep_we(self):
        '''Prepare internal state for WE recycle/split/merge.'''
        self._parent_map = {}
        self.next_iter_binning = self.bin_mapper.construct_bins()

    def _run_we(self):
        '''Run recycle/split/merge. Do not call this function directly; instead, use
        populate_initial(), rebin_current(), or construct_next().'''
        self._recycle_walkers()
        
        # sanity check
        self._check_pre()
        
        # Regardless of current particle count, always split overweight particles and merge underweight particles
        # Then and only then adjust for correct particle count
        for (ibin,bin) in enumerate(self.next_iter_binning):
            if len(bin) == 0: 
                continue
                        
            self._split_by_weight(ibin)
            self._merge_by_weight(ibin)
            if self.do_adjust_counts:
                self._adjust_count(ibin)
            
        self._check_post()
        
        self.new_weights = self.new_weights or []
        
        log.debug('used initial states: {!r}'.format(self.used_initial_states))
        log.debug('available initial states: {!r}'.format(self.avail_initial_states))

            
    def populate_initial(self, initial_states, weights, system=None):
        '''Create walkers for a new weighted ensemble simulation.
        
        One segment is created for each provided initial state, then binned and split/merged
        as necessary. After this function is called, next_iter_segments will yield the new
        segments to create, used_initial_states will contain data about which of the
        provided initial states were used, and avail_initial_states will contain data about
        which initial states were unused (because their corresponding walkers were merged
        out of existence).
        '''

        # This has to be down here to avoid an import race
        from west.data_manager import weight_dtype
        EPS = numpy.finfo(weight_dtype).eps                
                
        system = system or westpa.rc.get_system_driver()
        self.new_iteration(initial_states=[], target_states=[],
                           bin_mapper=system.bin_mapper, bin_target_counts=system.bin_target_counts)
        
        # Create dummy segments
        segments = []
        for (seg_id, (initial_state,weight)) in enumerate(zip(initial_states,weights)):
            dummy_segment = Segment(n_iter=0,
                                    seg_id=seg_id,
                                    parent_id=-(initial_state.state_id+1),
                                    weight=weight,
                                    wtg_parent_ids=set([-(initial_state.state_id+1)]),
                                    pcoord=system.new_pcoord_array(),
                                    status=Segment.SEG_STATUS_PREPARED)
            dummy_segment.pcoord[[0,-1]] = initial_state.pcoord
            segments.append(dummy_segment)
        
        # Adjust weights, if necessary
        tprob = sum(weights)
        if abs(1.0 - tprob) > len(weights) * EPS:
            pscale = 1.0/tprob
            log.warning('Weights of initial segments do not sum to unity; scaling by {:g}'.format(pscale))
            for segment in segments:
                segment.weight *= pscale
        
        self.assign(segments, initializing=True)
        self.construct_next()
        
        # We now have properly-constructed initial segments, except for parent information,
        # and we need to  mark initial states as used or unused
        istates_by_id = {state.state_id: state for state in initial_states}
        dummysegs_by_id = self._parent_map
        self.avail_initial_states = dict(istates_by_id)
        self.used_initial_states = {}
        for segment in self.next_iter_segments:
            segment.parent_id = dummysegs_by_id[segment.parent_id].parent_id
            segment.wtg_parent_ids=set([segment.parent_id])
            assert segment.initpoint_type == Segment.SEG_INITPOINT_NEWTRAJ
            istate = istates_by_id[segment.initial_state_id]
            try:
                self.used_initial_states[istate.state_id] = self.avail_initial_states.pop(istate.state_id)
            except KeyError:
                # Shared by more than one segment, and already marked as used
                pass
            
        for used_istate in self.used_initial_states.values():
            used_istate.iter_used = 1
                    
    def rebin_current(self, parent_segments):
        '''Reconstruct walkers for the current iteration based on (presumably) new binning.
        The previous iteration's segments must be provided (as ``parent_segments``) in order
        to update endpoint types appropriately.'''

        self._prep_we()
        self._parent_map = {segment.seg_id: segment for segment in parent_segments}
        
        # Create new segments for the next iteration        
        # We assume that everything is going to continue without being touched by recycling or WE, and
        # adjust later
        new_pcoord_array = self.system.new_pcoord_array
        n_iter = None
                
        for ibin, _bin in enumerate(self.final_binning):
            for segment in _bin:
                if n_iter is None:
                    n_iter = segment.n_iter
                else:
                    assert segment.n_iter == n_iter
                    
                new_segment = Segment(n_iter=segment.n_iter,
                                      parent_id=segment.parent_id,
                                      weight=segment.weight,
                                      wtg_parent_ids=set(segment.wtg_parent_ids or []),
                                      pcoord=new_pcoord_array(),
                                      status=Segment.SEG_STATUS_PREPARED)
                new_segment.pcoord[0] = segment.pcoord[0]
                self.next_iter_binning[ibin].add(new_segment)
                
        self._run_we()
                                    
    def construct_next(self):
        '''Construct walkers for the next iteration, by running weighted ensemble recycling
        and bin/split/merge on the segments previously assigned to bins using ``assign``.
        Enough unused initial states must be present in ``self.avail_initial_states`` for every recycled
        walker to be assigned an initial state.
        
        After this function completes, ``self.flux_matrix`` contains a valid flux matrix for this
        iteration (including any contributions from recycling from the previous iteration), and
        ``self.next_iter_segments`` contains a list of segments ready for the next iteration,
        with appropriate values set for weight, endpoint type, parent walkers, and so on.        
        '''
        
        self._prep_we()
        
        # Create new segments for the next iteration        
        # We assume that everything is going to continue without being touched by recycling or WE, and
        # adjust later
        new_pcoord_array = self.system.new_pcoord_array
        n_iter = None
                
        for ibin, _bin in enumerate(self.final_binning):
            for segment in _bin:
                if n_iter is None:
                    n_iter = segment.n_iter
                else:
                    assert segment.n_iter == n_iter
                    
                segment.endpoint_type = Segment.SEG_ENDPOINT_CONTINUES
                new_segment = Segment(n_iter=segment.n_iter+1,
                                      parent_id=segment.seg_id,
                                      weight=segment.weight,
                                      wtg_parent_ids=[segment.seg_id],
                                      pcoord=new_pcoord_array(),
                                      status=Segment.SEG_STATUS_PREPARED)
                new_segment.pcoord[0] = segment.pcoord[-1]
                self.next_iter_binning[ibin].add(new_segment)
                
                # Store a link to the parent segment, so we can update its endpoint status as we need,
                # based on its ID
                self._parent_map[segment.seg_id] = segment
                
        self._run_we()
        
        log.debug('used initial states: {!r}'.format(self.used_initial_states))
        log.debug('available initial states: {!r}'.format(self.avail_initial_states))
    
    def _log_bin_stats(self, bin, heading=None, level=logging.DEBUG):
        if log.isEnabledFor(level):
            weights = sorted(numpy.array(list(map(operator.attrgetter('weight'), bin))))
            bin_label = getattr(bin, 'label', None) or ''
            log_fmt = '\n      '.join(['', 
                                         'stats for bin {bin_label!r} {heading}',
                                         '  count: {bin.count:d}, target count: {bin.target_count:d}',
                                         '  total weight: {bin.weight:{weight_spec}},  ideal weight: {ideal_weight:{weight_spec}}',
                                         '  mean weight:  {mean_weight:{weight_spec}},  stdev weight: {stdev_weight:{weight_spec}}',
                                         '  min weight:   {min_weight:{weight_spec}},  med weight  : {median_weight:{weight_spec}}'
                                         +', max weight: {max_weight:{weight_spec}}'])
            log_msg = log_fmt.format(log_fmt, weight_spec='<12.6e', 
                                     bin_label=bin_label, heading=heading, bin=bin,
                                     ideal_weight = bin.weight/bin.target_count,
                                     mean_weight = weights.mean(), stdev_weight = weights.std(),
                                     min_weight = weights[0], median_weight=numpy.median(weights), max_weight=weights[-1])
            log.log(level, log_msg)
                    
            
```
