## Supplementary material for "Markovian Weighted Ensemble Milestoning (M-WEM): Long-time Kinetics from Short Trajectories": MWEM-main.zip: env.sh

```
#!/bin/bash

# Set up environment for westpa
export WEST_PYTHON=$(which python)
# Actviate a conda environment containing westpa, openmm, mdtraj;
# You may need to create this first (see install instructions)
source activate westpa
export WEST_SIM_ROOT="$PWD"
export SIM_NAME=$(basename $WEST_SIM_ROOT)
```
