## Supplementary material for "Markovian Weighted Ensemble Milestoning (M-WEM): Long-time Kinetics from Short Trajectories": MWEM-main.zip: manager.py

```
import time, operator, math, numpy, random
from itertools import zip_longest
from datetime import timedelta
import logging
log = logging.getLogger(__name__)

import westpa
import west
from west.states import InitialState
from westpa import extloader
from west import Segment

from west import wm_ops
from west.data_manager import weight_dtype

from pickle import PickleError

EPS = numpy.finfo(weight_dtype).eps

def grouper(n, iterable, fillvalue=None):
    "Collect data into fixed-length chunks or blocks"
    # grouper(3, 'ABCDEFG', 'x') --> ABC DEF Gxx
    args = [iter(iterable)] * n
    return zip_longest(fillvalue=fillvalue, *args)

class PropagationError(RuntimeError):
    pass 

class WESimManager:
    def process_config(self):
        config = self.rc.config
        for (entry, type_) in [('gen_istates', bool),
                               ('block_size', int),
                               ('save_transition_matrices', bool)]:
            config.require_type_if_present(['west', 'propagation', entry], type_)
            
        self.do_gen_istates = config.get(['west', 'propagation', 'gen_istates'], False) 
        self.propagator_block_size = config.get(['west', 'propagation', 'block_size'], 1)
        self.save_transition_matrices = config.get(['west', 'propagation', 'save_transition_matrices'], False)
        self.max_run_walltime = config.get(['west', 'propagation', 'max_run_wallclock'], default=None)
        self.max_total_iterations = config.get(['west', 'propagation', 'max_total_iterations'], default=None)
            
    
    def __init__(self, rc=None):        
        self.rc = rc or westpa.rc
        self.work_manager = self.rc.get_work_manager()
        self.data_manager = self.rc.get_data_manager()
        self.we_driver = self.rc.get_we_driver()
        self.system = self.rc.get_system_driver()
                                
        # A table of function -> list of (priority, name, callback) tuples
        self._callback_table = {}
        self._valid_callbacks = set((self.prepare_run, self.finalize_run,
                                     self.prepare_iteration, self.finalize_iteration,
                                     self.pre_propagation, self.post_propagation,
                                     self.pre_we, self.post_we, self.prepare_new_iteration))
        self._callbacks_by_name = {fn.__name__: fn for fn in self._valid_callbacks}
        self.n_propagated = 0
        
        # config items
        self.do_gen_istates = False
        self.propagator_block_size = 1
        self.save_transition_matrices = False
        self.max_run_walltime = None
        self.max_total_iterations = None
        self.process_config()
                
        # Per-iteration variables
        self.n_iter = None                  # current iteration
        
        # Basis and initial states for this iteration, in case the propagator needs them
        self.current_iter_bstates = None       # BasisStates valid at this iteration
        self.current_iter_istates = None       # InitialStates used in this iteration

        # Basis states for next iteration                
        self.next_iter_bstates = None          # BasisStates valid for the next iteration
        self.next_iter_bstate_cprobs = None    # Cumulative probabilities for basis states, used for selection
        
        # Initial states for next iteration
        #self.next_iter_istates = None
        #self.next_iter_avail_istates = None    # InitialStates available for use next iteration
        #self.next_iter_assigned_istates = None # InitialStates that were available or generated in this iteration but then used

        # Tracking of this iteration's segments        
        self.segments = None                # Mapping of seg_id to segment for all segments in this iteration
        self.completed_segments = None      # Mapping of seg_id to segment for all completed segments in this iteration
        self.incomplete_segments = None     # Mapping of seg_id to segment for all incomplete segments in this iteration
        
        # Tracking of binning
        self.bin_mapper_hash = None         # Hash of bin mapper from most recently-run WE, for use by post-WE analysis plugins
        
        
    def register_callback(self, hook, function, priority=0):
        '''Registers a callback to execute during the given ``hook`` into the simulation loop. The optional
        priority is used to order when the function is called relative to other registered callbacks.'''
        
        if hook not in self._valid_callbacks:
            try:
                hook = self._callbacks_by_name[hook]
            except KeyError:
                raise KeyError('invalid hook {!r}'.format(hook))
            
        try:
            self._callback_table[hook].add((priority,function.__name__,function))
        except KeyError:
            self._callback_table[hook] = set([(priority,function.__name__,function)])
        
        log.debug('registered callback {!r} for hook {!r}'.format(function, hook))
                
    def invoke_callbacks(self, hook, *args, **kwargs):
        callbacks = self._callback_table.get(hook, [])
        sorted_callbacks = sorted(callbacks)
        for (priority, name, fn) in sorted_callbacks:
            log.debug('invoking callback {!r} for hook {!r}'.format(fn,hook))
            fn(*args, **kwargs)
    
    def load_plugins(self):
        try:
            plugins_config = westpa.rc.config['west', 'plugins']
        except KeyError:
            return
        
        for plugin_config in (plugins_config or []):
            plugin_name = plugin_config['plugin']
            if plugin_config.get('enabled', True):
                log.info('loading plugin {!r}'.format(plugin_name))
                plugin = extloader.get_object(plugin_name)(self, plugin_config)
                log.debug('loaded plugin {!r}'.format(plugin))

    def report_bin_statistics(self, bins, save_summary=False):
        segments = list(self.segments.values())
        bin_counts = numpy.fromiter(map(len,bins), dtype=numpy.int_, count=len(bins))
        target_counts = self.we_driver.bin_target_counts

        # Do not include bins with target count zero (e.g. sinks, never-filled bins) in the (non)empty bins statistics
        n_active_bins = len(target_counts[target_counts!=0])
        seg_probs = numpy.fromiter(map(operator.attrgetter('weight'), segments), dtype=weight_dtype, count=len(segments))
        bin_probs = numpy.fromiter(map(operator.attrgetter('weight'), bins), dtype=weight_dtype, count=len(bins)) 
        norm = seg_probs.sum()
        
        assert abs(1 - norm) < EPS*(len(segments)+n_active_bins)
        
        min_seg_prob = seg_probs[seg_probs!=0].min()
        max_seg_prob = seg_probs.max()
        seg_drange   = math.log(max_seg_prob/min_seg_prob)
        min_bin_prob = bin_probs[bin_probs!=0].min()
        max_bin_prob = bin_probs.max()
        bin_drange = math.log(max_bin_prob/min_bin_prob)
        n_pop = int(len(segments)/target_counts[0])
        
        self.rc.pstatus('{:d} of {:d} ({:%}) active bins are populated'.format(n_pop, n_active_bins,n_pop/n_active_bins))
        self.rc.pstatus('per-bin minimum non-zero probability:       {:g}'.format(min_bin_prob))
        self.rc.pstatus('per-bin maximum probability:                {:g}'.format(max_bin_prob))
        self.rc.pstatus('per-bin probability dynamic range (kT):     {:g}'.format(bin_drange))
        self.rc.pstatus('per-segment minimum non-zero probability:   {:g}'.format(min_seg_prob))
        self.rc.pstatus('per-segment maximum non-zero probability:   {:g}'.format(max_seg_prob))
        self.rc.pstatus('per-segment probability dynamic range (kT): {:g}'.format(seg_drange))
        self.rc.pstatus('norm = {:g}, error in norm = {:g} ({:.2g}*epsilon)'.format(norm,(norm-1),(norm-1)/EPS))
        self.rc.pflush()
        
        if save_summary:
            iter_summary = self.data_manager.get_iter_summary()
            iter_summary['n_particles'] = len(segments)
            iter_summary['norm'] = norm
            iter_summary['min_bin_prob'] = min_bin_prob
            iter_summary['max_bin_prob'] = max_bin_prob
            iter_summary['min_seg_prob'] = min_seg_prob
            iter_summary['max_seg_prob'] = max_seg_prob
            if numpy.isnan(iter_summary['cputime']): iter_summary['cputime'] = 0.0
            if numpy.isnan(iter_summary['walltime']): iter_summary['walltime'] = 0.0
            self.data_manager.update_iter_summary(iter_summary)

    def get_bstate_pcoords(self, basis_states):
        '''For each of the given ``basis_states``, calculate progress coordinate values
        as necessary.  The HDF5 file is not updated.'''
        
        self.rc.pstatus('Calculating progress coordinate values for basis states.')
        futures = [self.work_manager.submit(wm_ops.get_pcoord, args=(basis_state,))
                   for basis_state in basis_states]
        fmap = {future: i for (i, future) in enumerate(futures)}
        for future in self.work_manager.as_completed(futures): 
            basis_states[fmap[future]].pcoord = future.get_result().pcoord
        
    def report_basis_states(self, basis_states):
        pstatus = self.rc.pstatus
        pstatus('{:d} basis state(s) present'.format(len(basis_states)), end='')
        if self.rc.verbose_mode:
            pstatus(':')
            pstatus('{:6s}    {:12s}    {:20s}    {:20s}    {}'
                             .format('ID', 'Label', 'Probability', 'Aux Reference', 'Progress Coordinate'))
            for basis_state in basis_states:
                pstatus('{:<6d}    {:12s}    {:<20.14g}    {:20s}    {}'.
                                 format(basis_state.state_id, basis_state.label, basis_state.probability, basis_state.auxref or '',
                                        ', '.join(map(str,basis_state.pcoord))))
        pstatus()
        self.rc.pflush()
        
    def report_target_states(self, target_states):
        pstatus = self.rc.pstatus
        pstatus('{:d} target state(s) present'.format(len(target_states)), end='')    
        if self.rc.verbose_mode and target_states:
            pstatus(':')
            pstatus('{:6s}    {:12s}    {}'.format('ID', 'Label', 'Progress Coordinate'))
            for target_state in target_states:
                pstatus('{:<6d}    {:12s}    {}'
                                 .format(target_state.state_id, target_state.label, ','.join(map(str,target_state.pcoord))))        
        pstatus()
        self.rc.pflush()
        
    def initialize_simulation(self, basis_states, target_states, segs_per_state=1, suppress_we=False):
        '''Initialize a new weighted ensemble simulation, taking ``segs_per_state`` initial
        states from each of the given ``basis_states``.
        
        ``w_init`` is the forward-facing version of this function'''
        
        data_manager = self.data_manager
        work_manager = self.work_manager
        pstatus = self.rc.pstatus
        system = self.system

        pstatus('Creating HDF5 file {!r}'.format(self.data_manager.we_h5filename))
        data_manager.prepare_backing()

        # Process target states
        data_manager.save_target_states(target_states)
        self.report_target_states(target_states)

        
        # Process basis states
        self.get_bstate_pcoords(basis_states)        
        self.data_manager.create_ibstate_group(basis_states)
        self.report_basis_states(basis_states)
        
        pstatus('Preparing initial states')
        initial_states = []
        weights = []
        if self.do_gen_istates:
            istate_type = InitialState.ISTATE_TYPE_GENERATED
        else:
            istate_type = InitialState.ISTATE_TYPE_BASIS
            
        for basis_state in basis_states:
            for _iseg in range(segs_per_state):
                initial_state = data_manager.create_initial_states(1,1)[0]
                initial_state.basis_state_id =  basis_state.state_id
                initial_state.basis_state = basis_state
                initial_state.istate_type = istate_type
                weights.append(basis_state.probability/segs_per_state)
                initial_states.append(initial_state)
                
        if self.do_gen_istates:
            futures = [work_manager.submit(wm_ops.gen_istate, args=(initial_state.basis_state, initial_state))
                       for initial_state in initial_states]
            for future in work_manager.as_completed(futures):
                rbstate, ristate = future.get_result()
                initial_states[ristate.state_id].pcoord = ristate.pcoord
        else:
            for initial_state in initial_states:
                basis_state = initial_state.basis_state
                initial_state.pcoord = basis_state.pcoord
                initial_state.istate_status = InitialState.ISTATE_STATUS_PREPARED
                
        for initial_state in initial_states:
            log.debug('initial state created: {!r}'.format(initial_state))            


        # save list of initial states just generated
        # some of these may not be used, depending on how WE shakes out                    
        data_manager.update_initial_states(initial_states, n_iter=1)
                        
        if not suppress_we:
            self.we_driver.populate_initial(initial_states, weights, system)
            segments = list(self.we_driver.next_iter_segments)
            binning = self.we_driver.next_iter_binning
        else:
            segments = list(self.we_driver.current_iter_segments)
            binning = self.we_driver.final_binning

        bin_occupancies = numpy.fromiter(map(len,binning), dtype=numpy.uint, count=self.we_driver.bin_mapper.nbins)
        target_occupancies = numpy.require(self.we_driver.bin_target_counts, dtype=numpy.uint)
    
        # Make sure we have 
        for segment in segments:
            segment.n_iter = 1
            segment.status = Segment.SEG_STATUS_PREPARED
            assert segment.parent_id < 0
            assert initial_states[segment.initial_state_id].iter_used == 1        
                    
        data_manager.prepare_iteration(1, segments)
        data_manager.update_initial_states(initial_states, n_iter=1)
                    
        if self.rc.verbose_mode:
            pstatus('\nSegments generated:')
            for segment in segments:
                pstatus('{!r}'.format(segment))
        
        
        pstatus('''
        Total bins:            {total_bins:d}
        Initial replicas:      {init_replicas:d} in {occ_bins:d} bins, total weight = {weight:g}
        Total target replicas: {total_replicas:d}
        '''.format(total_bins=len(bin_occupancies),
                   init_replicas=int(sum(bin_occupancies)),
                   occ_bins=len(bin_occupancies[bin_occupancies > 0]),
                   weight = float(sum(segment.weight for segment in segments)),
                   total_replicas = int(sum(target_occupancies))))
        
        # Send the segments over to the data manager to commit to disk            
        data_manager.current_iteration = 1
        
        # Report statistics
        pstatus('Simulation prepared.')
        self.segments = {segment.seg_id: segment for segment in segments}
        self.report_bin_statistics(binning,save_summary=True)
        data_manager.flush_backing()

    def prepare_iteration(self):
        log.debug('beginning iteration {:d}'.format(self.n_iter))
                
        # the WE driver needs a list of all target states for this iteration
        # along with information about any new weights introduced (e.g. by recycling)
        target_states = self.data_manager.get_target_states(self.n_iter)
        new_weights = self.data_manager.get_new_weight_data(self.n_iter)
        
        self.we_driver.new_iteration(target_states=target_states, new_weights= new_weights)
                
        # Get basis states used in this iteration
        self.current_iter_bstates = self.data_manager.get_basis_states(self.n_iter)
        
        # Get the segments for this iteration and separate into complete and incomplete
        if self.segments is None:
            segments = self.segments = {segment.seg_id: segment for segment in self.data_manager.get_segments()}
            log.debug('loaded {:d} segments'.format(len(segments)))
        else:
            segments = self.segments
            log.debug('using {:d} pre-existing segments'.format(len(segments)))
        
        completed_segments = self.completed_segments = {}
        incomplete_segments = self.incomplete_segments = {}
        for segment in segments.values():
            if segment.status == Segment.SEG_STATUS_COMPLETE:
                completed_segments[segment.seg_id] = segment
            else:
                incomplete_segments[segment.seg_id] = segment
        log.debug('{:d} segments are complete; {:d} are incomplete'.format(len(completed_segments), len(incomplete_segments)))
        
        if len(incomplete_segments) == len(segments):
            # Starting a new iteration
            self.rc.pstatus('Beginning iteration {:d}'.format(self.n_iter))
        elif incomplete_segments:
            self.rc.pstatus('Continuing iteration {:d}'.format(self.n_iter))
        self.rc.pstatus('{:d} segments remain in iteration {:d} ({:d} total)'.format(len(incomplete_segments), self.n_iter,
                                                                                     len(segments)))
        
        # Get the initial states active for this iteration (so that the propagator has them if necessary)
        self.current_iter_istates = {state.state_id: state for state in 
                                     self.data_manager.get_segment_initial_states(list(segments.values()))}
        log.debug('This iteration uses {:d} initial states'.format(len(self.current_iter_istates)))
        
        # Assign this iteration's segments' initial points to bins and report on bin population
        initial_pcoords = self.system.new_pcoord_array(len(segments))
        initial_binning = self.system.bin_mapper.construct_bins()
        for iseg, segment in enumerate(segments.values()):
            initial_pcoords[iseg] = segment.pcoord[0]
        initial_assignments = self.system.bin_mapper.assign(initial_pcoords)
        for (segment, assignment) in zip(iter(segments.values()), initial_assignments):
            initial_binning[assignment].add(segment)
        self.report_bin_statistics(initial_binning, save_summary=True)
        del initial_pcoords, initial_binning
        
        # Let the WE driver assign completed segments 
        if completed_segments:
            self.we_driver.assign(list(completed_segments.values()))
        
        # Get the basis states and initial states for the next iteration, necessary for doing on-the-fly recycling 
        self.next_iter_bstates = self.data_manager.get_basis_states(self.n_iter+1)
        self.next_iter_bstate_cprobs = numpy.add.accumulate([bstate.probability for bstate in self.next_iter_bstates])
        
        self.we_driver.avail_initial_states = {istate.state_id: istate 
                                               for istate in self.data_manager.get_unused_initial_states(n_iter=self.n_iter+1)}
        log.debug('{:d} unused initial states found'.format(len(self.we_driver.avail_initial_states)))
        
        # Invoke callbacks
        self.invoke_callbacks(self.prepare_iteration)
        
        # dispatch and immediately wait on result for prep_iter
        log.debug('dispatching propagator prep_iter to work manager')        
        self.work_manager.submit(wm_ops.prep_iter, args=(self.n_iter, segments)).get_result()
                
    def finalize_iteration(self):
        '''Clean up after an iteration and prepare for the next.'''
        log.debug('finalizing iteration {:d}'.format(self.n_iter))
        
        self.invoke_callbacks(self.finalize_iteration)
        
        # dispatch and immediately wait on result for post_iter
        log.debug('dispatching propagator post_iter to work manager')
        self.work_manager.submit(wm_ops.post_iter, args=(self.n_iter, list(self.segments.values()))).get_result()
        
        # Move existing segments into place as new segments
        del self.segments
        self.segments = {segment.seg_id: segment for segment in self.we_driver.next_iter_segments}
                        
    def get_istate_futures(self):
        '''Add ``n_states`` initial states to the internal list of initial states assigned to
        recycled particles.  Spare states are used if available, otherwise new states are created.
        If created new initial states requires generation, then a set of futures is returned
        representing work manager tasks corresponding to the necessary generation work.'''
        
        n_recycled = self.we_driver.n_recycled_segs
        n_istates_needed = self.we_driver.n_istates_needed

        log.debug('{:d} unused initial states available'.format(len(self.we_driver.avail_initial_states)))                
        log.debug('{:d} new initial states required for recycling {:d} walkers'.format(n_istates_needed,
                                                                                       n_recycled))
        
        futures = set()
        updated_states = []
        for _i in range(n_istates_needed):
            # Select a basis state according to its weight
            ibstate = numpy.digitize([random.random()], self.next_iter_bstate_cprobs)
            basis_state = self.next_iter_bstates[ibstate[0]]
            initial_state = self.data_manager.create_initial_states(1, n_iter=self.n_iter+1)[0]
            initial_state.iter_created = self.n_iter
            initial_state.basis_state_id = basis_state.state_id
            initial_state.istate_status = InitialState.ISTATE_STATUS_PENDING
            
            if self.do_gen_istates:
                log.debug('generating new initial state from basis state {!r}'.format(basis_state))
                initial_state.istate_type = InitialState.ISTATE_TYPE_GENERATED
                futures.add(self.work_manager.submit(wm_ops.gen_istate, args=(basis_state, initial_state)))
            else:
                log.debug('using basis state {!r} directly'.format(basis_state))
                initial_state.istate_type = InitialState.ISTATE_TYPE_BASIS
                initial_state.pcoord = basis_state.pcoord.copy()
                initial_state.istate_status = InitialState.ISTATE_STATUS_PREPARED
                self.we_driver.avail_initial_states[initial_state.state_id] = initial_state
            updated_states.append(initial_state)
        self.data_manager.update_initial_states(updated_states, n_iter=self.n_iter+1)
        return futures
                                    
    def propagate(self):
        segments = list(self.incomplete_segments.values())
        log.debug('iteration {:d}: propagating {:d} segments'.format(self.n_iter, len(segments)))
        
        # all futures dispatched for this iteration
        futures = set()        
        segment_futures = set()
        
        # Immediately dispatch any necessary initial state generation
        istate_gen_futures = self.get_istate_futures()
        futures.update(istate_gen_futures)
        
        # Dispatch propagation tasks using work manager                
        for segment_block in grouper(self.propagator_block_size, segments):
            segment_block = [_f for _f in segment_block if _f]
            pbstates, pistates = west.states.pare_basis_initial_states(self.current_iter_bstates, 
                                                                       list(self.current_iter_istates.values()), segment_block)
            future = self.work_manager.submit(wm_ops.propagate, args=(pbstates, pistates, segment_block))
            futures.add(future)
            segment_futures.add(future)
        
        while futures:
            # TODO: add capacity for timeout or SIGINT here
            future = self.work_manager.wait_any(futures)
            futures.remove(future)
            
            if future in segment_futures:
                segment_futures.remove(future)
                incoming = future.get_result()
                self.n_propagated += 1
                
                self.segments.update({segment.seg_id: segment for segment in incoming})
                self.completed_segments.update({segment.seg_id: segment for segment in incoming})
                
               # self.we_driver.assign(incoming)
                new_istate_futures = self.get_istate_futures()
                istate_gen_futures.update(new_istate_futures)
                futures.update(new_istate_futures)
                
                with self.data_manager.expiring_flushing_lock():                        
                    self.data_manager.update_segments(self.n_iter, incoming)

            elif future in istate_gen_futures:
                istate_gen_futures.remove(future)
                _basis_state, initial_state = future.get_result()
                log.debug('received newly-prepared initial state {!r}'.format(initial_state))
                initial_state.istate_status = InitialState.ISTATE_STATUS_PREPARED
                with self.data_manager.expiring_flushing_lock():
                    self.data_manager.update_initial_states([initial_state], n_iter=self.n_iter+1)
                self.we_driver.avail_initial_states[initial_state.state_id] = initial_state
            else:
                log.error('unknown future {!r} received from work manager'.format(future))
                raise AssertionError('untracked future {!r}'.format(future))                    
        #WARNING
        self.we_driver.assign(self.segments.values(), endprop=True)
        self.get_istate_futures()            
        log.debug('done with propagation')
        self.save_bin_data()
        self.data_manager.flush_backing()
        
    def save_bin_data(self):
        '''Calculate and write flux and transition count matrices to HDF5. Population and rate matrices 
        are likely useless at the single-tau level and are no longer written.'''
        # save_bin_data(self, populations, n_trans, fluxes, rates, n_iter=None)
        
        if self.save_transition_matrices:
            with self.data_manager.expiring_flushing_lock():
                iter_group = self.data_manager.get_iter_group(self.n_iter)
                for key in ['bin_ntrans', 'bin_fluxes']:
                    try:
                        del iter_group[key]
                    except KeyError:
                        pass
                iter_group['bin_ntrans'] = self.we_driver.transition_matrix
                iter_group['bin_fluxes'] = self.we_driver.flux_matrix
        
    def check_propagation(self):
        '''Check for failures in propagation or initial state generation, and raise an exception
        if any are found.'''
        
        failed_segments = [segment for segment in self.segments.values() if segment.status != Segment.SEG_STATUS_COMPLETE]
        
        if failed_segments:
            failed_ids = '  \n'.join(str(segment.seg_id) for segment in failed_segments)
            log.error('propagation failed for {:d} segment(s):\n{}'.format(len(failed_segments), failed_ids))
            raise PropagationError('propagation failed for {:d} segments'.format(len(failed_segments)))
        else:
            log.debug('propagation complete for iteration {:d}'.format(self.n_iter))
            
        failed_istates = [istate for istate in self.we_driver.used_initial_states.values()
                          if istate.istate_status != InitialState.ISTATE_STATUS_PREPARED]
        log.debug('{!r}'.format(failed_istates))
        if failed_istates:
            failed_ids = '  \n'.join(str(istate.state_id) for istate in failed_istates)
            log.error('initial state generation failed for {:d} states:\n{}'.format(len(failed_istates), failed_ids))
            raise PropagationError('initial state generation failed for {:d} states'.format(len(failed_istates)))
        else:
            log.debug('initial state generation complete for iteration {:d}'.format(self.n_iter))


    def run_we(self):
        '''Run the weighted ensemble algorithm based on the binning in self.final_bins and
        the recycled particles in self.to_recycle, creating and committing the next iteration's
        segments to storage as well.'''
        
        # The WE driver now does almost everything; we just have to record the
        # mapper used for binning this iteration, and update initial states
        # that have been used
         
        try:
            pickled, hashed = self.we_driver.bin_mapper.pickle_and_hash()
        except PickleError:
            pickled = hashed = ''

        self.bin_mapper_hash = hashed
        self.we_driver.construct_next()
        
        if self.we_driver.used_initial_states:
            for initial_state in self.we_driver.used_initial_states.values():
                initial_state.iter_used = self.n_iter+1
            self.data_manager.update_initial_states(list(self.we_driver.used_initial_states.values()))
            
        self.data_manager.update_segments(self.n_iter,list(self.segments.values()))
        
        self.data_manager.require_iter_group(self.n_iter+1)
        self.data_manager.save_iter_binning(self.n_iter+1, hashed, pickled, self.we_driver.bin_target_counts)
        
        # Report on recycling
        recycling_events = {}
        for nw in self.we_driver.new_weights:
            try:
                recycling_events[nw.target_state_id].add(nw.weight)
            except KeyError:
                recycling_events[nw.target_state_id] = set([nw.weight])
        
        tstates_by_id = {state.state_id: state for state in self.we_driver.target_states.values()}
        for tstate_id, weights in recycling_events.items():
            tstate = tstates_by_id[tstate_id]
            self.rc.pstatus('Recycled {:g} probability ({:d} walkers) from target state {!r}'.format(sum(weights),
                                                                                                     len(weights),
                                                                                                     tstate.label))
                
    def prepare_new_iteration(self):
        '''Commit data for the coming iteration to the HDF5 file.'''
        self.invoke_callbacks(self.prepare_new_iteration)

        if self.rc.debug_mode:
            self.rc.pstatus('\nSegments generated:')
            for segment in self.we_driver.next_iter_segments:
                self.rc.pstatus('{!r} pcoord[0]={!r}'.format(segment, segment.pcoord[0]))
        
        self.data_manager.prepare_iteration(self.n_iter+1, list(self.we_driver.next_iter_segments))
        self.data_manager.save_new_weight_data(self.n_iter+1, self.we_driver.new_weights)
        
    def run(self):   
        run_starttime = time.time()
        max_walltime = self.max_run_walltime
        if max_walltime:
            run_killtime = run_starttime + max_walltime
            self.rc.pstatus('Maximum wallclock time: %s' % timedelta(seconds=max_walltime or 0))
        else:
            run_killtime = None
        
        self.n_iter = self.data_manager.current_iteration    
        max_iter = self.max_total_iterations or self.n_iter+1

        iter_elapsed = 0
        while self.n_iter <= max_iter:
            
            if max_walltime and time.time() + 1.1*iter_elapsed >= run_killtime:
                self.rc.pstatus('Iteration {:d} would require more than the allotted time. Ending run.'
                                .format(self.n_iter))
                return
            
            try:
                iter_start_time = time.time()
                
                self.rc.pstatus('\n%s' % time.asctime())
                self.rc.pstatus('Iteration %d (%d requested)' % (self.n_iter, max_iter))
                                
                self.prepare_iteration()
                self.rc.pflush()
                
                self.pre_propagation()
                self.propagate()
                self.rc.pflush()
                self.check_propagation()
                self.rc.pflush()
                self.post_propagation()
                
                cputime = sum(segment.cputime for segment in self.segments.values())
                
                self.rc.pflush()
                self.pre_we()
                self.run_we()
                self.post_we()
                self.rc.pflush()
                
                self.prepare_new_iteration()
                
                self.finalize_iteration()
                
                iter_elapsed = time.time() - iter_start_time
                iter_summary = self.data_manager.get_iter_summary()
                iter_summary['walltime'] += iter_elapsed
                iter_summary['cputime'] = cputime
                self.data_manager.update_iter_summary(iter_summary)
    
                self.n_iter += 1
                self.data_manager.current_iteration += 1

                try:
                    #This may give NaN if starting a truncated simulation
                    walltime = timedelta(seconds=float(iter_summary['walltime']))
                except ValueError:
                    walltime = 0.0 
                
                try:
                    cputime = timedelta(seconds=float(iter_summary['cputime']))
                except ValueError:
                    cputime = 0.0      

                self.rc.pstatus('Iteration wallclock: {0!s}, cputime: {1!s}\n'\
                                          .format(walltime,
                                                  cputime))
                self.rc.pflush()
            finally:
                self.data_manager.flush_backing()
                
        self.rc.pstatus('\n%s' % time.asctime())
        self.rc.pstatus('WEST run complete.')
        
    def prepare_run(self):
        '''Prepare a new run.'''
        self.data_manager.prepare_run()
        self.system.prepare_run()
        self.invoke_callbacks(self.prepare_run)
    
    def finalize_run(self):
        '''Perform cleanup at the normal end of a run'''
        self.invoke_callbacks(self.finalize_run)
        self.system.finalize_run()
        self.data_manager.finalize_run()
        
    def pre_propagation(self):
        self.invoke_callbacks(self.pre_propagation)
        
    def post_propagation(self):
        self.invoke_callbacks(self.post_propagation)
        
    def pre_we(self):
        self.invoke_callbacks(self.pre_we)
    
    def post_we(self):
        self.invoke_callbacks(self.post_we)
```
