## Supplementary material for "Markovian Weighted Ensemble Milestoning (M-WEM): Long-time Kinetics from Short Trajectories": MWEM-main.zip: README.md

# Markovian Weighted Ensemble Milestoning (MWEM) 
This directory performs the weighted ensemble simulation for one cell. This directory is copied and modified for each milestone by the code build.py

## Authors

* **Dhiman Ray** - *Primary work* - [dhimanray](https://github.com/dhimanray)
The original readme file from WESTPA tutorial is copied below:
-----------------------------------------------------------------------------------------------------------

# Basic WESTPA tutorial: Na+ Cl- association
This tutorial is meant to introduce new users to the WESTPA software for running Weighted Ensemble (WE) simulations.

## Tutorial files

All files necessary for completing the tutorial can be found in this directory.

## Full tutorial 

Find the full tutorial in the LiveComs journal article on best practices for Weighted Ensemble simulations. (Link to be added)

## Authors

* **Anthony Bogetti** - *Primary work* - [atbogetti](https://github.com/atbogetti)
* **Alex DeGrave** - *Preliminary work* - [ajd98](https://github.com/ajd98)
