## Supplementary material for "Markovian Weighted Ensemble Milestoning (M-WEM): Long-time Kinetics from Short Trajectories": MWEM-main.zip: run.slurm

```
#!/bin/bash

#SBATCH --job-name=milestoning      ## job name
#SBATCH --account andricio_lab     ## account to charge
#SBATCH -p standard          ## partition/queue name
#SBATCH --nodes=1            ## (-N) number of nodes to use
#SBATCH --cpus-per-task=4    ## number of cores the job needs
#SBATCH -t 72:00:00
#SBATCH --mem=8gb
#SBATCH --error=slurm-%J.err ## error log file


source /data/homezvol2/dray1/Miniconda2/etc/profile.d/conda.sh

#equilibrate
cd equilibration
/data/homezvol2/dray1/NAMD_2.13_Linux-x86_64-multicore/namd2 +p 4 equilibration.conf > equilibration.log
#*** Make sure to provide the correct path to NAMD in your cluster in above line ***#

python calc_rxn_coor.py > distance.dat
cd ..

# Make sure environment is set
source env.sh


# Clean up from previous/ failed runs
rm -rf traj_segs seg_logs istates west.h5 binbounds.txt
mkdir   seg_logs traj_segs istates

#copy files to bstate directory for starting points
cp equilibration/distance.dat bstates/dist.dat
cp equilibration/milestone_equilibration.restart.coor bstates/seg.coor
cp equilibration/milestone_equilibration.colvars.traj  bstates/seg.colvars.traj
cp equilibration/milestone_equilibration.restart.vel  bstates/seg.vel
cp equilibration/milestone_equilibration.restart.xsc  bstates/seg.xsc
cp common_files/colvars.in bstates/colvars.in

# Set pointer to bstate and tstate
BSTATE_ARGS="--bstate-file $WEST_SIM_ROOT/bstates/bstates.txt"
#TSTATE_ARGS="--tstate-file $WEST_SIM_ROOT/tstate.file"

# Run w_init
$WEST_ROOT/bin/w_init $BSTATE_ARGS  $TSTATE_ARGS  --segs-per-state 4  --work-manager=threads "$@"


# Clean up
rm -f west.log 


# Run unrestrained dynamics
$WEST_ROOT/bin/w_run -r west.cfg --work-manager processes --n-workers 4 "$@" &> west.log
```
