## Supplementary material for "Markovian Weighted Ensemble Milestoning (M-WEM): Long-time Kinetics from Short Trajectories": MWEM-main.zip: west.cfg

```
# The master WEST configuration file for a simulation.
# vi: set filetype=yaml :
---
west: 
  system:
    driver: adaptive.System
    module_path: $WEST_SIM_ROOT
  drivers:
    sim_manager: manager.WESimManager
    module_path: $WEST_SIM_ROOT
    we_driver: driver.WEDriver
  propagation:
    max_total_iterations: 100
    max_run_wallclock:    94:00:00
    propagator:           executable
    gen_istates:          false
  data:
    west_data_file: west.h5
    datasets:
      - name:        pcoord
        scaleoffset: 4
      - name:        coord
        dtype:       float32
        scaleoffset: 3
    data_refs:
      segment:       $WEST_SIM_ROOT/traj_segs/{segment.n_iter:06d}/{segment.seg_id:06d}
      basis_state:   $WEST_SIM_ROOT/bstates/{basis_state.auxref}
      initial_state: $WEST_SIM_ROOT/istates/{initial_state.iter_created}/{initial_state.state_id}
  plugins:
  executable:
    environ:
      PROPAGATION_DEBUG: 1
    datasets:
      - name:    coords
        enabled: false
    propagator:
      executable: $WEST_SIM_ROOT/westpa_scripts/runseg.sh
      stdout:     $WEST_SIM_ROOT/seg_logs/{segment.n_iter:06d}-{segment.seg_id:06d}.log
      stderr:     stdout
      stdin:      null
      cwd:        null
      environ:
        SEG_DEBUG: 1
    get_pcoord:
      executable: $WEST_SIM_ROOT/westpa_scripts/get_pcoord.sh
      stdout:     /dev/null
      stderr:     stdout
    gen_istate:
      executable: $WEST_SIM_ROOT/westpa_scripts/gen_istate.sh
      stdout:     /dev/null
      stderr:     stdout
    post_iteration:
      enabled:    true
      executable: $WEST_SIM_ROOT/westpa_scripts/post_iter.sh
      stdout:     post_iter.log
      stderr:     stdout
    pre_iteration:
      enabled:    false
      executable: $WEST_SIM_ROOT/westpa_scripts/pre_iter.sh
      stderr:     stdout
#  analysis:
#       directory: ANALYSIS                
#       kinetics:                          
#         step_iter: 1
#         evolution: cumulative
#         extra: [ 'disable-correl' ]
#       analysis_schemes:                  
#         TEST:
#           enabled: True
#           bins:
#             - type: RectilinearBinMapper
#               boundaries: 
#                 - [0.0,2.4,20.8,'inf']
#           states:
#             - label: unbound
#               coords: 
#                 - [20.8] 
#             - label: bound
#               coords: 
#                 - [0]
```
