## Supplementary material for "Markovian Weighted Ensemble Milestoning (M-WEM): Long-time Kinetics from Short Trajectories": MWEM-main.zip: post_iter.sh

```
#!/bin/bash

if [ -n "$SEG_DEBUG" ] ; then
    set -x
    env | sort
fi

cd $WEST_SIM_ROOT || exit 1

ITER=$(printf "%06d" $WEST_CURRENT_ITER)
tar -cf seg_logs/$ITER.tar seg_logs/$ITER-*.log
rm  -f  seg_logs/$ITER-*.log
```
