## Supplementary material for "Markovian Weighted Ensemble Milestoning (M-WEM): Long-time Kinetics from Short Trajectories": MWEM-main.zip: runseg.sh

```
#!/bin/bash
#
# runseg.sh
#
# WESTPA runs this script for each trajectory segment. WESTPA supplies
# environment variables that are unique to each segment, such as:
#
#   WEST_CURRENT_SEG_DATA_REF: A path to where the current trajectory segment's
#       data will be stored. This will become "WEST_PARENT_DATA_REF" for any
#       child segments that spawn from this segment
#   WEST_PARENT_DATA_REF: A path to a file or directory containing data for the
#       parent segment.
#   WEST_CURRENT_SEG_INITPOINT_TYPE: Specifies whether this segment is starting
#       anew, or if this segment continues from where another segment left off.
#   WEST_RAND16: A random integer
#
# This script has the following three jobs:
#  1. Create a directory for the current trajectory segment, and set up the
#     directory for running NAMD 
#  2. Run the dynamics
#  3. Calculate the progress coordinates and return data to WESTPA


if [ -n "$SEG_DEBUG" ] ; then
  set -x
  env | sort
fi

cd $WEST_SIM_ROOT
mkdir -pv $WEST_CURRENT_SEG_DATA_REF
cd $WEST_CURRENT_SEG_DATA_REF

#ln -sv $WEST_SIM_ROOT/common_files/bstate.pdb .
#ln -sv $WEST_SIM_ROOT/common_files/DistanceReporter.py .
#ln -sv $WEST_PARENT_DATA_REF/dist.dat ./parent.dat

# Make symbolic links to the topology file and parameter files. These are not
# unique to each segment.
ln -sv $WEST_SIM_ROOT/common_files/*.psf .
ln -sv $WEST_SIM_ROOT/common_files/*.pdb .
ln -sv $WEST_SIM_ROOT/common_files/*.prm .
ln -sv $WEST_SIM_ROOT/bstates/colvars.in colvars.in   #make sure the colvars.in is from bstates and not common_files
#ln -sv $WEST_SIM_ROOT/common_files/toppar_water_ions_namd.str toppar_water_ions_namd.str

# Either continue an existing tractory, or start a new trajectory. Here, both
# cases are the same.  If you need to handle the cases separately, you can
# check the value of the environment variable "WEST_CURRENT_SEG_INIT_POINT",
# which is equal to either "SEG_INITPOINT_CONTINUES" or "SEG_INITPOINT_NEWTRAJ"
# for continuations of previous segments and new trajectories, respecitvely.
# For an example, see the nacl_amb tutorial.

# The weighted ensemble algorithm requires that dynamics are stochastic.
# We'll use the "sed" command to replace the string "RAND" with a randomly
# generated seed.
sed "s/RAND/$WEST_RAND16/g" \
  $WEST_SIM_ROOT/common_files/md.conf > md.conf

# This trajectory segment will start off where its parent segment left off.
# The "ln" command makes symbolic links to the parent segment's edr, gro, and
# and trr files. This is preferable to copying the files, since it doesn't
# require writing all the data again.
if [ "$WEST_CURRENT_SEG_INITPOINT_TYPE" = "SEG_INITPOINT_CONTINUES" ]; then
  ln -sv $WEST_PARENT_DATA_REF/seg.coor ./parent.coor
  ln -sv $WEST_PARENT_DATA_REF/seg.vel  ./parent.vel
  ln -sv $WEST_PARENT_DATA_REF/seg.xsc  ./parent.xsc
  ln -sv $WEST_PARENT_DATA_REF/dist.dat  ./parent.dat

elif [ "$WEST_CURRENT_SEG_INITPOINT_TYPE" = "SEG_INITPOINT_NEWTRAJ" ]; then
  ln -sv $WEST_SIM_ROOT/bstates/seg.coor ./parent.coor
  ln -sv $WEST_SIM_ROOT/bstates/seg.vel  ./parent.vel
  ln -sv $WEST_SIM_ROOT/bstates/seg.xsc  ./parent.xsc
  ln -sv $WEST_SIM_ROOT/bstates/dist.dat  ./parent.dat
fi
############################## Run the dynamics ################################
# Propagate the segment using namd2 
/data/homezvol2/dray1/NAMD_2.13_Linux-x86_64-multicore/namd2 md.conf > seg.log   
#*** Make sure to provide the correct path to NAMD in your cluster in above line ***#

########################## Calculate and return data ###########################

# Calculate the progress coordinate, which is the distance between the Na+ and
# Cl- ions.  This custom Python script looks for the files nacl.psf and seg.dcd
# The script outputs the distance between the ions at each timepoint, printing
# one distance value per line to STDOUT.
# Note that the script also loads the parent segment's .dcd file, in order to 
# include the current segment's starting configuration (which is the same as the
# parent segment's final configuration) in the calculation.

echo "$(pwd)"
#Calculate pcoord with MDAnalysis
#python $WEST_SIM_ROOT/common_files/get_distance.py
python $WEST_SIM_ROOT/westpa_scripts/calc_rxn_coor.py > dist.dat
cat dist.dat > $WEST_PCOORD_RETURN

# Clean up
mkdir keep
mv seg.coor keep/
mv seg.xsc keep/
mv seg.vel keep/
mv seg.colvars.traj keep/
mv seg.dcd keep/
mv dist.dat keep/
rm *
cp keep/* .
rm -r keep
```
