## Supplementary material for "Markovian Weighted Ensemble Milestoning (M-WEM): Long-time Kinetics from Short Trajectories": MWEM-main.zip: tar_segs.sh

```
#!/bin/bash

[ -z "$WEST_SIM_ROOT" ] &&
    exit 1
[ ! -d $WEST_SIM_ROOT/traj_segs ] &&
    exit 1

cd $WEST_SIM_ROOT/traj_segs

ITERS=($(ls | grep '^[0-9][0-9][0-9][0-9][0-9][0-9]$'))
ITERS=("${ITERS[@]:0:${#ITERS[@]}-1}")
for ITER in ${ITERS[@]}; do
    [ ! -f $ITER.tar ] &&
        tar -cvf $ITER.tar $ITER
done
```
