## Supplementary material for "Markovian Weighted Ensemble Milestoning (M-WEM): Long-time Kinetics from Short Trajectories": MWEM-main.zip: README.md

Scripts and templates for building the simulation setup. It creates 10 cells named cell_0, cell_1,...,cell_9
 (For Trypsin Benzamidine two additional milestones at RC=2.5 and RC=3.5 were added later manually. So they are not included in the script. Those new cells were named cell_2.5 and cell_3.5)
