## Supplementary material for "Markovian Weighted Ensemble Milestoning (M-WEM): Long-time Kinetics from Short Trajectories": MWEM-main.zip: restart.slurm

```
#!/bin/bash

#SBATCH --job-name=milestoning      ## job name
#SBATCH --account andricio_lab     ## account to charge
#SBATCH -p standard          ## partition/queue name
#SBATCH --nodes=1            ## (-N) number of nodes to use
#SBATCH --cpus-per-task=4    ## number of cores the job needs
#SBATCH --gpus-per-task=1
#SBATCH -t 72:00:00
#SBATCH --mem=8gb
#SBATCH --error=slurm-%J.err ## error log file

module load cuda/10.1.243
module load namd/2.14b2/gcc.8.4.0-cuda.10.1.243


source /data/homezvol2/dray1/Miniconda2/etc/profile.d/conda.sh


# Make sure environment is set
source env.sh


# Clean up
mv west.log west.log.old 
rm binbounds.txt

# Run unrestrained dynamics
$WEST_ROOT/bin/w_run -r west.cfg --work-manager processes --n-workers 1 "$@" &> west.log
```
