## Supplementary material for "Markovian Weighted Ensemble Milestoning (M-WEM): Long-time Kinetics from Short Trajectories": MWEM-main.zip: committor.py

```
import numpy as np
import mwem


milestones = [-80.0, -60.0, -40.0, -20.0, 0.0, 20.0, 40.0, 60.0, 80.0]

iters = [60, 70, 80, 90, 100]

for num_iter in iters:
    N_i_j = np.loadtxt('../N_i_j_files/N_i_j_%d.dat'%num_iter)
    K = mwem.K_matrix(N_i_j)
    #print(K)
    C = mwem.committor(K)

    f1 = open('committor_%d.dat'%num_iter,'w')

    for i in range(len(C)):
        print(milestones[i],C[i],file=f1)

    f1.close()
```
