## Supplementary material for "Markovian Weighted Ensemble Milestoning (M-WEM): Long-time Kinetics from Short Trajectories": MWEM-main.zip: plot-free-energy-landscape_40.ipynb

```
# %% [code]
import numpy as np
import pickle
import matplotlib.pyplot as plt
import scipy.interpolate

# %% [code]
#define function to convert long list to array
from itertools import chain
def longlist2array(longlist):
    flat = np.fromiter(chain.from_iterable(longlist), np.array(longlist[0][0]).dtype, -1) # Without intermediate list:)
    return flat.reshape((len(longlist), -1))

# %% [code]
p = np.loadtxt('cell_prob_40.dat')
p

# %% [code]
#trajs = []
#weights = []
cost = 0.0
probabilities = []
xedges = np.linspace(-175.0, 175.0, num=140)
for i in range(len(p)):
    a = pickle.load(open('cell_%d/trajectories_40.pkl'%i,'rb'))

    w = np.loadtxt('cell_%d/weights_40.txt'%i)

    N_traj = len(w)
    len_traj = len(a[0])


    weights_array = np.repeat(w,len_traj)
    #print(weights_array.shape)

    #patch all trajectories one after another
    trajs_stacked = [j for k in a for j in k]
    #trajs.append(trajs_stacked)
    trajs_stacked = longlist2array(trajs_stacked)
    #print(trajs_stacked.shape)
    cost += len(trajs_stacked)*10*1E-6
    
    histogm = plt.hist2d(trajs_stacked[:,0],trajs_stacked[:,1],bins=xedges,density=True,weights=weights_array)
    probabilities.append(histogm[0].transpose())

print("computational cost = ",cost,"ns")

# %% [code]
x = xedges
x = np.delete(x,-1)
y = xedges
y = np.delete(y,-1)

probs = np.zeros((len(x),len(y)))
for i in range(len(probabilities)):
    probs += probabilities[i]
fig = plt.figure()
cp = plt.contourf(x,y,probs, 40, cmap='nipy_spectral')
cbar = plt.colorbar(cp)
cbar.set_label('Probability density',fontsize=16)
plt.xlim(-100,100)
plt.ylabel("$\Psi (\circ)$",fontsize=16)
plt.xlabel("$\Phi (\circ)$",fontsize=16)
#plt.savefig('prob_no_rescale.png',dpi=200,bbox_inches = 'tight')

# %% [code]
x = xedges
x = np.delete(x,-1)
y = xedges
y = np.delete(y,-1)

probs = np.zeros((len(x),len(y)))
for i in range(len(probabilities)):
    probs += p[i,1]*probabilities[i]
fig = plt.figure()
cp = plt.contourf(x,y,probs, 40, cmap='nipy_spectral')
cbar = plt.colorbar(cp)
cbar.set_label('Probability density',fontsize=16)
plt.xlim(-100,100)
#plt.savefig('prob_rescaled.png',dpi=200,bbox_inches = 'tight')

# %% [code]
free_energy = -np.log(probs)
free_energy -= np.min(free_energy)

for i in range(len(free_energy)):
    for j in range(len(free_energy[i])):
        if free_energy[i,j] >= 14.0:
            free_energy[i,j] = np.inf
            
fig = plt.figure()
cp = plt.contourf(x,y,free_energy, 40, cmap='nipy_spectral')
plt.xlim(-100,100)
plt.clim(0.0,8)
cbar = plt.colorbar(cp)
cbar.set_label('Free Energy ($k_BT$)',fontsize=16)
#plt.contour(xi, yi, zi, m)
plt.ylabel("$\Psi (\circ)$",fontsize=16)
plt.xlabel("$\Phi (\circ)$",fontsize=16)
#plt.title("A6-DNA: PDB: 5UZF")
plt.savefig('freeE_no_rescale.png',dpi=200,bbox_inches = 'tight')

# %% [code]
free_energy = -np.log(probs)
free_energy -= np.min(free_energy)

for i in range(len(free_energy)):
    for j in range(len(free_energy[i])):
        if free_energy[i,j] >= 9.0:
            free_energy[i,j] = np.inf

fig = plt.figure()
cp = plt.contourf(x,y,free_energy, 40, cmap='nipy_spectral')
plt.xlim(-100,100)
plt.clim(0.0,8.5)
cbar = plt.colorbar(cp)
cbar.set_label('Free Energy ($k_BT$)',fontsize=16)
#plt.contour(xi, yi, zi, m)
plt.ylabel("$\Psi (\circ)$",fontsize=16)
plt.xlabel("$\Phi (\circ)$",fontsize=16)
#plt.title("A6-DNA: PDB: 5UZF")
#plt.savefig('freeE_rescaled.png',dpi=200,bbox_inches = 'tight')

# %% [code]
```
