## Supplementary material for "Markovian Weighted Ensemble Milestoning (M-WEM): Long-time Kinetics from Short Trajectories": MWEM-main.zip: plots_Tryp_Ben.ipynb

```
# %% [code]
import numpy as np
import matplotlib.pyplot as plt
import scipy.stats as st

# %% [code]
l = np.loadtxt('MFPT_convergence.dat')

# %% [code]
plt.plot(l[:,0], l[:,1], c='g', ls='--',lw=2, label='M-WEM' )

plt.fill_between(l[:,0], l[:,2], l[:,3], facecolor='xkcd:lightgreen')

#plt.yscale('log')
plt.xlabel('Number of iterations',fontsize=16)
plt.ylabel('Residence time (ms)',fontsize=16)
plt.ylim(-10,90)
plt.xticks(fontsize=14)
plt.yticks(fontsize=14)

plt.axhline(y=1.7,label='Experiment',color='b')
plt.axhline(y=12,label='SEEKR',color='k')
plt.axhline(y=5.6,label='MMVT SEEKR',color='r')
plt.axhline(y=16,color='r')
plt.legend(fontsize=14)
#plt.savefig('Trp_Ben_Residence-time-convergence.png',dpi=200,bbox_inches='tight')
plt.savefig('Trp_Ben_Residence-time-convergence-linear.pdf',bbox_inches='tight')

# %% [code]
plt.axhline(y=600,label='Experiment',color='b')
plt.axhspan(600-300,600+300,color='lightblue',alpha=0.4)
plt.axhline(y=83,label='SEEKR',color='k')
plt.axhspan(83-14,83+14,color='gray',alpha=0.4)
plt.axhline(y=174,label='MMVT SEEKR',color='r')
plt.axhspan(174-9,174+9,color='pink',alpha=0.4)
plt.axhline(y=62,color='r')
plt.axhspan(62-6,62+6,color='pink',alpha=0.4)


plt.plot(l[:,0], 1E+3/l[:,1], c='g', ls='--',lw=2, label='M-WEM' )

plt.fill_between(l[:,0], 1E+3/l[:,2], 1E+3/l[:,3], facecolor='xkcd:lightgreen')

plt.yscale('log')
plt.xlabel('Number of iterations',fontsize=16)
plt.ylabel('$k_{off}$ (s$^{-1}$)',fontsize=16)
plt.ylim(1,2000)
plt.xticks(fontsize=14)
plt.yticks(fontsize=14)


#plt.axhline(y=16,label='MMVT SEEKR')
plt.legend(fontsize=14,ncol=2)
#plt.savefig('Trp_Ben_k_off-convergence.png',dpi=200,bbox_inches='tight')
plt.savefig('Trp_Ben_k_off-convergence.pdf',bbox_inches='tight')

# %% [code]
#compute errors of free energy profile
prob1 = np.loadtxt('cell_probability/cell_prob_160.dat')
prob2 = np.loadtxt('cell_probability/cell_prob_170.dat')
prob3 = np.loadtxt('cell_probability/cell_prob_180.dat')
prob4 = np.loadtxt('cell_probability/cell_prob_190.dat')
prob5 = np.loadtxt('cell_probability/cell_prob_200.dat')

prob_mean = np.zeros(len(prob1))
prob_conf_low = np.zeros(len(prob1))
prob_conf_high = np.zeros(len(prob1))

for i in range(len(prob_mean)):
    a = [prob1[i,1],prob2[i,1],prob3[i,1],prob4[i,1],prob5[i,1]]
    a = np.array(a)
    prob_mean[i] = np.mean(a)
    conf = st.t.interval(alpha=0.95, df=len(a)-1, loc=np.mean(a), scale=st.sem(a))
    prob_conf_low[i] = conf[0]
    prob_conf_high[i] = conf[1]
    
G = -0.6*np.log(prob_mean)

G -= (G[0] - 1.5)

G_low = -0.6*np.log(prob_conf_low)

G_low -= (G_low[0] - 1.5)

G_high = -0.6*np.log(prob_conf_high)

G_high -= (G_high[0] - 1.5)
    


# %% [code]
#compute errors of free energy profile
recG1 = np.loadtxt('reconstructed_freeE_160.dat')
recG2 = np.loadtxt('reconstructed_freeE_170.dat')
recG3 = np.loadtxt('reconstructed_freeE_180.dat')
recG4 = np.loadtxt('reconstructed_freeE_190.dat')
recG5 = np.loadtxt('reconstructed_freeE_200.dat')

recG_mean = np.zeros(len(recG1))
recG_conf_low = np.zeros(len(recG1))
recG_conf_high = np.zeros(len(recG1))

for i in range(len(recG_mean)):
    a = [recG1[i,1],recG2[i,1],recG3[i,1],recG4[i,1],recG5[i,1]]
    a = np.array(a)
    recG_mean[i] = np.mean(a)
    conf = st.t.interval(alpha=0.95, df=len(a)-1, loc=np.mean(a), scale=st.sem(a))
    recG_conf_low[i] = conf[0]
    recG_conf_high[i] = conf[1]
    


#recG_mean -= (recG_mean[0] - 1.5)


#recG_conf_low -= (recG_conf_low[0] - 1.5)


#recG_conf_high -= (recG_conf_high[0] - 1.5)

# %% [code]
l = np.loadtxt('/home/dhiman/PhD/Markovian-WEM/WEM-Trypsin-Benzamidine/WTM_eABF_Tryp_Ben/convergence/mean_std_pmf.dat')


plt.plot(l[:,0], l[:,1], c='b', ls='-',lw=2,label='WTM-eABF' )

plt.fill_between(l[:,0], l[:,1]-l[:,2], l[:,1]+l[:,2], facecolor='lightblue')


plt.plot(prob1[:,0], G, 'g', ls='--',lw=2, label='M-WEM' )

plt.fill_between(prob1[:,0], G_low, G_high, facecolor='xkcd:lightgreen')

plt.plot(recG1[:,0], recG_mean, 'r', ls='--',lw=2, label='M-WEM reconstructed' )

plt.fill_between(recG1[:,0], recG_conf_low, recG_conf_high, facecolor='pink')


plt.xlabel('Protein-Ligand Distance ($\AA$)',fontsize=16)
plt.ylabel('Free Energy (kcal/mol)',fontsize=16)
#plt.legend(fontsize=16,ncol=1,loc="upper right")
plt.xticks(fontsize=14)
plt.yticks(fontsize=14)
plt.legend(fontsize=14)
plt.xlim(0,12)
plt.ylim(-1,14)
plt.savefig('Trp_Ben_1D-FreeE.pdf',bbox_inches='tight')
#plt.savefig('Trp_Ben_1D-FreeE.png',dpi=200,bbox_inches='tight')

# %% [code]
```
