## Supplementary material for "Markovian Weighted Ensemble Milestoning (M-WEM): Long-time Kinetics from Short Trajectories": MWEM-main.zip: calculate_coverage.sh

```
#!/bin/bash

touch fraction_coverage.dat

cd cell_2.5
rm all.dcd
cp ../cell_9/ligand_distribution.py .
cp ../cell_9/fraction_coverage.py .
cp ../cell_9/nowat.pdb .
mdconvert -o all.dcd -t nowat.pdb all.nc
python ligand_distribution.py
python fraction_coverage.py > ../fraction_coverage.dat
cd ..

cd cell_3.5
rm all.dcd
cp ../cell_9/ligand_distribution.py .
cp ../cell_9/fraction_coverage.py .
cp ../cell_9/nowat.pdb .
mdconvert -o all.dcd -t nowat.pdb all.nc
python ligand_distribution.py
python fraction_coverage.py >> ../fraction_coverage.dat
cd ..

for i in {0..8}
do
	cd cell_$i
	rm all.dcd
	cp ../cell_9/ligand_distribution.py .
	cp ../cell_9/fraction_coverage.py .
	cp ../cell_9/nowat.pdb .
	mdconvert -o all.dcd -t nowat.pdb all.nc
	python ligand_distribution.py
	python fraction_coverage.py >> ../fraction_coverage.dat
	cd ..
done
```
