## Supplementary material for "Markovian Weighted Ensemble Milestoning (M-WEM): Long-time Kinetics from Short Trajectories": MWEM-main.zip: combine_traj.sh

```
#!/bin/bash

#SBATCH --job-name=milestoning      ## job name
#SBATCH --account andricio_lab     ## account to charge
#SBATCH -p standard          ## partition/queue name
#SBATCH --nodes=1            ## (-N) number of nodes to use
#SBATCH --cpus-per-task=1    ## number of cores the job needs
#SBATCH -t 24:00:00
#SBATCH --mem=8gb
#SBATCH --error=slurm-%J.err ## error log file

#for i in {4..9}
#do
#	cd cell_$i
#	cp ../cell_0/combine.sh .
#	./combine.sh common_files/structure.pdb > combine.log
#	cd ..
#done

for i in {0..9}
do
	cd cell_$i
	./combine.sh common_files/structure.pdb > combine.log
	cd ..
done

cd cell_2.5
#cp ../cell_0/combine.sh .
./combine.sh common_files/structure.pdb > combine.log
cd ..

cd cell_3.5
#cp ../cell_0/combine.sh .
./combine.sh common_files/structure.pdb > combine.log
cd ..
```
