## Supplementary material for "Markovian Weighted Ensemble Milestoning (M-WEM): Long-time Kinetics from Short Trajectories": MWEM-main.zip: plot-ligand-coverage.ipynb

```
# %% [code]
import numpy as np
import matplotlib.pyplot as plt

# %% [code]
indices = [0,1,2,2.5,3,3.5,4,5,6,7,8,9]
milestones = [1.0,1.5,2.0,2.5,3.0,3.5,4.0,5.0,6.0,8.0,10.0,12.0,13.0]

fig, ax = plt.subplots(figsize=(10,10))

for i in range(len(indices)):
    l = np.loadtxt('cell_%s/ligand_positions_radial.dat'%str(indices[i]))

    r = l[:,0]
    theta = l[:,1]
    phi = l[:,2]

    x = r*np.cos(phi)
    y = r*np.sin(phi)

    plt.scatter(x,y,s=0.5)

for i in range(len(milestones)):
    
    circle1 = plt.Circle((0, 0), float(milestones[i]), color='k', fill=False)
    ax.add_patch(circle1)
    
plt.scatter([0],[0],s=2,color='k')
plt.ylim(-14,14)
plt.xlim(-14,14)
ax.axis('off')
#plt.show()
plt.savefig('ligand_coverage_300.png',dpi=200)

# %% [code]
```
