## Supplementary material for "Markovian Weighted Ensemble Milestoning (M-WEM): Long-time Kinetics from Short Trajectories": MWEM-main.zip: combine.sh

```
function removewat {
	local inp=$(mktemp)
	local d=$(dirname $3)
	local f=$(basename $3)
	cd $d || return
	cat << EOF >> $inp
	trajin $f
	strip :Cl-,WAT
	trajout nowat.nc
EOF
	cpptraj -p $2 -i $inp > /dev/null
	printf "%8d %8d %s\n" $1 $total "$d"
	rm $inp
}
export -f removewat
parm=$(realpath $1)
total=$(find . -name "seg.dcd" | wc -l)
export total
find . -name "seg.dcd" |  nl | \
	xargs -P4 -L 1 sh -c "removewat \$0 ${parm} \$1"

inp=`mktemp`
cat << EOF > $inp
parmstrip :Cl-,WAT
parmwrite out nowat.parm7
#autoimage
#trajout all.nc
EOF
cpptraj -p ${parm} -i $inp > /dev/null

find . -name "nowat.nc" | sed 's/^/trajin /' > $inp
cat << EOF >> $inp
autoimage
trajout all.nc
EOF
cpptraj -p nowat.parm7 -i $inp
find . -name "nowat.nc" -delete
unset total
unset removewat
```
