## Supplementary material for "Markovian Weighted Ensemble Milestoning (M-WEM): Long-time Kinetics from Short Trajectories": MWEM-main.zip: fraction_coverage.py

```
import numpy as np
import math as m

def cart2sph(x,y,z):
    XsqPlusYsq = x**2 + y**2
    r = m.sqrt(XsqPlusYsq + z**2)               # r
    elev = m.atan2(z,m.sqrt(XsqPlusYsq))     # theta
    az = m.atan2(y,x)                           # phi
    return r, elev, az

'''def cart2sphA(pts):
    return np.array([cart2sph(x,y,z) for x,y,z in pts])

def appendSpherical(xyz):
    np.hstack((xyz, cart2sphA(xyz)))
'''
l = np.loadtxt('ligand_positions.dat')

polar = np.zeros((len(l),3))

f1 = open('ligand_positions_radial.dat','w')
for i in range(len(l)):
    r, theta, phi = cart2sph(l[i,0],l[i,1],l[i,2])
    print(r, theta, phi,file=f1)
    polar[i,0] = r
    polar[i,1] = theta
    polar[i,2] = phi
f1.close()

hist, xedges, yedges = np.histogram2d(polar[:,1],polar[:,2],bins=[180,360],range=[[-3.14,0],[-6.28,0]])

#print(hist)

#elements for integral
dtheta = xedges[1] - xedges[0]
dphi = yedges[1] - yedges[0]

#print(xedges)
area = 0.0
total_area = 0.0
#Find occupied bins and corresponding 
for i in range(len(hist)):
    for j in range(len(hist[i])):
        theta = (xedges[i] + xedges[i+1])*0.5
        phi = (yedges[i] + yedges[i+1])*0.5
            #print(theta, phi, hist[i,j])
        total_area += np.sin(theta)*dtheta*dphi
        if hist[i,j] > 0.0:
            #theta = (xedges[i] + xedges[i+1])*0.5
            #phi = (yedges[i] + yedges[i+1])*0.5
            #print(theta, phi, hist[i,j])
            area += np.sin(theta)*dtheta*dphi
print(area/total_area)
```
