## Supplementary material for "Markovian Weighted Ensemble Milestoning (M-WEM): Long-time Kinetics from Short Trajectories": MWEM-main.zip: ligand_distribution.py

```
import numpy as np
import MDAnalysis as md

u = md.Universe('nowat.pdb','all.dcd')

protein = u.select_atoms("bynum 2479 2490 2500 2536 2719 2746 2770 2788 2795 2868 2927")

ligand = u.select_atoms("bynum 3222:3239")

f1 = open('ligand_positions.dat','w')
for ts in u.trajectory:
    a = protein.center_of_mass()
    b = ligand.center_of_mass()
    #r = np.linalg.norm(a-b)
    r = a-b

    print(r[0],r[1],r[2],file=f1)
f1.close()
```
