## Supplementary material for "Markovian Weighted Ensemble Milestoning (M-WEM): Long-time Kinetics from Short Trajectories": MWEM-main.zip: setup.cfg

```
# Helper file to handle all configs

[coverage:run]
# .coveragerc to control coverage.py and pytest-cov
omit =
    # Omit the tests
    */tests/*
    # Omit generated versioneer
    mwem/_version.py

[yapf]
# YAPF, in .style.yapf files this shows up as "[style]" header
COLUMN_LIMIT = 119
INDENT_WIDTH = 4
USE_TABS = False

[flake8]
# Flake8, PyFlakes, etc
max-line-length = 119

[versioneer]
# Automatic version numbering scheme
VCS = git
style = pep440
versionfile_source = mwem/_version.py
versionfile_build = mwem/_version.py
tag_prefix = ''

[aliases]
test = pytest
```
