## Supplementary material for "Markovian Weighted Ensemble Milestoning (M-WEM): Long-time Kinetics from Short Trajectories": MWEM-main.zip: setup.py

```
"""
MWEM
Markovian Weighted Ensemble Milestoning (MWEM)
"""
import sys
from setuptools import setup, find_packages
import versioneer

short_description = __doc__.split("\n")

# from https://github.com/pytest-dev/pytest-runner#conditional-requirement
needs_pytest = {'pytest', 'test', 'ptr'}.intersection(sys.argv)
pytest_runner = ['pytest-runner'] if needs_pytest else []

try:
    with open("README.md", "r") as handle:
        long_description = handle.read()
except:
    long_description = "\n".join(short_description[2:])


setup(
    # Self-descriptive entries which should always be present
    name='mwem',
    author='Dhiman Ray',
    author_email='',
    description=short_description[0],
    long_description=long_description,
    long_description_content_type="text/markdown",
    version=versioneer.get_version(),
    cmdclass=versioneer.get_cmdclass(),
    license='BSD-3-Clause',

    # Which Python importable modules should be included when your package is installed
    # Handled automatically by setuptools. Use 'exclude' to prevent some specific
    # subpackage(s) from being added, if needed
    packages=find_packages(),

    # Optional include package data to ship with your package
    # Customize MANIFEST.in if the general case does not suit your needs
    # Comment out this line to prevent the files from being packaged with your software
    include_package_data=True,

    # Allows `setup.py test` to work correctly with pytest
    setup_requires=[] + pytest_runner,

    # Additional entries you may want simply uncomment the lines you want and fill in the data
    # url='http://www.my_package.com',  # Website
    # install_requires=[],              # Required packages, pulls from pip if needed; do not use for Conda deployment
    # platforms=['Linux',
    #            'Mac OS-X',
    #            'Unix',
    #            'Windows'],            # Valid platforms your code works on, adjust to your flavor
    # python_requires=">=3.5",          # Python version restrictions

    # Manual control if final package is compressible or not, set False to prevent the .egg from being made
    # zip_safe=False,

)
```
