## Supplementary material for "Markovian Weighted Ensemble Milestoning (M-WEM): Long-time Kinetics from Short Trajectories": MWEM-main.zip: versioneer.py

```
# Version: 0.18

"""The Versioneer - like a rocketeer, but for versions.

The Versioneer
==============

* like a rocketeer, but for versions!
* https://github.com/warner/python-versioneer
* Brian Warner
* License: Public Domain
* Compatible With: python2.6, 2.7, 3.2, 3.3, 3.4, 3.5, 3.6, and pypy
* [![Latest Version]
(https://pypip.in/version/versioneer/badge.svg?style=flat)
](https://pypi.python.org/pypi/versioneer/)
* [![Build Status]
(https://travis-ci.org/warner/python-versioneer.png?branch=master)
](https://travis-ci.org/warner/python-versioneer)

This is a tool for managing a recorded version number in distutils-based
python projects. The goal is to remove the tedious and error-prone "update
the embedded version string" step from your release process. Making a new
release should be as easy as recording a new tag in your version-control
system, and maybe making new tarballs.


## Quick Install

* `pip install versioneer` to somewhere to your $PATH
* add a `[versioneer]` section to your setup.cfg (see below)
* run `versioneer install` in your source tree, commit the results

## Version Identifiers

Source trees come from a variety of places:

* a version-control system checkout (mostly used by developers)
* a nightly tarball, produced by build automation
* a snapshot tarball, produced by a web-based VCS browser, like github's
  "tarball from tag" feature
* a release tarball, produced by "setup.py sdist", distributed through PyPI

Within each source tree, the version identifier (either a string or a number,
this tool is format-agnostic) can come from a variety of places:

* ask the VCS tool itself, e.g. "git describe" (for checkouts), which knows
  about recent "tags" and an absolute revision-id
* the name of the directory into which the tarball was unpacked
* an expanded VCS keyword ($Id$, etc)
* a `_version.py` created by some earlier build step

For released software, the version identifier is closely related to a VCS
tag. Some projects use tag names that include more than just the version
string (e.g. "myproject-1.2" instead of just "1.2"), in which case the tool
needs to strip the tag prefix to extract the version identifier. For
unreleased software (between tags), the version identifier should provide
enough information to help developers recreate the same tree, while also
giving them an idea of roughly how old the tree is (after version 1.2, before
version 1.3). Many VCS systems can report a description that captures this,
for example `git describe --tags --dirty --always` reports things like
"0.7-1-g574ab98-dirty" to indicate that the checkout is one revision past the
0.7 tag, has a unique revision id of "574ab98", and is "dirty" (it has
uncommitted changes.

The version identifier is used for multiple purposes:

* to allow the module to self-identify its version: `myproject.__version__`
* to choose a name and prefix for a 'setup.py sdist' tarball

## Theory of Operation

Versioneer works by adding a special `_version.py` file into your source
tree, where your `__init__.py` can import it. This `_version.py` knows how to
dynamically ask the VCS tool for version information at import time.

`_version.py` also contains `$Revision$` markers, and the installation
process marks `_version.py` to have this marker rewritten with a tag name
during the `git archive` command. As a result, generated tarballs will
contain enough information to get the proper version.

To allow `setup.py` to compute a version too, a `versioneer.py` is added to
the top level of your source tree, next to `setup.py` and the `setup.cfg`
that configures it. This overrides several distutils/setuptools commands to
compute the version when invoked, and changes `setup.py build` and `setup.py
sdist` to replace `_version.py` with a small static file that contains just
the generated version data.

## Installation

See [INSTALL.md](./INSTALL.md) for detailed installation instructions.

## Version-String Flavors

Code which uses Versioneer can learn about its version string at runtime by
importing `_version` from your main `__init__.py` file and running the
`get_versions()` function. From the "outside" (e.g. in `setup.py`), you can
import the top-level `versioneer.py` and run `get_versions()`.

Both functions return a dictionary with different flavors of version
information:

* `['version']`: A condensed version string, rendered using the selected
  style. This is the most commonly used value for the project's version
  string. The default "pep440" style yields strings like `0.11`,
  `0.11+2.g1076c97`, or `0.11+2.g1076c97.dirty`. See the "Styles" section
  below for alternative styles.

* `['full-revisionid']`: detailed revision identifier. For Git, this is the
  full SHA1 commit id, e.g. "1076c978a8d3cfc70f408fe5974aa6c092c949ac".

* `['date']`: Date and time of the latest `HEAD` commit. For Git, it is the
  commit date in ISO 8601 format. This will be None if the date is not
  available.

* `['dirty']`: a boolean, True if the tree has uncommitted changes. Note that
  this is only accurate if run in a VCS checkout, otherwise it is likely to
  be False or None

* `['error']`: if the version string could not be computed, this will be set
  to a string describing the problem, otherwise it will be None. It may be
  useful to throw an exception in setup.py if this is set, to avoid e.g.
  creating tarballs with a version string of "unknown".

Some variants are more useful than others. Including `full-revisionid` in a
bug report should allow developers to reconstruct the exact code being tested
(or indicate the presence of local changes that should be shared with the
developers). `version` is suitable for display in an "about" box or a CLI
`--version` output: it can be easily compared against release notes and lists
of bugs fixed in various releases.

The installer adds the following text to your `__init__.py` to place a basic
version in `YOURPROJECT.__version__`:

    from ._version import get_versions
    __version__ = get_versions()['version']
    del get_versions

## Styles

The setup.cfg `style=` configuration controls how the VCS information is
rendered into a version string.

The default style, "pep440", produces a PEP440-compliant string, equal to the
un-prefixed tag name for actual releases, and containing an additional "local
version" section with more detail for in-between builds. For Git, this is
TAG[+DISTANCE.gHEX[.dirty]] , using information from `git describe --tags
--dirty --always`. For example "0.11+2.g1076c97.dirty" indicates that the
tree is like the "1076c97" commit but has uncommitted changes (".dirty"), and
that this commit is two revisions ("+2") beyond the "0.11" tag. For released
software (exactly equal to a known tag), the identifier will only contain the
stripped tag, e.g. "0.11".

Other styles are available. See [details.md](details.md) in the Versioneer
source tree for descriptions.

## Debugging

Versioneer tries to avoid fatal errors: if something goes wrong, it will tend
to return a version of "0+unknown". To investigate the problem, run `setup.py
version`, which will run the version-lookup code in a verbose mode, and will
display the full contents of `get_versions()` (including the `error` string,
which may help identify what went wrong).

## Known Limitations

Some situations are known to cause problems for Versioneer. This details the
most significant ones. More can be found on Github
[issues page](https://github.com/warner/python-versioneer/issues).

### Subprojects

Versioneer has limited support for source trees in which `setup.py` is not in
the root directory (e.g. `setup.py` and `.git/` are *not* siblings). The are
two common reasons why `setup.py` might not be in the root:

* Source trees which contain multiple subprojects, such as
  [Buildbot](https://github.com/buildbot/buildbot), which contains both
  "master" and "slave" subprojects, each with their own `setup.py`,
  `setup.cfg`, and `tox.ini`. Projects like these produce multiple PyPI
  distributions (and upload multiple independently-installable tarballs).
* Source trees whose main purpose is to contain a C library, but which also
  provide bindings to Python (and perhaps other langauges) in subdirectories.

Versioneer will look for `.git` in parent directories, and most operations
should get the right version string. However `pip` and `setuptools` have bugs
and implementation details which frequently cause `pip install .` from a
subproject directory to fail to find a correct version string (so it usually
defaults to `0+unknown`).

`pip install --editable .` should work correctly. `setup.py install` might
work too.

Pip-8.1.1 is known to have this problem, but hopefully it will get fixed in
some later version.

[Bug #38](https://github.com/warner/python-versioneer/issues/38) is tracking
this issue. The discussion in
[PR #61](https://github.com/warner/python-versioneer/pull/61) describes the
issue from the Versioneer side in more detail.
[pip PR#3176](https://github.com/pypa/pip/pull/3176) and
[pip PR#3615](https://github.com/pypa/pip/pull/3615) contain work to improve
pip to let Versioneer work correctly.

Versioneer-0.16 and earlier only looked for a `.git` directory next to the
`setup.cfg`, so subprojects were completely unsupported with those releases.

### Editable installs with setuptools <= 18.5

`setup.py develop` and `pip install --editable .` allow you to install a
project into a virtualenv once, then continue editing the source code (and
test) without re-installing after every change.

"Entry-point scripts" (`setup(entry_points={"console_scripts": ..})`) are a
convenient way to specify executable scripts that should be installed along
with the python package.

These both work as expected when using modern setuptools. When using
setuptools-18.5 or earlier, however, certain operations will cause
`pkg_resources.DistributionNotFound` errors when running the entrypoint
script, which must be resolved by re-installing the package. This happens
when the install happens with one version, then the egg_info data is
regenerated while a different version is checked out. Many setup.py commands
cause egg_info to be rebuilt (including `sdist`, `wheel`, and installing into
a different virtualenv), so this can be surprising.

[Bug #83](https://github.com/warner/python-versioneer/issues/83) describes
this one, but upgrading to a newer version of setuptools should probably
resolve it.

### Unicode version strings

While Versioneer works (and is continually tested) with both Python 2 and
Python 3, it is not entirely consistent with bytes-vs-unicode distinctions.
Newer releases probably generate unicode version strings on py2. It's not
clear that this is wrong, but it may be surprising for applications when then
write these strings to a network connection or include them in bytes-oriented
APIs like cryptographic checksums.

[Bug #71](https://github.com/warner/python-versioneer/issues/71) investigates
this question.


## Updating Versioneer

To upgrade your project to a new release of Versioneer, do the following:

* install the new Versioneer (`pip install -U versioneer` or equivalent)
* edit `setup.cfg`, if necessary, to include any new configuration settings
  indicated by the release notes. See [UPGRADING](./UPGRADING.md) for details.
* re-run `versioneer install` in your source tree, to replace
  `SRC/_version.py`
* commit any changed files

## Future Directions

This tool is designed to make it easily extended to other version-control
systems: all VCS-specific components are in separate directories like
src/git/ . The top-level `versioneer.py` script is assembled from these
components by running make-versioneer.py . In the future, make-versioneer.py
will take a VCS name as an argument, and will construct a version of
`versioneer.py` that is specific to the given VCS. It might also take the
configuration arguments that are currently provided manually during
installation by editing setup.py . Alternatively, it might go the other
direction and include code from all supported VCS systems, reducing the
number of intermediate scripts.


## License

To make Versioneer easier to embed, all its code is dedicated to the public
domain. The `_version.py` that it creates is also in the public domain.
Specifically, both are released under the Creative Commons "Public Domain
Dedication" license (CC0-1.0), as described in
https://creativecommons.org/publicdomain/zero/1.0/ .

"""

from __future__ import print_function
try:
    import configparser
except ImportError:
    import ConfigParser as configparser
import errno
import json
import os
import re
import subprocess
import sys


class VersioneerConfig:
    """Container for Versioneer configuration parameters."""


def get_root():
    """Get the project root directory.

    We require that all commands are run from the project root, i.e. the
    directory that contains setup.py, setup.cfg, and versioneer.py .
    """
    root = os.path.realpath(os.path.abspath(os.getcwd()))
    setup_py = os.path.join(root, "setup.py")
    versioneer_py = os.path.join(root, "versioneer.py")
    if not (os.path.exists(setup_py) or os.path.exists(versioneer_py)):
        # allow 'python path/to/setup.py COMMAND'
        root = os.path.dirname(os.path.realpath(os.path.abspath(sys.argv[0])))
        setup_py = os.path.join(root, "setup.py")
        versioneer_py = os.path.join(root, "versioneer.py")
    if not (os.path.exists(setup_py) or os.path.exists(versioneer_py)):
        err = ("Versioneer was unable to run the project root directory. "
               "Versioneer requires setup.py to be executed from "
               "its immediate directory (like 'python setup.py COMMAND'), "
               "or in a way that lets it use sys.argv[0] to find the root "
               "(like 'python path/to/setup.py COMMAND').")
        raise VersioneerBadRootError(err)
    try:
        # Certain runtime workflows (setup.py install/develop in a setuptools
        # tree) execute all dependencies in a single python process, so
        # "versioneer" may be imported multiple times, and python's shared
        # module-import table will cache the first one. So we can't use
        # os.path.dirname(__file__), as that will find whichever
        # versioneer.py was first imported, even in later projects.
        me = os.path.realpath(os.path.abspath(__file__))
        me_dir = os.path.normcase(os.path.splitext(me)[0])
        vsr_dir = os.path.normcase(os.path.splitext(versioneer_py)[0])
        if me_dir != vsr_dir:
            print("Warning: build in %s is using versioneer.py from %s"
                  % (os.path.dirname(me), versioneer_py))
    except NameError:
        pass
    return root


def get_config_from_root(root):
    """Read the project setup.cfg file to determine Versioneer config."""
    # This might raise EnvironmentError (if setup.cfg is missing), or
    # configparser.NoSectionError (if it lacks a [versioneer] section), or
    # configparser.NoOptionError (if it lacks "VCS="). See the docstring at
    # the top of versioneer.py for instructions on writing your setup.cfg .
    setup_cfg = os.path.join(root, "setup.cfg")
    parser = configparser.SafeConfigParser()
    with open(setup_cfg, "r") as f:
        parser.readfp(f)
    VCS = parser.get("versioneer", "VCS")  # mandatory

    def get(parser, name):
        if parser.has_option("versioneer", name):
            return parser.get("versioneer", name)
        return None
    cfg = VersioneerConfig()
    cfg.VCS = VCS
    cfg.style = get(parser, "style") or ""
    cfg.versionfile_source = get(parser, "versionfile_source")
    cfg.versionfile_build = get(parser, "versionfile_build")
    cfg.tag_prefix = get(parser, "tag_prefix")
    if cfg.tag_prefix in ("''", '""'):
        cfg.tag_prefix = ""
    cfg.parentdir_prefix = get(parser, "parentdir_prefix")
    cfg.verbose = get(parser, "verbose")
    return cfg


class NotThisMethod(Exception):
    """Exception raised if a method is not valid for the current scenario."""


# these dictionaries contain VCS-specific tools
LONG_VERSION_PY = {}
HANDLERS = {}


def register_vcs_handler(vcs, method):  # decorator
    """Decorator to mark a method as the handler for a particular VCS."""
    def decorate(f):
        """Store f in HANDLERS[vcs][method]."""
        if vcs not in HANDLERS:
            HANDLERS[vcs] = {}
        HANDLERS[vcs][method] = f
        return f
    return decorate


def run_command(commands, args, cwd=None, verbose=False, hide_stderr=False,
                env=None):
    """Call the given command(s)."""
    assert isinstance(commands, list)
    p = None
    for c in commands:
        try:
            dispcmd = str([c] + args)
            # remember shell=False, so use git.cmd on windows, not just git
            p = subprocess.Popen([c] + args, cwd=cwd, env=env,
                                 stdout=subprocess.PIPE,
                                 stderr=(subprocess.PIPE if hide_stderr
                                         else None))
            break
        except EnvironmentError:
            e = sys.exc_info()[1]
            if e.errno == errno.ENOENT:
                continue
            if verbose:
                print("unable to run %s" % dispcmd)
                print(e)
            return None, None
    else:
        if verbose:
            print("unable to find command, tried %s" % (commands,))
        return None, None
    stdout = p.communicate()[0].strip()
    if sys.version_info[0] >= 3:
        stdout = stdout.decode()
    if p.returncode != 0:
        if verbose:
            print("unable to run %s (error)" % dispcmd)
            print("stdout was %s" % stdout)
        return None, p.returncode
    return stdout, p.returncode


LONG_VERSION_PY['git'] = '''
# This file helps to compute a version number in source trees obtained from
# git-archive tarball (such as those provided by githubs download-from-tag
# feature). Distribution tarballs (built by setup.py sdist) and build
# directories (produced by setup.py build) will contain a much shorter file
# that just contains the computed version number.

# This file is released into the public domain. Generated by
# versioneer-0.18 (https://github.com/warner/python-versioneer)

"""Git implementation of _version.py."""

import errno
import os
import re
import subprocess
import sys


def get_keywords():
    """Get the keywords needed to look up the version information."""
    # these strings will be replaced by git during git-archive.
    # setup.py/versioneer.py will grep for the variable names, so they must
    # each be defined on a line of their own. _version.py will just call
    # get_keywords().
    git_refnames = "%(DOLLAR)sFormat:%%d%(DOLLAR)s"
    git_full = "%(DOLLAR)sFormat:%%H%(DOLLAR)s"
    git_date = "%(DOLLAR)sFormat:%%ci%(DOLLAR)s"
    keywords = {"refnames": git_refnames, "full": git_full, "date": git_date}
    return keywords


class VersioneerConfig:
    """Container for Versioneer configuration parameters."""


def get_config():
    """Create, populate and return the VersioneerConfig() object."""
    # these strings are filled in when 'setup.py versioneer' creates
    # _version.py
    cfg = VersioneerConfig()
    cfg.VCS = "git"
    cfg.style = "%(STYLE)s"
    cfg.tag_prefix = "%(TAG_PREFIX)s"
    cfg.parentdir_prefix = "%(PARENTDIR_PREFIX)s"
    cfg.versionfile_source = "%(VERSIONFILE_SOURCE)s"
    cfg.verbose = False
    return cfg


class NotThisMethod(Exception):
    """Exception raised if a method is not valid for the current scenario."""


LONG_VERSION_PY = {}
HANDLERS = {}


def register_vcs_handler(vcs, method):  # decorator
    """Decorator to mark a method as the handler for a particular VCS."""
    def decorate(f):
        """Store f in HANDLERS[vcs][method]."""
        if vcs not in HANDLERS:
            HANDLERS[vcs] = {}
        HANDLERS[vcs][method] = f
        return f
    return decorate


def run_command(commands, args, cwd=None, verbose=False, hide_stderr=False,
                env=None):
    """Call the given command(s)."""
    assert isinstance(commands, list)
    p = None
    for c in commands:
        try:
            dispcmd = str([c] + args)
            # remember shell=False, so use git.cmd on windows, not just git
            p = subprocess.Popen([c] + args, cwd=cwd, env=env,
                                 stdout=subprocess.PIPE,
                                 stderr=(subprocess.PIPE if hide_stderr
                                         else None))
            break
        except EnvironmentError:
            e = sys.exc_info()[1]
            if e.errno == errno.ENOENT:
                continue
            if verbose:
                print("unable to run %%s" %% dispcmd)
                print(e)
            return None, None
    else:
        if verbose:
            print("unable to find command, tried %%s" %% (commands,))
        return None, None
    stdout = p.communicate()[0].strip()
    if sys.version_info[0] >= 3:
        stdout = stdout.decode()
    if p.returncode != 0:
        if verbose:
            print("unable to run %%s (error)" %% dispcmd)
            print("stdout was %%s" %% stdout)
        return None, p.returncode
    return stdout, p.returncode


def versions_from_parentdir(parentdir_prefix, root, verbose):
    """Try to determine the version from the parent directory name.

    Source tarballs conventionally unpack into a directory that includes both
    the project name and a version string. We will also support searching up
    two directory levels for an appropriately named parent directory
    """
    rootdirs = []

    for i in range(3):
        dirname = os.path.basename(root)
        if dirname.startswith(parentdir_prefix):
            return {"version": dirname[len(parentdir_prefix):],
                    "full-revisionid": None,
                    "dirty": False, "error": None, "date": None}
        else:
            rootdirs.append(root)
            root = os.path.dirname(root)  # up a level

    if verbose:
        print("Tried directories %%s but none started with prefix %%s" %%
              (str(rootdirs), parentdir_prefix))
    raise NotThisMethod("rootdir doesn't start with parentdir_prefix")


@register_vcs_handler("git", "get_keywords")
def git_get_keywords(versionfile_abs):
    """Extract version information from the given file."""
    # the code embedded in _version.py can just fetch the value of these
    # keywords. When used from setup.py, we don't want to import _version.py,
    # so we do it with a regexp instead. This function is not used from
    # _version.py.
    keywords = {}
    try:
        f = open(versionfile_abs, "r")
        for line in f.readlines():
            if line.strip().startswith("git_refnames ="):
                mo = re.search(r'=\s*"(.*)"', line)
                if mo:
                    keywords["refnames"] = mo.group(1)
            if line.strip().startswith("git_full ="):
                mo = re.search(r'=\s*"(.*)"', line)
                if mo:
                    keywords["full"] = mo.group(1)
            if line.strip().startswith("git_date ="):
                mo = re.search(r'=\s*"(.*)"', line)
                if mo:
                    keywords["date"] = mo.group(1)
        f.close()
    except EnvironmentError:
        pass
    return keywords


@register_vcs_handler("git", "keywords")
def git_versions_from_keywords(keywords, tag_prefix, verbose):
    """Get version information from git keywords."""
    if not keywords:
        raise NotThisMethod("no keywords at all, weird")
    date = keywords.get("date")
    if date is not None:
        # git-2.2.0 added "%%cI", which expands to an ISO-8601 -compliant
        # datestamp. However we prefer "%%ci" (which expands to an "ISO-8601
        # -like" string, which we must then edit to make compliant), because
        # it's been around since git-1.5.3, and it's too difficult to
        # discover which version we're using, or to work around using an
        # older one.
        date = date.strip().replace(" ", "T", 1).replace(" ", "", 1)
    refnames = keywords["refnames"].strip()
    if refnames.startswith("$Format"):
        if verbose:
            print("keywords are unexpanded, not using")
        raise NotThisMethod("unexpanded keywords, not a git-archive tarball")
    refs = set([r.strip() for r in refnames.strip("()").split(",")])
    # starting in git-1.8.3, tags are listed as "tag: foo-1.0" instead of
    # just "foo-1.0". If we see a "tag: " prefix, prefer those.
    TAG = "tag: "
    tags = set([r[len(TAG):] for r in refs if r.startswith(TAG)])
    if not tags:
        # Either we're using git < 1.8.3, or there really are no tags. We use
        # a heuristic: assume all version tags have a digit. The old git %%d
        # expansion behaves like git log --decorate=short and strips out the
        # refs/heads/ and refs/tags/ prefixes that would let us distinguish
        # between branches and tags. By ignoring refnames without digits, we
        # filter out many common branch names like "release" and
        # "stabilization", as well as "HEAD" and "master".
        tags = set([r for r in refs if re.search(r'\d', r)])
        if verbose:
            print("discarding '%%s', no digits" %% ",".join(refs - tags))
    if verbose:
        print("likely tags: %%s" %% ",".join(sorted(tags)))
    for ref in sorted(tags):
        # sorting will prefer e.g. "2.0" over "2.0rc1"
        if ref.startswith(tag_prefix):
            r = ref[len(tag_prefix):]
            if verbose:
                print("picking %%s" %% r)
            return {"version": r,
                    "full-revisionid": keywords["full"].strip(),
                    "dirty": False, "error": None,
                    "date": date}
    # no suitable tags, so version is "0+unknown", but full hex is still there
    if verbose:
        print("no suitable tags, using unknown + full revision id")
    return {"version": "0+unknown",
            "full-revisionid": keywords["full"].strip(),
            "dirty": False, "error": "no suitable tags", "date": None}


@register_vcs_handler("git", "pieces_from_vcs")
def git_pieces_from_vcs(tag_prefix, root, verbose, run_command=run_command):
    """Get version from 'git describe' in the root of the source tree.

    This only gets called if the git-archive 'subst' keywords were *not*
    expanded, and _version.py hasn't already been rewritten with a short
    version string, meaning we're inside a checked out source tree.
    """
    GITS = ["git"]
    if sys.platform == "win32":
        GITS = ["git.cmd", "git.exe"]

    out, rc = run_command(GITS, ["rev-parse", "--git-dir"], cwd=root,
                          hide_stderr=True)
    if rc != 0:
        if verbose:
            print("Directory %%s not under git control" %% root)
        raise NotThisMethod("'git rev-parse --git-dir' returned error")

    # if there is a tag matching tag_prefix, this yields TAG-NUM-gHEX[-dirty]
    # if there isn't one, this yields HEX[-dirty] (no NUM)
    describe_out, rc = run_command(GITS, ["describe", "--tags", "--dirty",
                                          "--always", "--long",
                                          "--match", "%%s*" %% tag_prefix],
                                   cwd=root)
    # --long was added in git-1.5.5
    if describe_out is None:
        raise NotThisMethod("'git describe' failed")
    describe_out = describe_out.strip()
    full_out, rc = run_command(GITS, ["rev-parse", "HEAD"], cwd=root)
    if full_out is None:
        raise NotThisMethod("'git rev-parse' failed")
    full_out = full_out.strip()

    pieces = {}
    pieces["long"] = full_out
    pieces["short"] = full_out[:7]  # maybe improved later
    pieces["error"] = None

    # parse describe_out. It will be like TAG-NUM-gHEX[-dirty] or HEX[-dirty]
    # TAG might have hyphens.
    git_describe = describe_out

    # look for -dirty suffix
    dirty = git_describe.endswith("-dirty")
    pieces["dirty"] = dirty
    if dirty:
        git_describe = git_describe[:git_describe.rindex("-dirty")]

    # now we have TAG-NUM-gHEX or HEX

    if "-" in git_describe:
        # TAG-NUM-gHEX
        mo = re.search(r'^(.+)-(\d+)-g([0-9a-f]+)$', git_describe)
        if not mo:
            # unparseable. Maybe git-describe is misbehaving?
            pieces["error"] = ("unable to parse git-describe output: '%%s'"
                               %% describe_out)
            return pieces

        # tag
        full_tag = mo.group(1)
        if not full_tag.startswith(tag_prefix):
            if verbose:
                fmt = "tag '%%s' doesn't start with prefix '%%s'"
                print(fmt %% (full_tag, tag_prefix))
            pieces["error"] = ("tag '%%s' doesn't start with prefix '%%s'"
                               %% (full_tag, tag_prefix))
            return pieces
        pieces["closest-tag"] = full_tag[len(tag_prefix):]

        # distance: number of commits since tag
        pieces["distance"] = int(mo.group(2))

        # commit: short hex revision ID
        pieces["short"] = mo.group(3)

    else:
        # HEX: no tags
        pieces["closest-tag"] = None
        count_out, rc = run_command(GITS, ["rev-list", "HEAD", "--count"],
                                    cwd=root)
        pieces["distance"] = int(count_out)  # total number of commits

    # commit date: see ISO-8601 comment in git_versions_from_keywords()
    date = run_command(GITS, ["show", "-s", "--format=%%ci", "HEAD"],
                       cwd=root)[0].strip()
    pieces["date"] = date.strip().replace(" ", "T", 1).replace(" ", "", 1)

    return pieces


def plus_or_dot(pieces):
    """Return a + if we don't already have one, else return a ."""
    if "+" in pieces.get("closest-tag", ""):
        return "."
    return "+"


def render_pep440(pieces):
    """Build up version string, with post-release "local version identifier".

    Our goal: TAG[+DISTANCE.gHEX[.dirty]] . Note that if you
    get a tagged build and then dirty it, you'll get TAG+0.gHEX.dirty

    Exceptions:
    1: no tags. git_describe was just HEX. 0+untagged.DISTANCE.gHEX[.dirty]
    """
    if pieces["closest-tag"]:
        rendered = pieces["closest-tag"]
        if pieces["distance"] or pieces["dirty"]:
            rendered += plus_or_dot(pieces)
            rendered += "%%d.g%%s" %% (pieces["distance"], pieces["short"])
            if pieces["dirty"]:
                rendered += ".dirty"
    else:
        # exception #1
        rendered = "0+untagged.%%d.g%%s" %% (pieces["distance"],
                                          pieces["short"])
        if pieces["dirty"]:
            rendered += ".dirty"
    return rendered


def render_pep440_pre(pieces):
    """TAG[.post.devDISTANCE] -- No -dirty.

    Exceptions:
    1: no tags. 0.post.devDISTANCE
    """
    if pieces["closest-tag"]:
        rendered = pieces["closest-tag"]
        if pieces["distance"]:
            rendered += ".post.dev%%d" %% pieces["distance"]
    else:
        # exception #1
        rendered = "0.post.dev%%d" %% pieces["distance"]
    return rendered


def render_pep440_post(pieces):
    """TAG[.postDISTANCE[.dev0]+gHEX] .

    The ".dev0" means dirty. Note that .dev0 sorts backwards
    (a dirty tree will appear "older" than the corresponding clean one),
    but you shouldn't be releasing software with -dirty anyways.

    Exceptions:
    1: no tags. 0.postDISTANCE[.dev0]
    """
    if pieces["closest-tag"]:
        rendered = pieces["closest-tag"]
        if pieces["distance"] or pieces["dirty"]:
            rendered += ".post%%d" %% pieces["distance"]
            if pieces["dirty"]:
                rendered += ".dev0"
            rendered += plus_or_dot(pieces)
            rendered += "g%%s" %% pieces["short"]
    else:
        # exception #1
        rendered = "0.post%%d" %% pieces["distance"]
        if pieces["dirty"]:
            rendered += ".dev0"
        rendered += "+g%%s" %% pieces["short"]
    return rendered


def render_pep440_old(pieces):
    """TAG[.postDISTANCE[.dev0]] .

    The ".dev0" means dirty.

    Eexceptions:
    1: no tags. 0.postDISTANCE[.dev0]
    """
    if pieces["closest-tag"]:
        rendered = pieces["closest-tag"]
        if pieces["distance"] or pieces["dirty"]:
            rendered += ".post%%d" %% pieces["distance"]
            if pieces["dirty"]:
                rendered += ".dev0"
    else:
        # exception #1
        rendered = "0.post%%d" %% pieces["distance"]
        if pieces["dirty"]:
            rendered += ".dev0"
    return rendered


def render_git_describe(pieces):
    """TAG[-DISTANCE-gHEX][-dirty].

    Like 'git describe --tags --dirty --always'.

    Exceptions:
    1: no tags. HEX[-dirty]  (note: no 'g' prefix)
    """
    if pieces["closest-tag"]:
        rendered = pieces["closest-tag"]
        if pieces["distance"]:
            rendered += "-%%d-g%%s" %% (pieces["distance"], pieces["short"])
    else:
        # exception #1
        rendered = pieces["short"]
    if pieces["dirty"]:
        rendered += "-dirty"
    return rendered


def render_git_describe_long(pieces):
    """TAG-DISTANCE-gHEX[-dirty].

    Like 'git describe --tags --dirty --always -long'.
    The distance/hash is unconditional.

    Exceptions:
    1: no tags. HEX[-dirty]  (note: no 'g' prefix)
    """
    if pieces["closest-tag"]:
        rendered = pieces["closest-tag"]
        rendered += "-%%d-g%%s" %% (pieces["distance"], pieces["short"])
    else:
        # exception #1
        rendered = pieces["short"]
    if pieces["dirty"]:
        rendered += "-dirty"
    return rendered


def render(pieces, style):
    """Render the given version pieces into the requested style."""
    if pieces["error"]:
        return {"version": "unknown",
                "full-revisionid": pieces.get("long"),
                "dirty": None,
                "error": pieces["error"],
                "date": None}

    if not style or style == "default":
        style = "pep440"  # the default

    if style == "pep440":
        rendered = render_pep440(pieces)
    elif style == "pep440-pre":
        rendered = render_pep440_pre(pieces)
    elif style == "pep440-post":
        rendered = render_pep440_post(pieces)
    elif style == "pep440-old":
        rendered = render_pep440_old(pieces)
    elif style == "git-describe":
        rendered = render_git_describe(pieces)
    elif style == "git-describe-long":
        rendered = render_git_describe_long(pieces)
    else:
        raise ValueError("unknown style '%%s'" %% style)

    return {"version": rendered, "full-revisionid": pieces["long"],
            "dirty": pieces["dirty"], "error": None,
            "date": pieces.get("date")}


def get_versions():
    """Get version information or return default if unable to do so."""
    # I am in _version.py, which lives at ROOT/VERSIONFILE_SOURCE. If we have
    # __file__, we can work backwards from there to the root. Some
    # py2exe/bbfreeze/non-CPython implementations don't do __file__, in which
    # case we can only use expanded keywords.

    cfg = get_config()
    verbose = cfg.verbose

    try:
        return git_versions_from_keywords(get_keywords(), cfg.tag_prefix,
                                          verbose)
    except NotThisMethod:
        pass

    try:
        root = os.path.realpath(__file__)
        # versionfile_source is the relative path from the top of the source
        # tree (where the .git directory might live) to this file. Invert
        # this to find the root from __file__.
        for i in cfg.versionfile_source.split('/'):
            root = os.path.dirname(root)
    except NameError:
        return {"version": "0+unknown", "full-revisionid": None,
                "dirty": None,
                "error": "unable to find root of source tree",
                "date": None}

    try:
        pieces = git_pieces_from_vcs(cfg.tag_prefix, root, verbose)
        return render(pieces, cfg.style)
    except NotThisMethod:
        pass

    try:
        if cfg.parentdir_prefix:
            return versions_from_parentdir(cfg.parentdir_prefix, root, verbose)
    except NotThisMethod:
        pass

    return {"version": "0+unknown", "full-revisionid": None,
            "dirty": None,
            "error": "unable to compute version", "date": None}
'''


@register_vcs_handler("git", "get_keywords")
def git_get_keywords(versionfile_abs):
    """Extract version information from the given file."""
    # the code embedded in _version.py can just fetch the value of these
    # keywords. When used from setup.py, we don't want to import _version.py,
    # so we do it with a regexp instead. This function is not used from
    # _version.py.
    keywords = {}
    try:
        f = open(versionfile_abs, "r")
        for line in f.readlines():
            if line.strip().startswith("git_refnames ="):
                mo = re.search(r'=\s*"(.*)"', line)
                if mo:
                    keywords["refnames"] = mo.group(1)
            if line.strip().startswith("git_full ="):
                mo = re.search(r'=\s*"(.*)"', line)
                if mo:
                    keywords["full"] = mo.group(1)
            if line.strip().startswith("git_date ="):
                mo = re.search(r'=\s*"(.*)"', line)
                if mo:
                    keywords["date"] = mo.group(1)
        f.close()
    except EnvironmentError:
        pass
    return keywords


@register_vcs_handler("git", "keywords")
def git_versions_from_keywords(keywords, tag_prefix, verbose):
    """Get version information from git keywords."""
    if not keywords:
        raise NotThisMethod("no keywords at all, weird")
    date = keywords.get("date")
    if date is not None:
        # git-2.2.0 added "%cI", which expands to an ISO-8601 -compliant
        # datestamp. However we prefer "%ci" (which expands to an "ISO-8601
        # -like" string, which we must then edit to make compliant), because
        # it's been around since git-1.5.3, and it's too difficult to
        # discover which version we're using, or to work around using an
        # older one.
        date = date.strip().replace(" ", "T", 1).replace(" ", "", 1)
    refnames = keywords["refnames"].strip()
    if refnames.startswith("$Format"):
        if verbose:
            print("keywords are unexpanded, not using")
        raise NotThisMethod("unexpanded keywords, not a git-archive tarball")
    refs = set([r.strip() for r in refnames.strip("()").split(",")])
    # starting in git-1.8.3, tags are listed as "tag: foo-1.0" instead of
    # just "foo-1.0". If we see a "tag: " prefix, prefer those.
    TAG = "tag: "
    tags = set([r[len(TAG):] for r in refs if r.startswith(TAG)])
    if not tags:
        # Either we're using git < 1.8.3, or there really are no tags. We use
        # a heuristic: assume all version tags have a digit. The old git %d
        # expansion behaves like git log --decorate=short and strips out the
        # refs/heads/ and refs/tags/ prefixes that would let us distinguish
        # between branches and tags. By ignoring refnames without digits, we
        # filter out many common branch names like "release" and
        # "stabilization", as well as "HEAD" and "master".
        tags = set([r for r in refs if re.search(r'\d', r)])
        if verbose:
            print("discarding '%s', no digits" % ",".join(refs - tags))
    if verbose:
        print("likely tags: %s" % ",".join(sorted(tags)))
    for ref in sorted(tags):
        # sorting will prefer e.g. "2.0" over "2.0rc1"
        if ref.startswith(tag_prefix):
            r = ref[len(tag_prefix):]
            if verbose:
                print("picking %s" % r)
            return {"version": r,
                    "full-revisionid": keywords["full"].strip(),
                    "dirty": False, "error": None,
                    "date": date}
    # no suitable tags, so version is "0+unknown", but full hex is still there
    if verbose:
        print("no suitable tags, using unknown + full revision id")
    return {"version": "0+unknown",
            "full-revisionid": keywords["full"].strip(),
            "dirty": False, "error": "no suitable tags", "date": None}


@register_vcs_handler("git", "pieces_from_vcs")
def git_pieces_from_vcs(tag_prefix, root, verbose, run_command=run_command):
    """Get version from 'git describe' in the root of the source tree.

    This only gets called if the git-archive 'subst' keywords were *not*
    expanded, and _version.py hasn't already been rewritten with a short
    version string, meaning we're inside a checked out source tree.
    """
    GITS = ["git"]
    if sys.platform == "win32":
        GITS = ["git.cmd", "git.exe"]

    out, rc = run_command(GITS, ["rev-parse", "--git-dir"], cwd=root,
                          hide_stderr=True)
    if rc != 0:
        if verbose:
            print("Directory %s not under git control" % root)
        raise NotThisMethod("'git rev-parse --git-dir' returned error")

    # if there is a tag matching tag_prefix, this yields TAG-NUM-gHEX[-dirty]
    # if there isn't one, this yields HEX[-dirty] (no NUM)
    describe_out, rc = run_command(GITS, ["describe", "--tags", "--dirty",
                                          "--always", "--long",
                                          "--match", "%s*" % tag_prefix],
                                   cwd=root)
    # --long was added in git-1.5.5
    if describe_out is None:
        raise NotThisMethod("'git describe' failed")
    describe_out = describe_out.strip()
    full_out, rc = run_command(GITS, ["rev-parse", "HEAD"], cwd=root)
    if full_out is None:
        raise NotThisMethod("'git rev-parse' failed")
    full_out = full_out.strip()

    pieces = {}
    pieces["long"] = full_out
    pieces["short"] = full_out[:7]  # maybe improved later
    pieces["error"] = None

    # parse describe_out. It will be like TAG-NUM-gHEX[-dirty] or HEX[-dirty]
    # TAG might have hyphens.
    git_describe = describe_out

    # look for -dirty suffix
    dirty = git_describe.endswith("-dirty")
    pieces["dirty"] = dirty
    if dirty:
        git_describe = git_describe[:git_describe.rindex("-dirty")]

    # now we have TAG-NUM-gHEX or HEX

    if "-" in git_describe:
        # TAG-NUM-gHEX
        mo = re.search(r'^(.+)-(\d+)-g([0-9a-f]+)$', git_describe)
        if not mo:
            # unparseable. Maybe git-describe is misbehaving?
            pieces["error"] = ("unable to parse git-describe output: '%s'"
                               % describe_out)
            return pieces

        # tag
        full_tag = mo.group(1)
        if not full_tag.startswith(tag_prefix):
            if verbose:
                fmt = "tag '%s' doesn't start with prefix '%s'"
                print(fmt % (full_tag, tag_prefix))
            pieces["error"] = ("tag '%s' doesn't start with prefix '%s'"
                               % (full_tag, tag_prefix))
            return pieces
        pieces["closest-tag"] = full_tag[len(tag_prefix):]

        # distance: number of commits since tag
        pieces["distance"] = int(mo.group(2))

        # commit: short hex revision ID
        pieces["short"] = mo.group(3)

    else:
        # HEX: no tags
        pieces["closest-tag"] = None
        count_out, rc = run_command(GITS, ["rev-list", "HEAD", "--count"],
                                    cwd=root)
        pieces["distance"] = int(count_out)  # total number of commits

    # commit date: see ISO-8601 comment in git_versions_from_keywords()
    date = run_command(GITS, ["show", "-s", "--format=%ci", "HEAD"],
                       cwd=root)[0].strip()
    pieces["date"] = date.strip().replace(" ", "T", 1).replace(" ", "", 1)

    return pieces


def do_vcs_install(manifest_in, versionfile_source, ipy):
    """Git-specific installation logic for Versioneer.

    For Git, this means creating/changing .gitattributes to mark _version.py
    for export-subst keyword substitution.
    """
    GITS = ["git"]
    if sys.platform == "win32":
        GITS = ["git.cmd", "git.exe"]
    files = [manifest_in, versionfile_source]
    if ipy:
        files.append(ipy)
    try:
        me = __file__
        if me.endswith(".pyc") or me.endswith(".pyo"):
            me = os.path.splitext(me)[0] + ".py"
        versioneer_file = os.path.relpath(me)
    except NameError:
        versioneer_file = "versioneer.py"
    files.append(versioneer_file)
    present = False
    try:
        f = open(".gitattributes", "r")
        for line in f.readlines():
            if line.strip().startswith(versionfile_source):
                if "export-subst" in line.strip().split()[1:]:
                    present = True
        f.close()
    except EnvironmentError:
        pass
    if not present:
        f = open(".gitattributes", "a+")
        f.write("%s export-subst\n" % versionfile_source)
        f.close()
        files.append(".gitattributes")
    run_command(GITS, ["add", "--"] + files)


def versions_from_parentdir(parentdir_prefix, root, verbose):
    """Try to determine the version from the parent directory name.

    Source tarballs conventionally unpack into a directory that includes both
    the project name and a version string. We will also support searching up
    two directory levels for an appropriately named parent directory
    """
    rootdirs = []

    for i in range(3):
        dirname = os.path.basename(root)
        if dirname.startswith(parentdir_prefix):
            return {"version": dirname[len(parentdir_prefix):],
                    "full-revisionid": None,
                    "dirty": False, "error": None, "date": None}
        else:
            rootdirs.append(root)
            root = os.path.dirname(root)  # up a level

    if verbose:
        print("Tried directories %s but none started with prefix %s" %
              (str(rootdirs), parentdir_prefix))
    raise NotThisMethod("rootdir doesn't start with parentdir_prefix")


SHORT_VERSION_PY = """
# This file was generated by 'versioneer.py' (0.18) from
# revision-control system data, or from the parent directory name of an
# unpacked source archive. Distribution tarballs contain a pre-generated copy
# of this file.

import json

version_json = '''
%s
'''  # END VERSION_JSON


def get_versions():
    return json.loads(version_json)
"""


def versions_from_file(filename):
    """Try to determine the version from _version.py if present."""
    try:
        with open(filename) as f:
            contents = f.read()
    except EnvironmentError:
        raise NotThisMethod("unable to read _version.py")
    mo = re.search(r"version_json = '''\n(.*)'''  # END VERSION_JSON",
                   contents, re.M | re.S)
    if not mo:
        mo = re.search(r"version_json = '''\r\n(.*)'''  # END VERSION_JSON",
                       contents, re.M | re.S)
    if not mo:
        raise NotThisMethod("no version_json in _version.py")
    return json.loads(mo.group(1))


def write_to_version_file(filename, versions):
    """Write the given version number to the given _version.py file."""
    os.unlink(filename)
    contents = json.dumps(versions, sort_keys=True,
                          indent=1, separators=(",", ": "))
    with open(filename, "w") as f:
        f.write(SHORT_VERSION_PY % contents)

    print("set %s to '%s'" % (filename, versions["version"]))


def plus_or_dot(pieces):
    """Return a + if we don't already have one, else return a ."""
    if "+" in pieces.get("closest-tag", ""):
        return "."
    return "+"


def render_pep440(pieces):
    """Build up version string, with post-release "local version identifier".

    Our goal: TAG[+DISTANCE.gHEX[.dirty]] . Note that if you
    get a tagged build and then dirty it, you'll get TAG+0.gHEX.dirty

    Exceptions:
    1: no tags. git_describe was just HEX. 0+untagged.DISTANCE.gHEX[.dirty]
    """
    if pieces["closest-tag"]:
        rendered = pieces["closest-tag"]
        if pieces["distance"] or pieces["dirty"]:
            rendered += plus_or_dot(pieces)
            rendered += "%d.g%s" % (pieces["distance"], pieces["short"])
            if pieces["dirty"]:
                rendered += ".dirty"
    else:
        # exception #1
        rendered = "0+untagged.%d.g%s" % (pieces["distance"],
                                          pieces["short"])
        if pieces["dirty"]:
            rendered += ".dirty"
    return rendered


def render_pep440_pre(pieces):
    """TAG[.post.devDISTANCE] -- No -dirty.

    Exceptions:
    1: no tags. 0.post.devDISTANCE
    """
    if pieces["closest-tag"]:
        rendered = pieces["closest-tag"]
        if pieces["distance"]:
            rendered += ".post.dev%d" % pieces["distance"]
    else:
        # exception #1
        rendered = "0.post.dev%d" % pieces["distance"]
    return rendered


def render_pep440_post(pieces):
    """TAG[.postDISTANCE[.dev0]+gHEX] .

    The ".dev0" means dirty. Note that .dev0 sorts backwards
    (a dirty tree will appear "older" than the corresponding clean one),
    but you shouldn't be releasing software with -dirty anyways.

    Exceptions:
    1: no tags. 0.postDISTANCE[.dev0]
    """
    if pieces["closest-tag"]:
        rendered = pieces["closest-tag"]
        if pieces["distance"] or pieces["dirty"]:
            rendered += ".post%d" % pieces["distance"]
            if pieces["dirty"]:
                rendered += ".dev0"
            rendered += plus_or_dot(pieces)
            rendered += "g%s" % pieces["short"]
    else:
        # exception #1
        rendered = "0.post%d" % pieces["distance"]
        if pieces["dirty"]:
            rendered += ".dev0"
        rendered += "+g%s" % pieces["short"]
    return rendered


def render_pep440_old(pieces):
    """TAG[.postDISTANCE[.dev0]] .

    The ".dev0" means dirty.

    Eexceptions:
    1: no tags. 0.postDISTANCE[.dev0]
    """
    if pieces["closest-tag"]:
        rendered = pieces["closest-tag"]
        if pieces["distance"] or pieces["dirty"]:
            rendered += ".post%d" % pieces["distance"]
            if pieces["dirty"]:
                rendered += ".dev0"
    else:
        # exception #1
        rendered = "0.post%d" % pieces["distance"]
        if pieces["dirty"]:
            rendered += ".dev0"
    return rendered


def render_git_describe(pieces):
    """TAG[-DISTANCE-gHEX][-dirty].

    Like 'git describe --tags --dirty --always'.

    Exceptions:
    1: no tags. HEX[-dirty]  (note: no 'g' prefix)
    """
    if pieces["closest-tag"]:
        rendered = pieces["closest-tag"]
        if pieces["distance"]:
            rendered += "-%d-g%s" % (pieces["distance"], pieces["short"])
    else:
        # exception #1
        rendered = pieces["short"]
    if pieces["dirty"]:
        rendered += "-dirty"
    return rendered


def render_git_describe_long(pieces):
    """TAG-DISTANCE-gHEX[-dirty].

    Like 'git describe --tags --dirty --always -long'.
    The distance/hash is unconditional.

    Exceptions:
    1: no tags. HEX[-dirty]  (note: no 'g' prefix)
    """
    if pieces["closest-tag"]:
        rendered = pieces["closest-tag"]
        rendered += "-%d-g%s" % (pieces["distance"], pieces["short"])
    else:
        # exception #1
        rendered = pieces["short"]
    if pieces["dirty"]:
        rendered += "-dirty"
    return rendered


def render(pieces, style):
    """Render the given version pieces into the requested style."""
    if pieces["error"]:
        return {"version": "unknown",
                "full-revisionid": pieces.get("long"),
                "dirty": None,
                "error": pieces["error"],
                "date": None}

    if not style or style == "default":
        style = "pep440"  # the default

    if style == "pep440":
        rendered = render_pep440(pieces)
    elif style == "pep440-pre":
        rendered = render_pep440_pre(pieces)
    elif style == "pep440-post":
        rendered = render_pep440_post(pieces)
    elif style == "pep440-old":
        rendered = render_pep440_old(pieces)
    elif style == "git-describe":
        rendered = render_git_describe(pieces)
    elif style == "git-describe-long":
        rendered = render_git_describe_long(pieces)
    else:
        raise ValueError("unknown style '%s'" % style)

    return {"version": rendered, "full-revisionid": pieces["long"],
            "dirty": pieces["dirty"], "error": None,
            "date": pieces.get("date")}


class VersioneerBadRootError(Exception):
    """The project root directory is unknown or missing key files."""


def get_versions(verbose=False):
    """Get the project version from whatever source is available.

    Returns dict with two keys: 'version' and 'full'.
    """
    if "versioneer" in sys.modules:
        # see the discussion in cmdclass.py:get_cmdclass()
        del sys.modules["versioneer"]

    root = get_root()
    cfg = get_config_from_root(root)

    assert cfg.VCS is not None, "please set [versioneer]VCS= in setup.cfg"
    handlers = HANDLERS.get(cfg.VCS)
    assert handlers, "unrecognized VCS '%s'" % cfg.VCS
    verbose = verbose or cfg.verbose
    assert cfg.versionfile_source is not None, \
        "please set versioneer.versionfile_source"
    assert cfg.tag_prefix is not None, "please set versioneer.tag_prefix"

    versionfile_abs = os.path.join(root, cfg.versionfile_source)

    # extract version from first of: _version.py, VCS command (e.g. 'git
    # describe'), parentdir. This is meant to work for developers using a
    # source checkout, for users of a tarball created by 'setup.py sdist',
    # and for users of a tarball/zipball created by 'git archive' or github's
    # download-from-tag feature or the equivalent in other VCSes.

    get_keywords_f = handlers.get("get_keywords")
    from_keywords_f = handlers.get("keywords")
    if get_keywords_f and from_keywords_f:
        try:
            keywords = get_keywords_f(versionfile_abs)
            ver = from_keywords_f(keywords, cfg.tag_prefix, verbose)
            if verbose:
                print("got version from expanded keyword %s" % ver)
            return ver
        except NotThisMethod:
            pass

    try:
        ver = versions_from_file(versionfile_abs)
        if verbose:
            print("got version from file %s %s" % (versionfile_abs, ver))
        return ver
    except NotThisMethod:
        pass

    from_vcs_f = handlers.get("pieces_from_vcs")
    if from_vcs_f:
        try:
            pieces = from_vcs_f(cfg.tag_prefix, root, verbose)
            ver = render(pieces, cfg.style)
            if verbose:
                print("got version from VCS %s" % ver)
            return ver
        except NotThisMethod:
            pass

    try:
        if cfg.parentdir_prefix:
            ver = versions_from_parentdir(cfg.parentdir_prefix, root, verbose)
            if verbose:
                print("got version from parentdir %s" % ver)
            return ver
    except NotThisMethod:
        pass

    if verbose:
        print("unable to compute version")

    return {"version": "0+unknown", "full-revisionid": None,
            "dirty": None, "error": "unable to compute version",
            "date": None}


def get_version():
    """Get the short version string for this project."""
    return get_versions()["version"]


def get_cmdclass():
    """Get the custom setuptools/distutils subclasses used by Versioneer."""
    if "versioneer" in sys.modules:
        del sys.modules["versioneer"]
        # this fixes the "python setup.py develop" case (also 'install' and
        # 'easy_install .'), in which subdependencies of the main project are
        # built (using setup.py bdist_egg) in the same python process. Assume
        # a main project A and a dependency B, which use different versions
        # of Versioneer. A's setup.py imports A's Versioneer, leaving it in
        # sys.modules by the time B's setup.py is executed, causing B to run
        # with the wrong versioneer. Setuptools wraps the sub-dep builds in a
        # sandbox that restores sys.modules to it's pre-build state, so the
        # parent is protected against the child's "import versioneer". By
        # removing ourselves from sys.modules here, before the child build
        # happens, we protect the child from the parent's versioneer too.
        # Also see https://github.com/warner/python-versioneer/issues/52

    cmds = {}

    # we add "version" to both distutils and setuptools
    from distutils.core import Command

    class cmd_version(Command):
        description = "report generated version string"
        user_options = []
        boolean_options = []

        def initialize_options(self):
            pass

        def finalize_options(self):
            pass

        def run(self):
            vers = get_versions(verbose=True)
            print("Version: %s" % vers["version"])
            print(" full-revisionid: %s" % vers.get("full-revisionid"))
            print(" dirty: %s" % vers.get("dirty"))
            print(" date: %s" % vers.get("date"))
            if vers["error"]:
                print(" error: %s" % vers["error"])
    cmds["version"] = cmd_version

    # we override "build_py" in both distutils and setuptools
    #
    # most invocation pathways end up running build_py:
    #  distutils/build -> build_py
    #  distutils/install -> distutils/build ->..
    #  setuptools/bdist_wheel -> distutils/install ->..
    #  setuptools/bdist_egg -> distutils/install_lib -> build_py
    #  setuptools/install -> bdist_egg ->..
    #  setuptools/develop -> ?
    #  pip install:
    #   copies source tree to a tempdir before running egg_info/etc
    #   if .git isn't copied too, 'git describe' will fail
    #   then does setup.py bdist_wheel, or sometimes setup.py install
    #  setup.py egg_info -> ?

    # we override different "build_py" commands for both environments
    if "setuptools" in sys.modules:
        from setuptools.command.build_py import build_py as _build_py
    else:
        from distutils.command.build_py import build_py as _build_py

    class cmd_build_py(_build_py):
        def run(self):
            root = get_root()
            cfg = get_config_from_root(root)
            versions = get_versions()
            _build_py.run(self)
            # now locate _version.py in the new build/ directory and replace
            # it with an updated value
            if cfg.versionfile_build:
                target_versionfile = os.path.join(self.build_lib,
                                                  cfg.versionfile_build)
                print("UPDATING %s" % target_versionfile)
                write_to_version_file(target_versionfile, versions)
    cmds["build_py"] = cmd_build_py

    if "cx_Freeze" in sys.modules:  # cx_freeze enabled?
        from cx_Freeze.dist import build_exe as _build_exe
        # nczeczulin reports that py2exe won't like the pep440-style string
        # as FILEVERSION, but it can be used for PRODUCTVERSION, e.g.
        # setup(console=[{
        #   "version": versioneer.get_version().split("+", 1)[0], # FILEVERSION
        #   "product_version": versioneer.get_version(),
        #   ...

        class cmd_build_exe(_build_exe):
            def run(self):
                root = get_root()
                cfg = get_config_from_root(root)
                versions = get_versions()
                target_versionfile = cfg.versionfile_source
                print("UPDATING %s" % target_versionfile)
                write_to_version_file(target_versionfile, versions)

                _build_exe.run(self)
                os.unlink(target_versionfile)
                with open(cfg.versionfile_source, "w") as f:
                    LONG = LONG_VERSION_PY[cfg.VCS]
                    f.write(LONG %
                            {"DOLLAR": "$",
                             "STYLE": cfg.style,
                             "TAG_PREFIX": cfg.tag_prefix,
                             "PARENTDIR_PREFIX": cfg.parentdir_prefix,
                             "VERSIONFILE_SOURCE": cfg.versionfile_source,
                             })
        cmds["build_exe"] = cmd_build_exe
        del cmds["build_py"]

    if 'py2exe' in sys.modules:  # py2exe enabled?
        try:
            from py2exe.distutils_buildexe import py2exe as _py2exe  # py3
        except ImportError:
            from py2exe.build_exe import py2exe as _py2exe  # py2

        class cmd_py2exe(_py2exe):
            def run(self):
                root = get_root()
                cfg = get_config_from_root(root)
                versions = get_versions()
                target_versionfile = cfg.versionfile_source
                print("UPDATING %s" % target_versionfile)
                write_to_version_file(target_versionfile, versions)

                _py2exe.run(self)
                os.unlink(target_versionfile)
                with open(cfg.versionfile_source, "w") as f:
                    LONG = LONG_VERSION_PY[cfg.VCS]
                    f.write(LONG %
                            {"DOLLAR": "$",
                             "STYLE": cfg.style,
                             "TAG_PREFIX": cfg.tag_prefix,
                             "PARENTDIR_PREFIX": cfg.parentdir_prefix,
                             "VERSIONFILE_SOURCE": cfg.versionfile_source,
                             })
        cmds["py2exe"] = cmd_py2exe

    # we override different "sdist" commands for both environments
    if "setuptools" in sys.modules:
        from setuptools.command.sdist import sdist as _sdist
    else:
        from distutils.command.sdist import sdist as _sdist

    class cmd_sdist(_sdist):
        def run(self):
            versions = get_versions()
            self._versioneer_generated_versions = versions
            # unless we update this, the command will keep using the old
            # version
            self.distribution.metadata.version = versions["version"]
            return _sdist.run(self)

        def make_release_tree(self, base_dir, files):
            root = get_root()
            cfg = get_config_from_root(root)
            _sdist.make_release_tree(self, base_dir, files)
            # now locate _version.py in the new base_dir directory
            # (remembering that it may be a hardlink) and replace it with an
            # updated value
            target_versionfile = os.path.join(base_dir, cfg.versionfile_source)
            print("UPDATING %s" % target_versionfile)
            write_to_version_file(target_versionfile,
                                  self._versioneer_generated_versions)
    cmds["sdist"] = cmd_sdist

    return cmds


CONFIG_ERROR = """
setup.cfg is missing the necessary Versioneer configuration. You need
a section like:

 [versioneer]
 VCS = git
 style = pep440
 versionfile_source = src/myproject/_version.py
 versionfile_build = myproject/_version.py
 tag_prefix =
 parentdir_prefix = myproject-

You will also need to edit your setup.py to use the results:

 import versioneer
 setup(version=versioneer.get_version(),
       cmdclass=versioneer.get_cmdclass(), ...)

Please read the docstring in ./versioneer.py for configuration instructions,
edit setup.cfg, and re-run the installer or 'python versioneer.py setup'.
"""

SAMPLE_CONFIG = """
# See the docstring in versioneer.py for instructions. Note that you must
# re-run 'versioneer.py setup' after changing this section, and commit the
# resulting files.

[versioneer]
#VCS = git
#style = pep440
#versionfile_source =
#versionfile_build =
#tag_prefix =
#parentdir_prefix =

"""

INIT_PY_SNIPPET = """
from ._version import get_versions
__version__ = get_versions()['version']
del get_versions
"""


def do_setup():
    """Main VCS-independent setup function for installing Versioneer."""
    root = get_root()
    try:
        cfg = get_config_from_root(root)
    except (EnvironmentError, configparser.NoSectionError,
            configparser.NoOptionError) as e:
        if isinstance(e, (EnvironmentError, configparser.NoSectionError)):
            print("Adding sample versioneer config to setup.cfg",
                  file=sys.stderr)
            with open(os.path.join(root, "setup.cfg"), "a") as f:
                f.write(SAMPLE_CONFIG)
        print(CONFIG_ERROR, file=sys.stderr)
        return 1

    print(" creating %s" % cfg.versionfile_source)
    with open(cfg.versionfile_source, "w") as f:
        LONG = LONG_VERSION_PY[cfg.VCS]
        f.write(LONG % {"DOLLAR": "$",
                        "STYLE": cfg.style,
                        "TAG_PREFIX": cfg.tag_prefix,
                        "PARENTDIR_PREFIX": cfg.parentdir_prefix,
                        "VERSIONFILE_SOURCE": cfg.versionfile_source,
                        })

    ipy = os.path.join(os.path.dirname(cfg.versionfile_source),
                       "__init__.py")
    if os.path.exists(ipy):
        try:
            with open(ipy, "r") as f:
                old = f.read()
        except EnvironmentError:
            old = ""
        if INIT_PY_SNIPPET not in old:
            print(" appending to %s" % ipy)
            with open(ipy, "a") as f:
                f.write(INIT_PY_SNIPPET)
        else:
            print(" %s unmodified" % ipy)
    else:
        print(" %s doesn't exist, ok" % ipy)
        ipy = None

    # Make sure both the top-level "versioneer.py" and versionfile_source
    # (PKG/_version.py, used by runtime code) are in MANIFEST.in, so
    # they'll be copied into source distributions. Pip won't be able to
    # install the package without this.
    manifest_in = os.path.join(root, "MANIFEST.in")
    simple_includes = set()
    try:
        with open(manifest_in, "r") as f:
            for line in f:
                if line.startswith("include "):
                    for include in line.split()[1:]:
                        simple_includes.add(include)
    except EnvironmentError:
        pass
    # That doesn't cover everything MANIFEST.in can do
    # (http://docs.python.org/2/distutils/sourcedist.html#commands), so
    # it might give some false negatives. Appending redundant 'include'
    # lines is safe, though.
    if "versioneer.py" not in simple_includes:
        print(" appending 'versioneer.py' to MANIFEST.in")
        with open(manifest_in, "a") as f:
            f.write("include versioneer.py\n")
    else:
        print(" 'versioneer.py' already in MANIFEST.in")
    if cfg.versionfile_source not in simple_includes:
        print(" appending versionfile_source ('%s') to MANIFEST.in" %
              cfg.versionfile_source)
        with open(manifest_in, "a") as f:
            f.write("include %s\n" % cfg.versionfile_source)
    else:
        print(" versionfile_source already in MANIFEST.in")

    # Make VCS-specific changes. For git, this means creating/changing
    # .gitattributes to mark _version.py for export-subst keyword
    # substitution.
    do_vcs_install(manifest_in, cfg.versionfile_source, ipy)
    return 0


def scan_setup_py():
    """Validate the contents of setup.py against Versioneer's expectations."""
    found = set()
    setters = False
    errors = 0
    with open("setup.py", "r") as f:
        for line in f.readlines():
            if "import versioneer" in line:
                found.add("import")
            if "versioneer.get_cmdclass()" in line:
                found.add("cmdclass")
            if "versioneer.get_version()" in line:
                found.add("get_version")
            if "versioneer.VCS" in line:
                setters = True
            if "versioneer.versionfile_source" in line:
                setters = True
    if len(found) != 3:
        print("")
        print("Your setup.py appears to be missing some important items")
        print("(but I might be wrong). Please make sure it has something")
        print("roughly like the following:")
        print("")
        print(" import versioneer")
        print(" setup( version=versioneer.get_version(),")
        print("        cmdclass=versioneer.get_cmdclass(),  ...)")
        print("")
        errors += 1
    if setters:
        print("You should remove lines like 'versioneer.VCS = ' and")
        print("'versioneer.versionfile_source = ' . This configuration")
        print("now lives in setup.cfg, and should be removed from setup.py")
        print("")
        errors += 1
    return errors


if __name__ == "__main__":
    cmd = sys.argv[1]
    if cmd == "setup":
        errors = do_setup()
        errors += scan_setup_py()
        if errors:
            sys.exit(1)
```
