## Supporting information for "Markovian Weighted Ensemble Milestoning (M-WEM): Long-time Kinetics from Short Trajectories"

Dhiman Ray,[†] Sharon Emily Stone,[†] and Ioan Andricioaei[*,‡,‡]

†*Department of Chemistry, University of California Irvine, Irvine CA 92697*

‡*Department of Physics and Astronomy, University of California Irvine, Irvine CA 92697*

### Convergence of M-WEM simulation of Alanine Dipeptide

The convergence of the mean first passage times (MFPT) of the conformational transition of alanine dipeptide, computed from M-WEM simulation, is shown in Fig. S1. For all three trials the results converge in less than 30 iterations which incur a computational cost of about 40 ns, more than one order of magnitude less than the length of the conventional MD simulation. Although the results of the trial 2 and 3 are slightly lower than the error bars of conventional MD simulation, the discrepancy is less than 20% for most part of the M-WEM calculation.

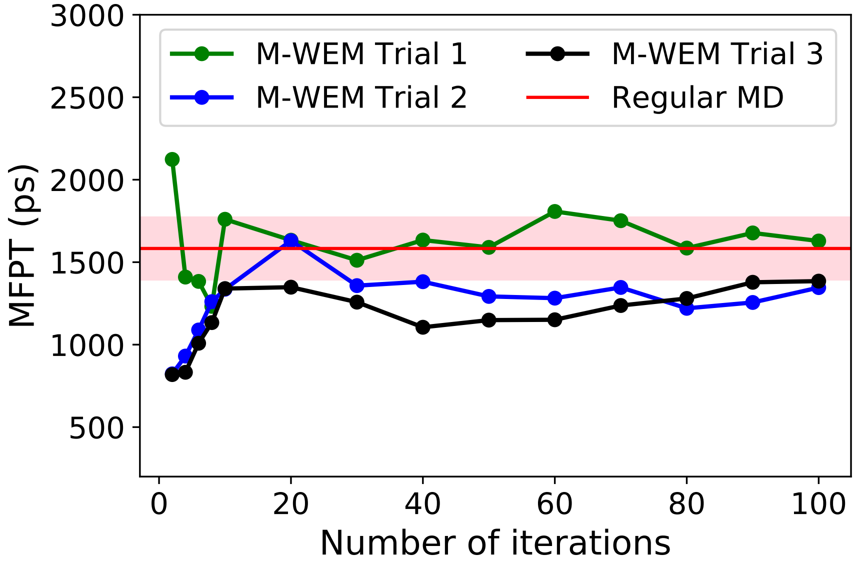


Figure S1: Convergence of the MFPT for the conformational transition in alanine dipeptide as a function of M-WEM iterations.

### Details of meta-eABF simulation of Trypsin-Benzamidine complex

To compare the free energy profile of ligand unbinding for the trypsin-benzamidine complex obtained from M-WEM simulation, we performed a long well tempered meta-eABF simulation to independently calculate the one dimensional free energy profile along the milestoning coordinate described in the main manuscript. In meta-eABF approach, two biasing techniques, metadynamics and extended system adaptive biasing force (eABF) are employed simultaneously for an accelerated sampling of the configurational space and a quicker convergence of the free energy. In well tempered meta-eABF (WTM-eABF) technique, the metadynamics is replaced by well tempered metadynamics which ensures a smooth convergence of the free energy landscape. Details of this method can be found elsewhere.[1–3] The equilibrated bound state structure was used as a starting point. Gaussian hills of width 0.6 Å were deposited every 2 ps along the distance between the Benzamidine ligand and the trypsin binding pocket. The initial heights of the hills were 0.2 kcal/mol which was gradually reduced during the course of the simulation using a bias factor of 6.0. The adaptive biasing force was applied on a fictitious particle coupled to the original reaction coordinate via stiff spring of force constant 1500 kcal mol $^{-1}$ Å $^{-2}$ with an oscillation period of 200 fs. External forces are applied only when at least 1000 samples are recorded in each bin of width 0.2 Å along the reaction coordinate. The WTM-eABF simulation is propagated for $\sim$ 550 ns. The convergence of the free energy profile is monitored by measuring the RMS deviation of the curve with respect to the final curve. The RMS deviation remained under 1 kcal/mol over the second half of the trajectory and went below 0.25 kcal/mol in last 50 ns (Fig. S2). The error bars of reported in the main manuscript is the 95 % confidence interval of 5 free energy profiles sampled at 8 ns interval from the last part of the meta-eABF trajectory.

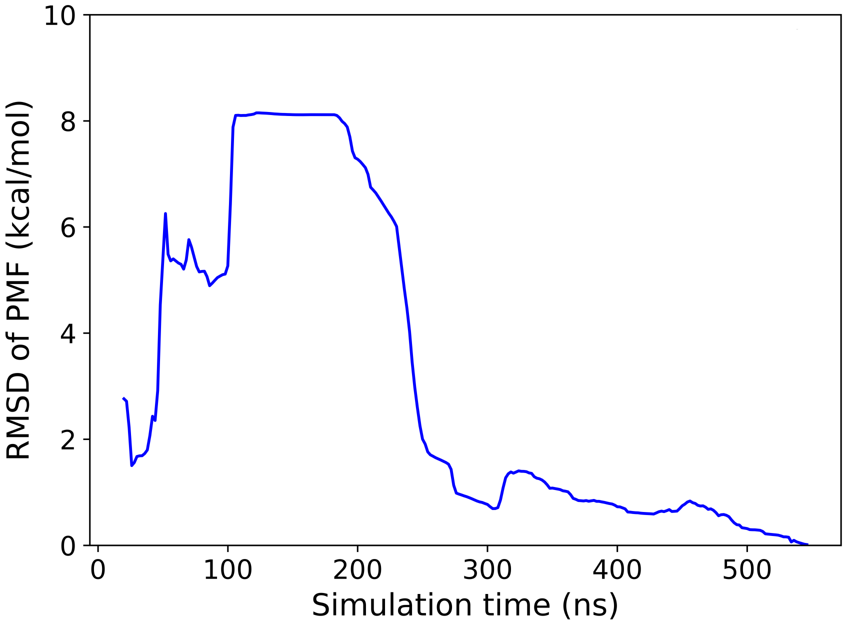


Figure S2: Convergence plot of WTM-eABF simulation of the trypsin-benzamidine complex.

### Calculation of diffusion coefficient of Benzamidine in water

Equilibrium molecular dynamics simulation was performed to estimate the diffusion coefficient of benzamidine in water for the $k_{\text{off}}$ calculation, as mentioned in the main text. A single molecule of the benzamidinium cation is solvated in TIP4P-Ew water using the CHARMM-GUI web server.[4] The water padding around the ligand is 20 Å in each direction. One $Cl^-$ ion is added to the system to balance the charge. This resulted in approximately 10000 atoms in the system. Molecular dynamics simulation on this system is performed using GROMACS 2018.6[5] software with identical simulation parameters as protein-ligand complex system. The energy of the solvated ligand was minimized using the steepest descent algorithm for 5000 steps. The minimized system was then equilibrated for 125 ps in NVT ensemble with 1 fs timestep followed by a 10 ns equilibration in NPT ensemble with 2 fs timestep. The temperature was kept constant at 298 K with Nose-Hoover thermostat[6,7] and the Pressure was controlled using a Parrinello-Rahman barostat.[8] A production run of 50 ns has been carried out from the final frame of the equilibration simulation in NPT ensemble.

Trajectory snapshots were saved at an interval of 100 ps for further analysis.

The mean squared displacement of the center of mass the benzamidinium ion was calculated from which the diffusion constant ($D$) is estimated as

$$D = \lim_{t\to\infty} \frac{\langle(\mathbf{r}(t) - \mathbf{r}(0))^2\rangle}{6t} \tag{1}$$

where $\mathbf{r}(t)$ is the vector of the center of mass of the ligand at time $t$. The estimated value of diffusion coefficient is 38 ± 1.1 $\text{Å}^2\text{ns}^{-1}$ or (0.38 ± 0.011) $\times 10^{-5}$ $\text{cm}^2\text{s}^{-1}$ (Fig. S3).

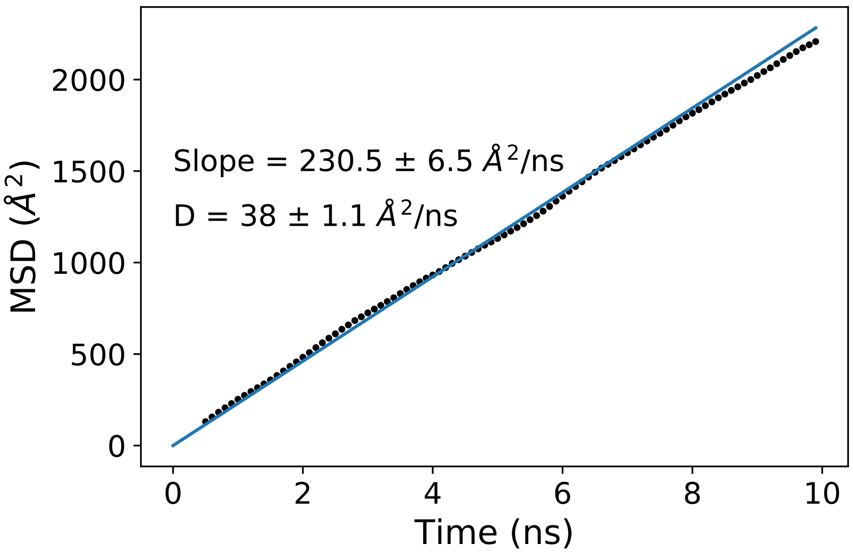


Figure S3: Mean square displacement of the center of mass benzamidinium ion, in TIP4P-ew water, as a function of time. The linear fit has been performed for the region between 2 ns and 10 ns.

### Discussion on appropriate placement of milestones based on velocity auto-correlation.

For calculating the kinetics and free energy profile from milestoning based methods the milestones need to be placed sufficiently far apart so the trajectories arriving at a given milestone looses the memory if the previously visited milestones.[9] To ensure this, the transition times between a pair of milestones should be larger than the decay time of the velocity auto-correlation function of the reaction coordinate.[10] But the velocity auto-correlation functions can have different values at different regions of the configurational space. So we computed the velocity auto-correlation function for each individual cell for the trypsin-benzamidine complex from 20 ns of regular MD simulation (Fig. S4). We also depict the average timescales ($\tau_\alpha$) of milestone to milestones transitions from M-WEM simulations for each cell computed as

$$\tau_\alpha = \frac{T_\alpha}{\sum_{\beta}^{\Lambda} N_{\alpha,\beta}}$$

where $N_{\alpha,\beta}$ is the number of transitions from cell $\alpha$ to cell $\beta$ recorded from a trajectory propagated for time $T_\alpha$ in cell $\alpha$. We observe that in our choice of milestone positions the velocity auto-correlations decays close to zero before completing a transition between two nearby milestones (Fig. S4).

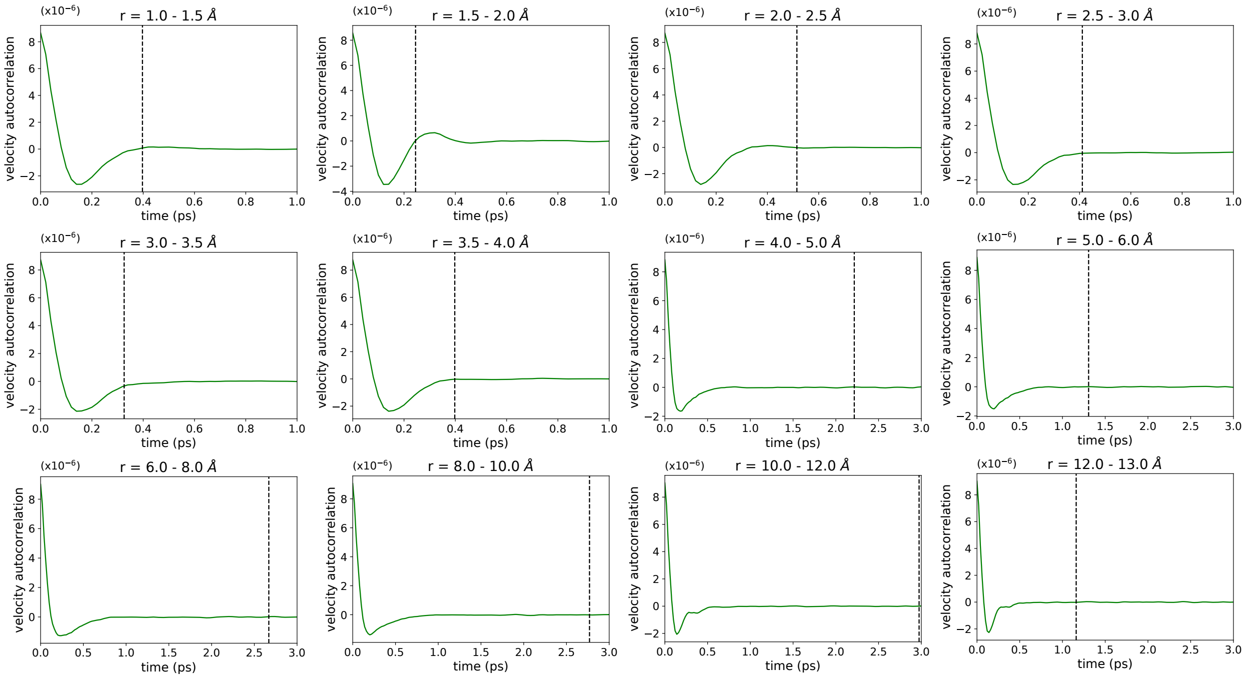


Figure S4: Velocity autocorrelation function of the reaction coordinate for each individual cell. The dashed vertical line indicate the average timescale of milestone to milestone transition in those cells in the M-WEM calculation.
